# A hydrolysis operon in *Lachnospiraceae* provides a genetic basis to understand castalagin metabolism by the human microbiome in a clinical study

**DOI:** 10.64898/2026.10.05.756759

**Authors:** Arianna Giurleo, Reilly Pidgeon, Emmanuel Gonzalez, Emma Thauvin, Lharbi Dridi, Corinne F. Maurice, André Marette, Bastien Castagner

**Affiliations:** Department of Pharmacology & Therapeutics, McGill University, 3655 Prom. Sir-William- Osler, Montreal, Quebec, H3G 1Y6, Canada; McGill Centre for Microbiome Research, Montreal, Quebec, Canada; Canadian Center for Computational Genomics, McGill University and Genome Quebec Innovation Center, Montreal, QC, Canada; Department of Human Genetics, McGill University, Montreal, QC, Canada; Institute of Nutrition and Functional Foods, Laval University, QC, Canada; Department of Medicine and IUCPQ, Laval University, Quebec, QC, Canada; Department of Microbiology & Immunology, McGill University, 3775 University Street, Montreal, Quebec, H3A 2B4, Canada

**Keywords:** Castalagin, polyphenol, ellagitannin, gut microbiota, metatranscriptomics, metagenomics, proteomics, uncultured, ellagic acid, castalin, urolithin, flavogallonic acid, human cohort

## Abstract

Gut microbes metabolize the ellagitannin castalagin to ellagic acid and urolithins, a process associated with host benefits. Despite clinical trials with the castalagin-rich camu camu extracts, the genes and taxa responsible for castalagin metabolism remain elusive. We discovered a castalagin hydrolase (*cah*) operon sparsely distributed in *Lachnospiraceae* using multi-omics. Analysis of fecal samples from a camu camu supplementation clinical trial in metabolic diseases demonstrates functional redundancy with prevalence of *cah* and other ellagic acid metabolism genes in different taxa, including uncultured species, which explains the prevalence of metabolites. We further identified flavogallonic acid as another prevalent metabolite of castalagin in human fecal samples. Fecal metabolite abundance correlated with patient response in the absence of pronounced microbiome compositional changes, suggesting a role for the metabolites in hepatic steatosis. Collectively, this work provides a genetic basis for castalagin metabolism and sets the stage to study the role of metabolism in clinical trials.

## Introduction

The human gastrointestinal tract is colonized by a highly diverse and genetically rich community of microorganisms, collectively forming the gut microbiota^1,2^. These microorganisms play key roles in maintaining gut barrier integrity, colonization resistance, host immune function, and in metabolizing dietary components^3^. This ecological niche exceeds the genetic repertoire of the human genome by approximately 100-fold^4^. Despite this diversity, nearly half of predicted protein-coding genes remain uncharacterized^4,5^. Gut bacteria encode extensive metabolic enzymes, catalyzing hydrolysis, conjugation, reduction, and cleavage reactions, enabling the biotransformation of a wide range of endogenous^6,7^ and exogenous compounds^8–10^. Through these enzymatic reactions, gut microbes facilitate absorption of host-inaccessible nutrients and produce immunomodulatory molecules, driving both host-microbe and microbe-microbe interactions that have implications in host health^3,11^.

Dietary ellagitannins, a class of polyphenols belonging to the family of hydrolyzable tannins, are non-bioavailable plant secondary metabolites consumed through various products such as fruits, vegetables, nuts and teas^12,13^. Due to their size and polarity, ellagitannins like punicalagin, castalagin, and vescalagin reach the colon relatively intact, relying on the enzymatic abilities of gut microbes for their breakdown^14^. Castalagin, a dietary *C*-glucosidic ellagitannin found in the camu-camu (CC, *Myrciaria dubia*) berry^15^, walnuts^16,17^ and in oak barrel aged beverages^18^, has been demonstrated to reduce hepatic steatosis in overweight adults with hypertriglyceridemia (NCT04130321) and prevent obesity in diet-induced mouse studies^19,20^. Additionally, CC supplementation has been associated with beneficial effects in cancer, circumventing anti-PD-1 resistance and potentiating immunotherapy^15,21,22^. Thus, castalagin-rich extracts have also been used in clinical trials in conjunction with cancer immunotherapy (NCT05303493 and NCT06049576). This polyphenol has also been linked with the inhibition of pro- inflammatory cytokines IL-8 and NF-kB^23,24^.

Ellagitannins are extensively broken down to bioavailable urolithins by gut bacteria through cross-feeding^25,26^. Polyphenol metabolites have been associated with various beneficial health effects such as enhancing mitophagy, antioxidant, anti-inflammatory, and cardioprotective effects and have been detected in human urine, plasma and feces at varying concentrations among individuals^14,27,28^. These various metabolite profiles reflect the interindividual variability in polyphenol metabolism based on gut microbial composition^25,29–31^. In the gut, castalagin is hydrolyzed to produce ellagic acid (EA) and castalin^32^ [**Fig. 1a**]. The decarboxylation of EA and subsequent dehydroxylation reactions of urolithins, are attributed to the recently-described *eam* (EA metabolism)^33^, *ucd* (uroC dehydroxylase)^34^, and *uxd* (urolithin 10-position dehydroxylase)^26^ gene clusters [**Supplementary Fig. 1**] in *Gordonibacter* and *Enterocloster* spp., while the fate of castalin is unknown.

**Figure 1.**
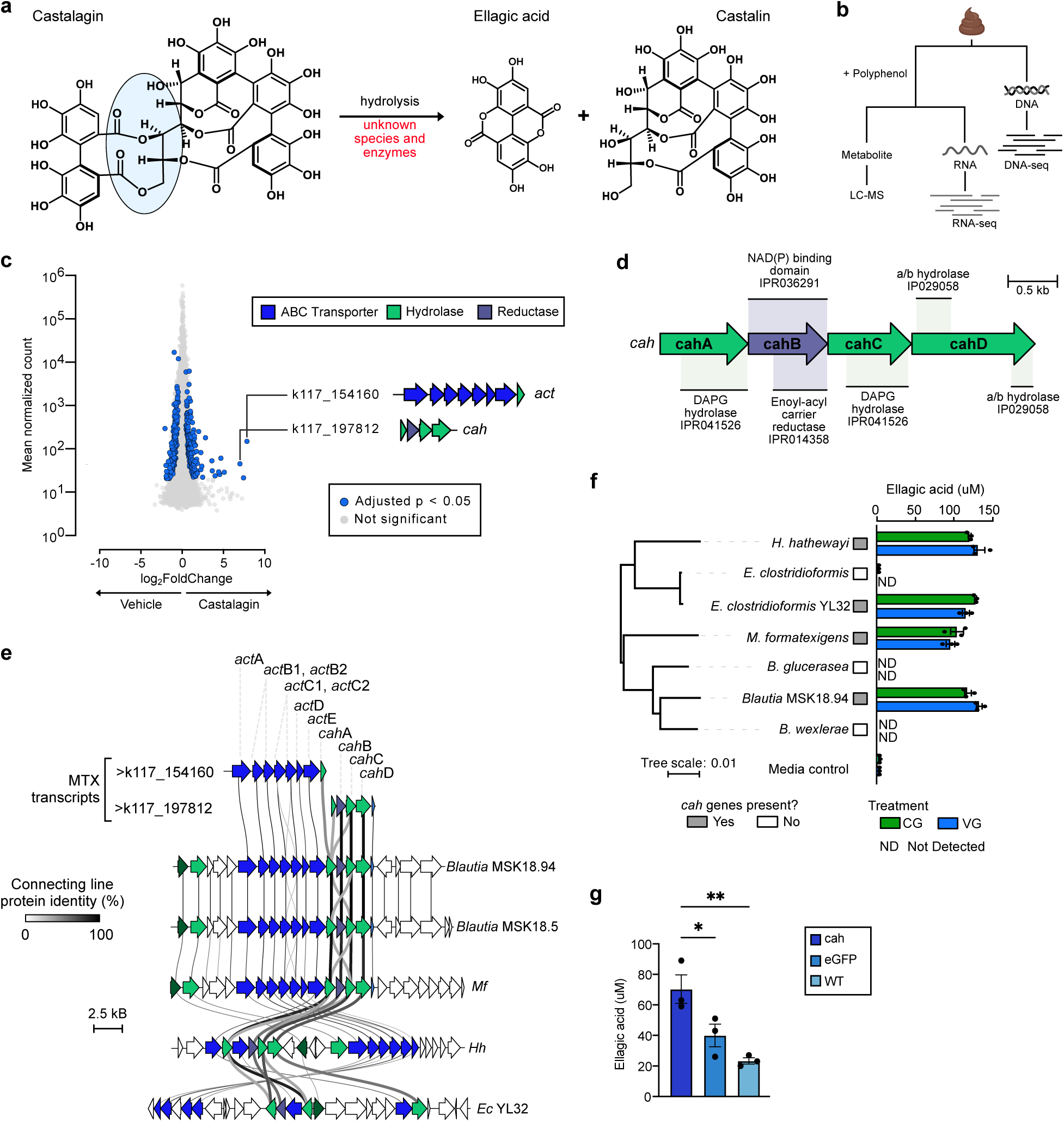
A hydrolase gene cluster induced by castalagin correlates with production of ellagic acid. **a** Castalagin hydrolysis scheme to ellagic acid and castalin by unknown enzymes. **b** Schematic representing the multi-omics workflow starting from a complex microbial community from a healthy human donor fecal sample. **c** MA plot of transcripts in MO67 fecal slurry treated with castalagin (n = 4 biological replicates). Dots in blue represent significantly upregulated transcripts (Benjamini-Hochberg (BH) FDR < 0.05), grey dots include non-significant transcripts and those without an adjusted p-value. Transcripts are indicated by transcript identity label, and individual genes are coloured based on annotated function. **d** Domain annotations of individual Cah proteins in *Blautia* MSK.18.94 based on InterPro accessions. **e** Comparative genomics between upregulated transcripts and the top BLASTx search results from *Blautia* MSK.18.94, *Blautia luti* MSK.18.5 (*Blautia* MSK.18.5), *M. formatexigens* (*Mf*), *H. hathewayi* (*Hh*) and *Enterocloster clostridioformis* YL32 (Ec YL32). Alignments were performed using clinker with GenBank accessions provided in **Supplementary Table 7**. Connecting lines represent protein percent identity and only sequences with >30% identity are connected. Individual genes are coloured according to functional annotations as indicated in panel **d**. **f** Relative concentration of ellagic acid (μM) measured in liquid cultures of bacterial isolates incubated with castalagin (green) or vescalagin (blue) (n=3 biological replicates, error bars represent mean ± SEM). *Cah* gene cluster harbouring species are indicated by grey boxes. **g** Quantification of ellagic acid concentration (μM) in cell lysates of *C. sporogenes* expressing Cah from *Blautia* MSK.18.94 or eGFP compared to the wild-type (WT) control (n=3 biological replicates) incubated with castalagin (300 μM) at 37 °C after 48 hours. Error bars represent mean ± SEM, analyses were performed using One-way ANOVA with Dunnett’s multiple comparison correction, *P ≤ 0.05, **P ≤ 0.01.

Despite the associations between castalagin, and its metabolites, with host health, the microorganisms and enzymes responsible for the hydrolysis of castalagin remain elusive. Versatile tannin acyl hydrolases (tannases, EC 3.1.1.20) have been identified in *Lactiplantibacillus plantarum* but are specific to galloylated ellagitannins and are unable to hydrolyse *C*-glucosidic ellagitannins like punicalagin, castalagin and vescalagin, to EA^35,36^. Here, we report the discovery of genes and enzymes in gut bacteria involved in the metabolism of castalagin. Using multi-omics and comparative genomics approaches, we discover the *cah* operon in *Lachnospiraceae* genomes that enables the conversion of castalagin to EA and castalin. By interrogating a human supplementation study, we find that, despite its sparse taxonomic distribution and strain-specificity, the *cah* operon is prevalent in human gut metagenomes. We additionally identify a subset of *cah*-encoding species that can metabolize castalin into flavogallonic acid, a metabolite previously undescribed in humans. The abundance of fecal metabolites correlated with patient response in the absence of drastic microbial composition changes, suggesting their potential role in metabolic diseases. Finally, using a genome mining approach, we discover and functionally validate EA and urolithin metabolism genes in uncharacterized and uncultured metagenomes. Overall, our study identifies novel microbial contributors to multiple steps in the ellagitannin metabolism pathway by the human gut microbiome and explains the prevalence of ellagitannin metabolism in human populations.

## Results

### Castalagin induces a hydrolase gene cluster in *Lachnospiraceae*

Considering the absence of known castalagin-metabolizing species [**Fig. 1a**] and the vast distribution of gut-microbiome derived hydrolases and esterases^37^, we sought to investigate ellagitannin metabolism in complex fecal communities by leveraging metatranscriptomics, hypothesizing inducible metabolism genes [**Fig. 1b**]. To identify an appropriate fecal community and timepoint for RNA extraction, we performed *ex vivo* metabolism assays on three fecal samples with castalagin and monitored metabolism over time by liquid chromatography coupled with mass spectrometry (LC-MS). Due to the propensity of ellagitannins to complex with proteins in culture media^38,39^, castalagin metabolism was inferred from the detection of its downstream metabolites: EA and urolithins. Castalagin metabolism was most rapid in sample MO67, as urolithin A (uroA) was detected after only 4 hours [**Supplementary Fig. 2a**]. Therefore, we selected both the MO67 fecal sample and the 4-hour timepoint for metatranscriptomics analysis, as it captured peak metabolism occurring in the sample, while limiting shifts in the microbial community.

To capture differentially-expressed transcripts involved in the initial step of castalagin hydrolysis, fecal slurries of MO67 were supplemented in quadruplicate with castalagin, its immediate downstream metabolites castalin or EA, or a vehicle control prior to RNA isolation. Consistent with our initial observation, castalagin-derived metabolites, including EA, urolithin M5 (uroM5), urolithin M6 (uroM6), and urolithin C (uroC), were readily detected after 4 hours. A similar profile was observed in EA-treated cultures; however, no downstream metabolites were detected in samples treated with castalin or vehicle [**Supplementary Fig. 2b**]. We additionally isolated total DNA from MO67 to determine species relative abundance and gene content [**Fig. 1b**]. Metagenomics analysis of MO67 revealed a sample primarily composed of Bacillota (previously Firmicutes), with *Lachnospiraceae* representing the dominant family within the phylum [**Supplementary Fig. 2c**]. To identify putative castalagin metabolism genes, we co-assembled reads from all four treatment groups into a common reference and quantified transcript abundance separately for each sample against this assembly. We then analyzed all 16 samples jointly in DESeq2 to assess treatment effects while accounting for differences between matched incubation blocks^40^. We identified 357, 1125 and 824 significantly differentially expressed transcripts (Benjamini-Hochberg FDR < 0.05) in response to castalagin, castalin and EA versus vehicle, respectively, demonstrating that these molecules induce distinct changes in gene expression [**Supplementary Fig. 2d, e**]. Functional annotations of upregulated genes were predominantly associated with primary active transport, efflux, and hydrolysis [**Fig. 1c**].

We identified 90 shared significant transcripts between the castalagin and castalin conditions, 143 shared between the castalagin and EA condition and 9 shared between the castalin and EA groups. We centered our efforts on the two mostly differentially expressed transcripts in the castalagin condition, which were also significantly differentially abundant in the castalin group [**Fig. 1c, Supplementary Fig. 2d**]. Transcript k117_197812, which we termed the <u>ca</u>stalagin <u>h</u>ydrolase (*cah*) gene cluster, was significantly differentially expressed (log_2_FC = 7.013 and adjusted p = 1.70 × 10^−7^) in the castalagin condition [**Fig. 1c**]. This gene cluster contained genes encoding two 2,4- diacetylphloroglucinol (DAPG) hydrolase family proteins (EC 3.7.1.24) (*cahA* and *cahC*), a short chain dehydrogenase/reductase (SDR) family protein (EC 1.1.1) (*cahB*), and an alpha/beta hydrolase family protein (*cahD*) [**Fig. 1d**]. Interestingly, the most highly differentially-expressed transcript k117_154160 (log_2_FC = 7.852 and adjusted p = 1.17 × 10^−2^) was comprised of ABC transporter subunit genes that we termed the active castalagin transport (*act*) cluster, suggesting a link between active transport and castalagin metabolism. By integrating contigs derived from the metagenomics assembly of MO67 with the identified RNA-seq transcripts, we observed that both transcripts k117_197812 (*cah*) and k117_154160 (*act*) co-localized within the same genomic region, sharing portions of the *cahA* gene. The upregulation of *cah* and *act* by both castalagin and castalin suggests that both compounds are recognized by the same transcriptional machinery, despite castalin being a metabolite of castalagin.

### Castalagin metabolism correlates with *cah* presence in gut bacteria

Based on the induction of *cah* and *act,* and the predicted hydrolase function of *cah*- encoded proteins (CahA, CahC, and CahD), we hypothesized that species harbouring the *cah* gene cluster can hydrolyze castalagin. A sequence-based homology search of the k117_197812 (*cah*) and k117_154160 (*act*) transcripts against the NCBI refseq_genomes database identified *Blautia* MSK.18.94 and MSK.18.5 (Blautia_A sp905206915 in the Genome Taxonomy Database R232, **Supplementary Discussion 1**) as the closest culturable species, with 85.60% and 86.56% nucleotide identity to *cah* and *act*, respectively. Additionally, we identified *cah* gene homologues in other *Lachnospiraceae*, including *Marvinbryantia formatexigens* (*Mf*), *Hungatella hathewayi* (*Hh*) and *Enterocloster clostridioformis* YL32 (*Ec* YL32) [**Fig. 1e**]. Each of these taxa harbour protein-coding sequences with high identity and similarity (57-95%) to MSK.18.94 Cah proteins [**Supplementary Fig. 3a, b**], and encode ABC transport subunits that are homologous to Act proteins [**Fig. 1e**].

To functionally validate the metabolic capabilities of these bacteria, we obtained available strains of *Blautia* MSK.18.94, *Mf* (DSM 14469), *Hh* (DSM 13479), *Ec* YL32 (DSM 26114), as well as the type strains of related *Blautia glucerasea* (*Bg*, DSM 22028), *Blautia wexlerae* (*Bw*, DSM 19850) and *E. clostridioformis* (*Ec*, DSM 933), which lack the *cah* gene cluster. After incubation with castalagin, EA detection was higher in cultures of *Blautia* MSK.18.94, *Hh*, *Mf* and *Ec* YL32 compared to sterile media and vehicle-treated controls. In contrast, no EA was detected in cultures of *Bg, Bw* or *Ec* compared to baseline detection in media alone [**Fig. 1f**]. Similar results were obtained from cultures treated with vescalagin, a C-1 epimer of castalagin [**Fig. 1f, Supplementary Fig. 4a, b]**. These data demonstrate that *cah* presence correlates with EA production and that the stereochemistry of the C-1 position of castalagin and vescalagin does not impact its metabolism by *cah*-harbouring gut bacteria.

Next, we wondered whether castalagin- and vescalagin-metabolizing species could metabolize other plant-derived hydrolysable tannins. We performed metabolism assays with punicalagin, a *C*-glycosidic ellagitannin found in pomegranates, and corilagin, an ellagitannin with a glucopyranose core [**Supplementary Fig. 5a**]. No EA was detected in the punicalagin-treated cultures. Conversely, EA was detected at low levels in corilagin- treated cultures of *Hh* and *Ec* YL32, suggesting additional metabolic capability in these species compared to the other identified castalagin metabolizers [**Supplementary Fig. 5b**]. To validate the function of the *Blautia* MSK.18.94 *cah*, we expressed the CahABCD proteins in *Clostridium sporogenes* (*Cs*), a genetically tractable heterologous host^41^. Primary assays were performed in whole cells; however, no conversion of castalagin to EA was observed [**Supplementary Fig. 6**]. This further suggests that active transport mediated by the *act* gene cluster is required for the biotransformation of castalagin. Considering this limitation, we performed activity assays in induced cell lysates. Significantly more EA was detected in castalagin-treated lysates of *Cs* expressing CahABCD compared to the wild type (WT) or enhanced green fluorescent protein (eGFP) -expressing controls [**Fig. 1g**].

### A subset of *cah*-encoding bacteria convert castalin to flavogallonic acid

Since *cah* genes were also induced in castalin-treated conditions [**Supplementary Fig. 2d**], we next asked whether *cah* can act on castalin. A previous report of an intermediate metabolite corresponding to flavogallonic acid (FGA) produced by the microbiota of post weaning piglets suggests it may be derived from castalagin^42^ [**Fig. 2a**]. Metabolism assays with castalagin and castalin resulted in the emergence of a previously unobserved metabolite peak with a m/z of 469 ([M-H]^−^) in only a subset of the species: *H. hathewayi* and *E. clostridioformis* YL32 [**Fig. 2b**]. This was validated to be flavogallonic acid (FGA) using a commercial standard. Metabolism of these two substrates by *Hh* overtime demonstrates a marked increase in FGA with the decrease of castalin, with no effect imparted from the production of EA [**Fig. 2c**]. To our knowledge, this is the first report of the conversion of castalin to FGA in bacterial isolates.

**Figure 2.**
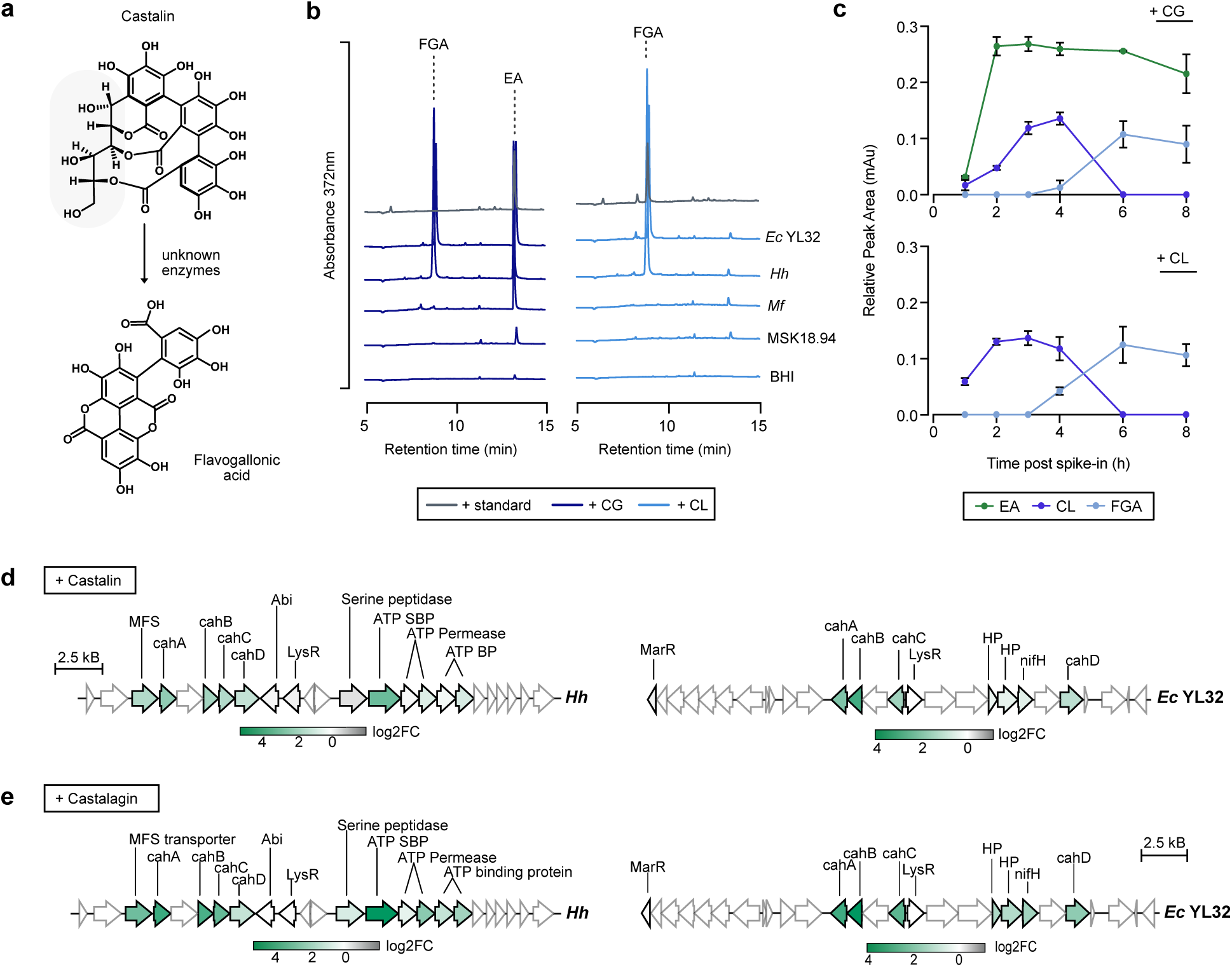
Flavogallonic acid is a product of castalin metabolism. **a** Conversion of castalin to flavogallonic acid by unknown enzymes. **b** Aligned chromatograms (λ = 372 nm) of castalagin (200 μM, dark blue lines) or castalin (200 μM, light blue lines) treated cultures of *E. clostridioformis* YL32 (*Ec* YL32), *H. hathewayi* (*Hh*), *M. formatexigens* (*Mf*), or *Blautia* MSK.18.94 (MSK18.94) in BHI media after 48 hours. Retention times of EA and FGA standards are shown by grey lines. Chromatograms are n = 1 representative of n = 3 biological replicates **c** Quantification of ellagic acid (EA), castalin, and flavogallonic acid (FGA), during metabolism of castalagin or castalin by *H. hathewayi* at 1, 2, 3, 4, 6, and 8 hours post treatment (mean ± SEM). **d, e** Genomic context of proteins differentially expressed in *Hh* and *Ec* YL32 cultures treated with (**d**) castalagin or (**e**) castalin based on log_2_(fold change) relative to vehicle. Proteins bordered in grey were not detected in the proteomic analysis. HP: hypothetical protein, LysR: transcriptional regulator, MarR: multiple antibiotic resistance regulator, MFS: major facilitator superfamily.

To confirm that *cah* encoded proteins were expressed in *Hh* and *Ec* YL32 upon castalin induction, we performed untargeted proteomics on castalagin-, castalin-, or vehicle- treated isolates. Consistent with the metatranscriptomics data, all four proteins were differentially expressed in castalagin and castalin treatment groups in both isolates [**Fig. 2d, e, Supplementary Fig. 7a, b**]. Interestingly, significantly differentially expressed proteins in the vehicle condition were associated with ribosomal functions [**Supplementary Fig. 7c**] suggesting a possible shift in translational activity in the presence of these substrates^43^.

### *cah* is strain specific and sparsely distributed in *Lachnospiraceae*

To investigate the diversity and taxonomic distribution of Cah*-*harbouring species, we queried individual Cah proteins against genomes belonging to the *Blautia*, *Marvinbryantia*, *Hungatella*, and *Enterocloster* genera. Within these genera, Cah and Act were limited to a subset of species [**Fig. 3a**, **Supplementary Fig. 8a**]. At the strain level, we found that Cah proteins in these genera were widespread in the genomes assigned to *Hh* and *Mf* but less prevalent in the other species. In *Hh*, all but two available genomes have at least three Cah proteins and in *Mf*, Cah proteins were detected in all genomes (> 60% ID), suggesting functional conservation in these species [**Fig. 3a**]. Similarly, Act proteins with >60% ID are also conserved in *Hh*, *Mf* and MSK.18.94 but sparsely distributed in *Ec* YL32 [**Supplementary Fig. 8b**]. These comparative genomic analyses suggest that Cah, and potentially castalagin metabolism, vary in a strain-specific manner. To further investigate the taxonomic distribution of Cah, we queried genomes across o Christensenellales and o Lachnospirales for homologues of functionally validated Cah proteins. This analysis revealed that Cah homologues were rare, being detected in only five species from Christensenellales and 21 species from Lachnospirales. Additionally, these homologues were sparsely distributed across 15 genera [**Supplementary Fig. 9a**] with only 46% (12/26) encoding the full Cah cluster [**Supplementary Fig. 9b**]. These results further indicate that Cah has a sparse, strain specific distribution and does not cluster among closely related taxa.

**Figure 3.**
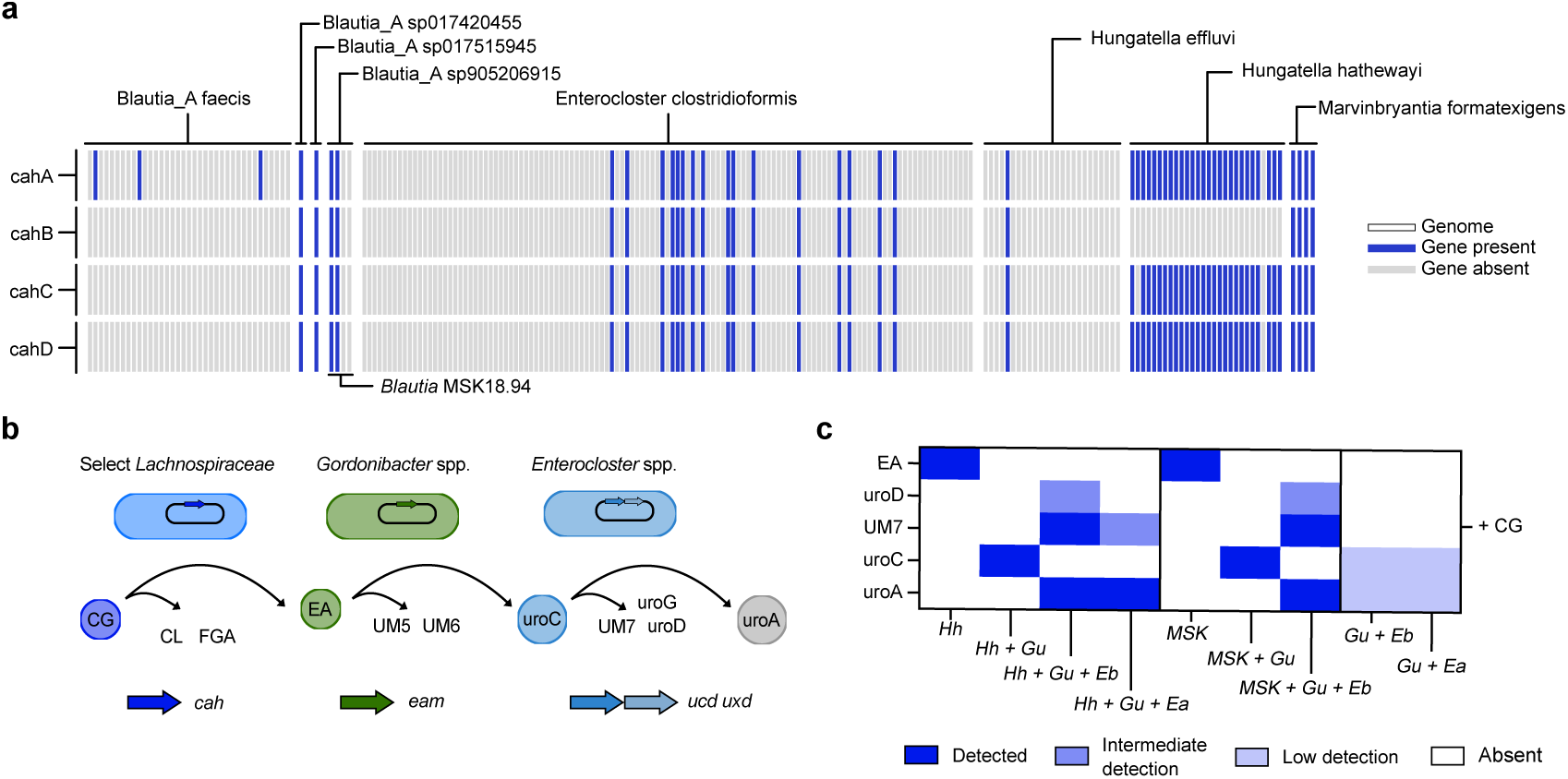
Strain-specific 3-member bacterial communities dictate metabolite production. **a** Heat map of individual *cah* genes present in *Blautia*, *Enterocloster*, *Hungatella* and *Marvinbryantia* genomes from the NCBI based on BLASTx matches to reference genes from *Blautia* MSK.18.94 with ≥ 60% identity. Species with at least one match are included. Bars represent an individual genome within the indicated species; blue bars represent a match for the indicated gene in that genome. **b** Schematic demonstrating the cross-feeding of castalagin metabolites. Select *cah*-harbouring *Lachnospiraceae* convert castalagin to ellagic acid (EA) and castalin (CL). *eam*-harbouring *Gordonibacter* spp. and *Ellagibacter* spp. metabolize ellagic acid (EA) to urolithins. Select intermediate urolithins like urolithin C (uroC) can be further converted to terminal metabolite urolithin A (uroA) by *Enterocloster* spp. via the *uxd* and *ucd* operons. **c** Heatmap of metabolites detected in castalagin (CG)-treated cultures of *H. hathewayi* (Hh), Hh + *G. urolithinfaciens* (Gu), Hh + Gu + *E. asparagiformis* (Ea), Hh + Gu + *E. bolteae* (Eb), *Blautia* MSK18.94 (MSK), MSK + Gu, MSK + Gu + Eb, Gu + Eb and Gu + Ea.

In our metatranscriptomics dataset, the *cah* gene cluster was assembled into a single transcript; however, due to potential assembly errors, the *cahA* gene was split in two. To validate that all *cah* genes were part of an inducible operon, we used RT-PCR with primers flanking the *cah* region of *Blautia* MSK.18.94, which was treated with castalagin or vehicle [**Supplementary Fig. 10a]**. A single polycistronic RNA of the expected size, which included genes from *cahA* to a gene following *cahD*, was detected only in castalagin-treated cultures [**Supplementary Fig. 10b**]. The conserved arrangement of the *cah* gene cluster in *Mf* suggests a similar operon structure in this species. In contrast, the *cah* genes in *Hh* and *Ec* YL32 are interspersed with other loci [**Fig. 1e**]. Interestingly, some of these interspersed genes are mobile genetic elements (MGEs) [**Supplementary Fig. 10c**]. These differences support that the *cah* cluster may not be universally present as an operon in all *cah* harbouring species^44,45^.

### Minimal 3-member bacterial communities dictate uroA production

EA-metabolizing bacteria produce urolithin intermediates that can subsequently be utilized by other bacterial species for further degradation^25,26,33^ [**Fig. 3b, Supplementary Fig. 11a**]. We investigated whether castalagin metabolism could be directed by manipulating the composition of metabolizing species and their corresponding genetic capabilities. To evaluate the potential for cross-feeding interactions during castalagin metabolism, we established co-cultures of a three species community and assessed metabolite production profiles. We incubated *cah*-encoding *Hh* with known metabolizers of EA, urolithin M6 (uroM6), and urolithin C (uroC): *G. urolithinfaciens* (*Gu*), *E. asparagiformis* (*Ea*) and *E. bolteae* (*Eb*), respectively. Co-incubation of *Gu* and *Eb* or *Ea* with castalagin produced minimal amounts of uroC and uroA due to some spontaneous hydrolysis of castalagin in media. Co-incubation of *Hh*, *Gu*, and *Eb* mainly produced the intermediate urolithin M7 (uroM7) and terminal uroA, whereas replacing *Eb* by *Ea* that encodes the urolithin 10 dehydroxylase (*uxd*) yielded mostly uroA [**Fig. 3c**, **Supplementary Fig. 11b**]. Omission of the *Enterocloster* species from the co-culture led to accumulation of the UroC metabolite. Similar results were obtained with the co- incubation of *Blautia* MSK.18.94, *Gu*, and *Eb* [**Fig. 3c, Supplementary Fig. 11c**]. These results suggest that the presence or absence of a single species can direct metabolic flux. Further, the nature of the species present in the community dictates the metabolite output, shaping the availability of substrates for downstream conversion. Together, these findings define a minimal community capable of converting castalagin to terminal metabolite uroA. This simplified system, however, does not capture the taxonomic and genetic complexity present in the human gut microbiome.

### Castalagin metabolism is prevalent in a CC supplementation clinical trial

Recognizing the numerous reports of beneficial effects upon ellagitannin supplementation, we further examined the prevalence of castalagin metabolism in human microbiome samples. We analyzed fecal samples from a patient cohort of overweight adults with hypertriglyceridemia supplemented with castalagin-rich CC extract (NCT04130321)^19^, to assess metabolite production, bacterial species, and gene presence. Fecal samples from 33 patients, collected pre- and post-CC, were processed for targeted metabolomics of castalagin metabolites EA, urolithin M5 (uroM5), uroM6, uroM7, uroC, urolithin D (uroD), urolithin B (uroB), uroA, isourolithin A (isoA) and FGA using LC-MS/MS. Normalized metabolite abundances revealed interindividual variability in metabolite production profiles within the cohort [**Supplementary Fig. 12a**]. Despite this heterogeneity, CC-supplementation resulted in the significant increase (P < 0.01) in castalagin-derived metabolites EA, uroM5, uroC, uroD, uroA, and FGA which were prevalent among patients [**Fig. 4a, b**, **Supplementary Fig. 12b**]. This interindividual variability and metabolite increase is consistent with observations from ellagitannin-rich supplementation studies, whereby production of metabolites, primarily uroA have been observed^46–48^. In addition to uroA, we observed high relative proportions of early metabolites uroC and uroD post CC [**Fig. 4a**]. Due to the significant increase in metabolites, a correlation analysis was performed to investigate the relationship between total metabolite production and precent (%) change in hepatic fat fraction (HFF). A significant negative correlation was observed between change in total metabolite production and change in response (P = 0.01) such that a decrease in %HFF correlates with an increase in metabolite abundance [**Fig. 4c**]. Further, we investigated individual metabolite correlation with response and observed negative correlations (*ρ* > −0.5) with EA, uroM5 and uroD production and change in HFF, however these correlations were not found to be significant after multiple comparison correction [**Fig. 4d, Supplementary Fig. 12c**]. Additionally, in the top and bottom quartiles of patient response there is an increase in EA (P = 0.02), uroM5 and uroD (P = 0.06) abundance in the top quartile of responders compared to the bottom quartile [**Fig. 4e**]. Due to the small number of subjects in this cohort, the response cannot be attributed to any one metabolite, warranting this analysis to be conducted in a larger scale trial.

**Figure 4.**
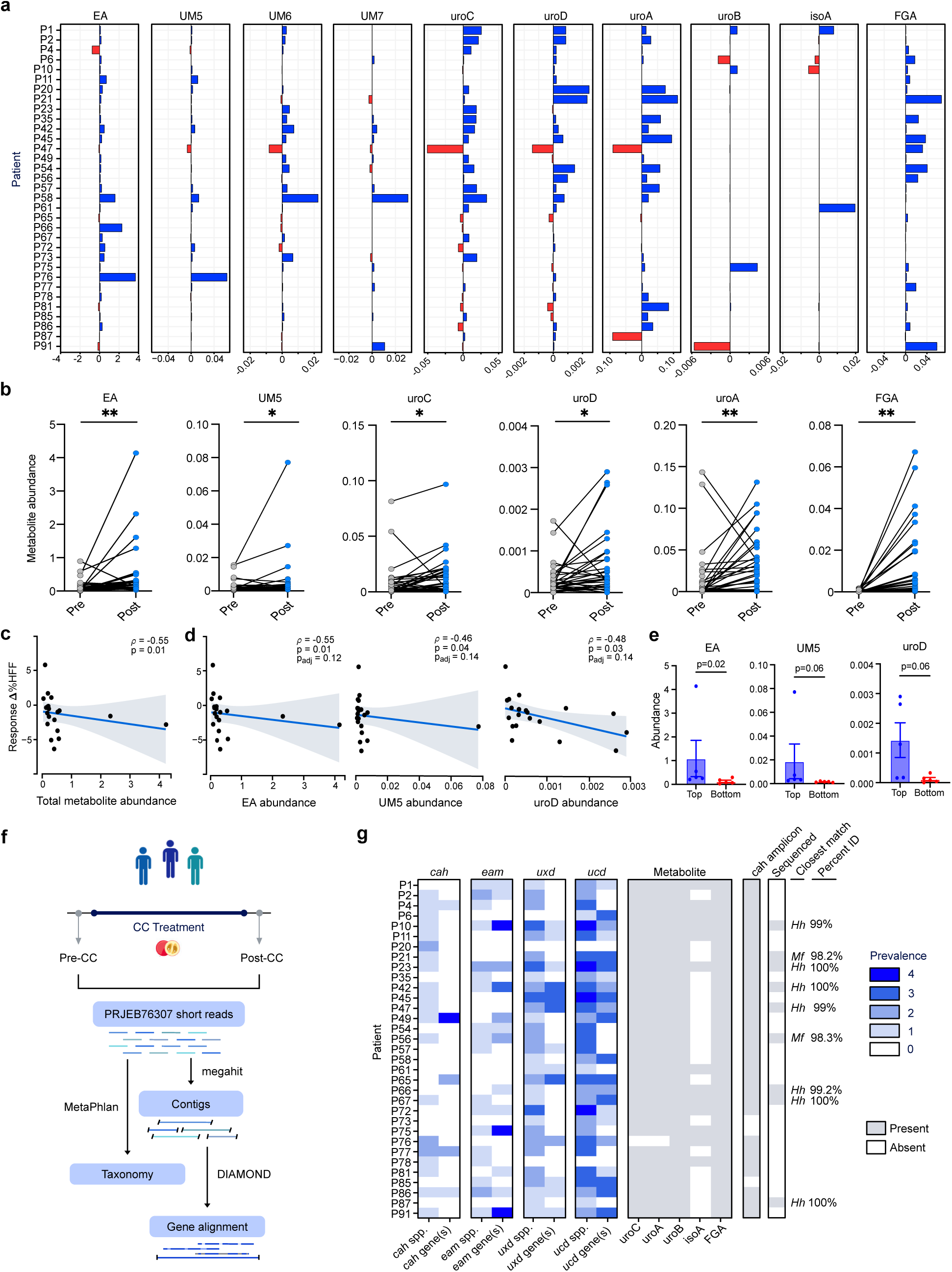
Ellagitannin metabolism genes are prevalent in human microbiome samples from a patient cohort. **a** Difference (post-CC minus pre-CC values) in detected metabolite abundance following camu camu (CC) supplementation per patient. Blue bars indicate an increase in metabolite abundance; red bars indicate a decrease in metabolite abundance. **b** Individual metabolite abundance per patient, pre- and post-CC supplementation. Analyses were performed using a two-tailed paired Wilcoxon signed rank test with Pratt’s method and Holm-Sidak multiple comparison correction. *P ≤ 0.05, **P ≤ 0.01, ***P ≤0.001, ****P ≤ 0.0001. **c** Correlation analysis between total metabolite abundance and change in hepatic fat fraction (HFF) (n=20) **d** Correlation analysis for ellagic acid (EA), urolithin M5 (UM5) and urolithin D (uroD) in the top and bottom responders (n=10). Correlation analyses performed using two-tailed Spearman rho correlation with Benjamini-Hochberg (BH) multiple comparison test. Spearman rho (*ρ*) and FDR (p) values are indicated on associated correlation graph. **e** Abundance of ellagic acid (EA) urolithin M5 (UM5) and urolithin D (uroD) in the top and bottom responders (n=10). Analyses performed using multiple Mann-Whitney test with Holm-Šídák multiple comparison test (p < 0.05). **f** Schematic of bioinformatic workflow for the detection of taxa and homologous ellagitannin metabolism genes from metagenomic reads of patient stool samples. **g** Prevalence of *cah*, *eam*, *uxd* and *ucd* species and genes in patient stool metagenomes post-CC treatment. Species were included in the prevalence count if the relative abundance > 0. Blue gradient indicates number of distinct species associated with each gene cluster. Species included in prevalence counts are as follows: *cah* - *H. hathewayi*, *Blautia* MSK.18.94, *M. formatexigens*; *eam* - *Gordonibacter* spp., *Ellagibacter* spp.; *uxd* and *ucd* - *Enterocloster* spp.. Contigs included in the prevalence count are ≥ 50 amino acids in length with ≥ 97% identity to reference genes. Presence (grey) or absence (white) of representative metabolites urolithin C (uroC), urolithin A (uroA), urolithin B (uroB), isourolithin A (isoA) and flavogallonic acid (FGA) are indicated per patient. A positive PCR-based gene detection result is indicated by a grey box. Amplicons selected for sequencing are indicated in grey with the corresponding closest species match to the queried cahC-D amplicon on the right.

### CC supplementation does not broadly impact microbial composition

To investigate whether a subset of species dictates the observed metabolite production profiles, metagenomic data from this cohort was processed to determine the relative species abundance in patient samples [**Fig. 4f**]. We queried known ellagitannin metabolizers in the *Enterocloster*, *Gordonibacter*, *Ellagibacter*, *Blautia*, *Hungatella* and *Marvinbryantia* genera and found that the validated species responsible for various steps of the metabolism pathway are distributed across the patient cohort [**Fig. 4g**]. While microbial taxa involved in the breakdown of ellagitannins were prevalent in metagenomes, there were no notable taxonomic composition changes at the phylum or species level when comparing pre- and post-CC samples [**Supplementary Fig. 13a**]. Although there were minor, significant differences (P < 0.05) in alpha-diversity, as measured by Shannon and Simpson indices [**Supplementary Fig. 13b**], there was no significant clustering (P > 0.05) between groups based on Bray-Curtis, Jaccard, or robust Aitchison distances [**Supplementary Fig. 13c**]. Additionally, no clear enrichment of taxa relative abundance was observed post-CC when accounting for the number of reads for each sample [**Supplementary Fig. 13d**]. Notably, there were no significant differences in prevalence nor abundance of polyphenol metabolizing species [**Supplementary Fig. 13e**]. Overall, these data suggest that human gut microbial composition is not significantly altered upon CC supplementation and ellagitannin metabolizing species are not differentially abundant in response to treatment. To this end, we focused on post-CC samples for metagenome assembly and protein prediction.

### Castalagin metabolism genes are prevalent in human gut microbiomes

To address whether the presence of metabolizing genes, rather than taxa correlates with observed metabolite profiles, we interrogated the abundance of polyphenol metabolism gene homologues in patient metagenomes. We created a database of protein sequences based on annotated contigs from patient-specific metagenomic data and searched for proteins encoded by *cah*, *eam*, *ucd*, and *uxd* gene clusters from functionally validated microorganisms [**Fig. 4f**]. Homologous protein and protein-coding sequences for *ucd* and *uxd* operons were detected in most patient samples. Conversely, despite detection of *cah* and *eam* species, the genes were sparsely detected [**Fig. 4g**]. Interestingly, in some samples, we detected *cah* and *eam* cluster genes without the presence of known castalagin and EA-metabolizing species. Puzzled by these observations, we investigated whether the apparent absence of *cah* genes in metagenomic datasets reflected limitations in metagenomic sequencing detection. To this end, we designed *cah-*specific primers targeting conserved regions in *Blautia* MSK.18.94, *Mf*, and *Hh,* spanning *cahC-D* genes [**Supplementary Fig. 14a**]. PCR amplification on post-CC stool genomic DNA, revealed the presence of this gene region in nearly all (30/33) patient samples [**Fig. 4g**, **Supplementary Fig. 14b**]. We sequenced selected amplicons and performed homology searches with mmseqs2 using the consensus sequences against reference amplicons from *Blautia* MSK.18.94, *Mf*, and *Hh.* These samples revealed high percent identities with the expected *cahC*-*D* amplicon. Out of nine sequenced amplicons, seven had high protein sequence identity (≥99%) to *Hh*, whereas the two other sequenced samples had high percent identities (>98%) to *Mf* [**Fig. 4g**, **Supplementary Fig. 14c**]. This assay demonstrates that *cah* genes are considerably more prevalent in this patient cohort than initially inferred from metagenomic analyses alone. Together, our meta-analysis and gene-specific PCR assays identified homologues of characterized polyphenol metabolism genes, yet these sequences were not all representative of the functionally validated reference genes. These findings suggest that the prevalent metabolic activity in this cohort is due to the presence of homologous genes in uncharacterized or uncultured gut microorganisms.

To further identify the *cah* homologues in these patient samples, the identified protein sequences were queried against metagenomic datasets from the unified human gastrointestinal proteome (UHGP). This analysis revealed that many of these homologous sequences originate from uncultured and poorly characterized microorganisms, with many matches originating from metagenome assembled genomes (MAGs). Interestingly, several sequences with matches to *cah* genes were assigned to Blautia_A faecis, a species detected in most patient metagenomes **[Supplementary Fig. 15a]**. The taxonomic assignment of this species by MetaPhlAn is limited and the specific strain or reference genome which may be imparting this activity is unknown, therefore making it difficult to validate this observation^49^. These taxonomic classification issues are further exacerbated by differences in NCBI and GTDB taxonomy, which can vary substantially between releases [**Supplementary Table 1**]. Despite limitations in taxonomic validation using this method, the identified homologues from different taxonomic groups may contribute to the metabolic activity observed in the patient samples, explaining the presence of metabolism in patients lacking validated castalagin metabolizers [**Supplementary Fig. 15a, b**].

### Functional homologous polyphenol metabolism genes are found in uncultured bacteria

The microbial taxa involved in polyphenol metabolism are likely underestimated, as most gut bacteria remain uncultured and uncharacterized^50^. In our screen of the castalagin-metabolizing stool sample MO67 [**Supplementary Fig. 2a, b**] we did not observe the canonical species and genes responsible for EA and urolithin metabolism, however there was a metabolite profile consistent with the activities of proteins encoded by the *cah*, *eam*, *ucd*, and *uxd* gene clusters. These trends were also observed in the patient metagenomes above [**Fig. 4g**], suggesting that other taxa are responsible for the observed metabolism profiles. We therefore sought to identify alternative genetic determinants of ellagitannin metabolism using culture-free genome mining approaches that leverage large protein catalogues derived from metagenomes, including MO67, patients in NCT04130321, and the UHGP [**Fig. 5a**], or from genomes in the NCBI.

**Figure 5.**
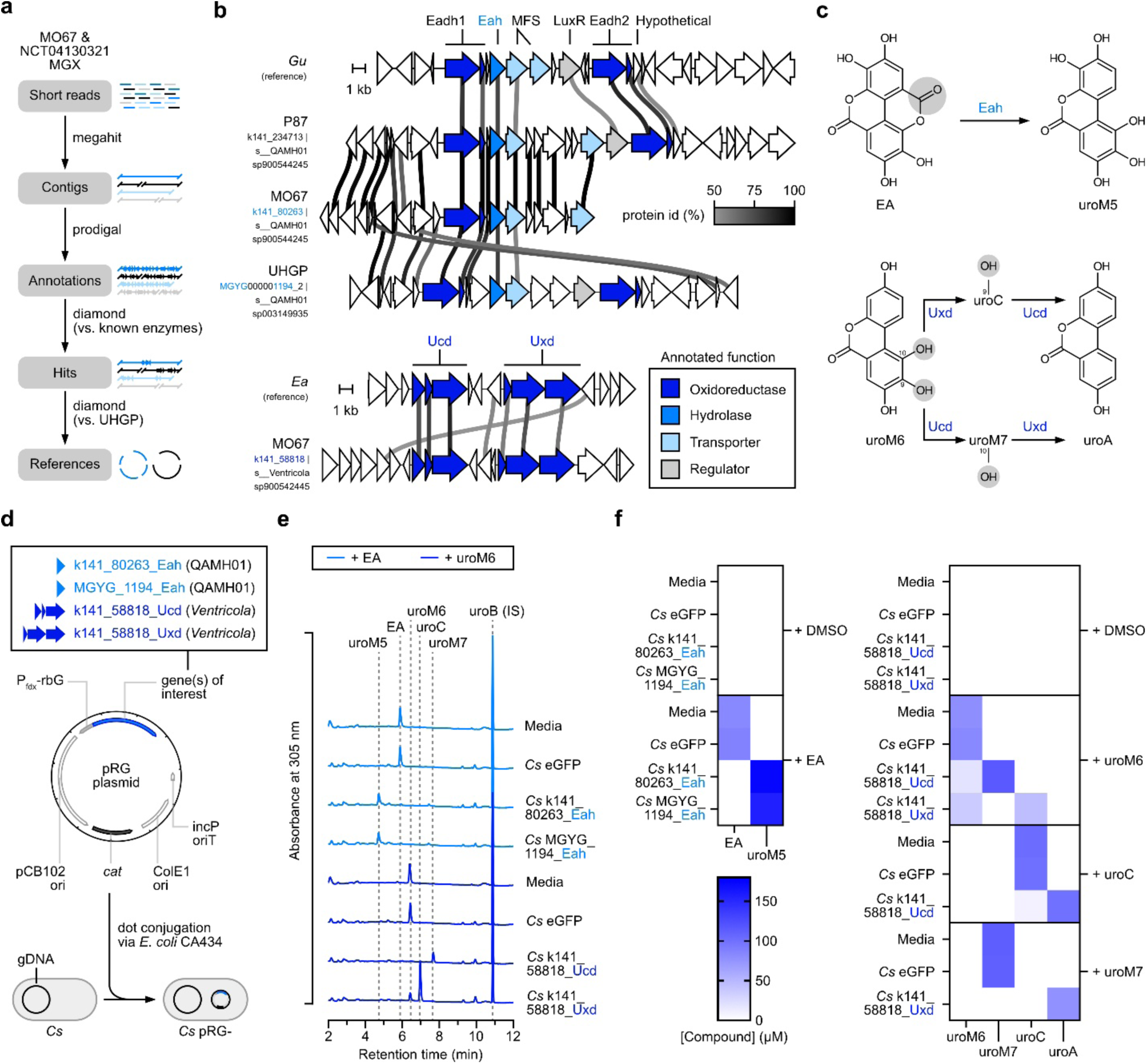
Discovery of homologous polyphenol metabolism enzymes in human gut metagenomes. **a** Bioinformatics workflow to identify polyphenol metabolism enzymes in human gut metagenomes. Protein annotations from assembled contiguous sequences (contigs) were searched against known, functionally validated EA and urolithin-metabolism proteins sequences to identify hits. These hits were further compared to the Unified Human Gastrointestinal Proteome (UHGP) database, which includes GTDB taxonomy (release 202) and more complete metagenome-derived genomes for use as references. **b** Genomic contexts of known polyphenol metabolizers, *G. urolithinfaciens* (*Gu*) and *E. asparagiformis* (*Ea*), with annotated contigs derived from metagenomics assemblies from sample MO67, NCT04130321 patient sample 87 (P87), or the UHGP. Alignments were performed using the clinker tool with genome accessions in **Supplementary Table 7**. Protein sequences with identities ≥ 50% are connected by lines colored according to the percent identity value. GTDB taxonomy (release 232) for each contig is denoted below the contig number. **c** Metabolic reactions catalyzed by putative Eah, Ucd, and Uxd enzymes from uncultured bacteria. Eah is predicted to convert ellagic acid (EA) into urolithin M5 (uroM5) through lactone hydrolysis and decarboxylation. Ucd and Uxd are predicted to dehydroxylate urolithins at the 9- and 10-positions, respectively, yielding a variety of products depending on the substrate urolithin. **d** Schematic representation of the conjugation and heterologous expression workflow to validate Eah, Ucd, and Uxd homologues in *C. sporogenes* (*Cs*) whole cell assays. The generic pRG plasmid map is labeled with the origin of replication (ori), chloramphenicol acetyltransferase (*cat*), and promoter of the *Cs* ferredoxin gene with riboswitch G (Pfdx-rbG). **e** Aligned chromatograms (λ = 305 nm) of substrates and metabolites from mid-exponential phase *Cs* cultures treated with 100 μM of EA or urolithin M6 (uroM6). Tested conditions included sterile media or *Cs* transconjugants expressing enhanced green fluorescent protein (eGFP), k141_80263_Eah, MGYG_1194_Eah, k141_58818_Ucd, or k141_58818_Uxd. Retention times of EA and urolithin standards are shown with grey dotted lines. Data from n = 1 of 3 biological replicates for each transconjugant are shown. **f** Heatmap of EA and urolithin concentrations in cultures (n = 3 biological replicates) shown in (**e**) with additional controls and conditions, including treatments with 100 μM urolithin C (uroC) or urolithin M7 (uroM7).

By searching sequences of known, functionally validated proteins from EA and urolithin- metabolizing bacteria against multiple databases (**Methods**), we identified homologues of Eah, Eadh, Ucd, and Uxd in bacterial genomes derived from human and non-human hosts [**Supplementary Fig. 9b**]. Eah and Eadh proteins were detected in 43 *Coriobacteriales* members that belong to clades within the *Eggerthellaceae* and QAMH01 families that do not cluster with known EA-metabolizing *Gordonibacter* or *Ellagibacter* species [**Fig. 5b, Supplementary Fig. 16a**]. Around 37% (16/43) of these taxa were derived from the human gut, particularly within the QAMH01 family [**Supplementary Fig. 16b**]. Based on the above findings, we selected a subset of hits from diverse genera for further investigation. Inspection of the genomic regions surrounding Eah coding sequences in QAMH01 sp900544245, QAMH01 sp003149935, Adlercreutzia_A murintestinalis, and CAAEEV01 sp011038975 (sp. ZJ176) revealed neighbouring features with high identity to Eadh1 (4-position dehydroxylase), Eadh2 (10-position dehydroxylase), MFS (transporter), and LuxR (regulator) genes in *G. urolithinfaciens* [**Fig. 5b**, **Supplementary Fig. 17a**].

Homologues of Ucd and Uxd dehydroxylases were identified in uncultured *Clostridia* belonging to the *Christensenellales* and *Lachnospirales* orders and were sparsely distributed across multiple taxa [**Supplementary Fig. 9a**]. Despite diverging from urolithin-metabolizing *Enterocloster* spp. at the order level, putative *Ventricola* dehydroxylases are encoded by gene clusters with the same organization as *Enterocloster asparagiformis ucd* and *uxd* operons [**Fig. 5b**]. Based on the protein identities and conserved organization of gene clusters, we hypothesized that the proteins identified above can metabolize EA and urolithins to their respective products [**Fig. 5c**].

Therefore, the genes encoding Eah, Ucd, and Uxd homologues from MO67 contigs k141_80263 (QAMH01 sp900544245) and k141_58818 (*Ventricola* sp900542445) or from the UHGP contig MGYG000001194_2 (QAMH01 sp003149935) were heterologously expressed in *C. sporogenes* [**Fig. 5d**]. An identical approach was taken to validate Eah homologues from CAAEEV01 sp011038975 (sp. ZJ176) and *A. murintestinalis* [**Supplementary Fig. 17b**].

Assays in whole cells confirmed that the identified Eah homologues hydrolyzed and decarboxylated EA to uroM5 in the same manner as *G. urolithinfaciens*, compared to non Eah controls [**Fig. 5e, f, Supplementary Fig. 17c, d**]. Similarly, *Ventricola* sp900542445 Ucd and Uxd homologues dehydroxylated uroM6, uroC, and uroM7 with the same regioselectivity for the 9- and 10-positions [**Fig. 5e, f**]. These experiments validate the predicted EA and urolithin-metabolizing activity of enzymes identified through genome mining and explain the observed urolithins in sample MO67 containing detectable *eam* (QAMH01 sp900544245) and *ucd* (*Ventricola* sp900542445) gene clusters [**Fig. 5b**]. In sum, the genome mining approach described above highlights the potential contribution of multiple uncultured taxa from diverse evolutionary lineages to the EA and urolithin metabolism pathways in humans.

## Discussion

Despite the growing interest in the health benefits associated with ellagitannin metabolism by the gut microbiota, the microbial enzymes associated with conversion of castalagin and vescalagin to ellagic acid had not yet been characterized. In this study we demonstrate that a subset of *Lachnospiraceae* convert castalagin to ellagic acid and castalin via the strain-specific *cah* operon. We further identify FGA as a castalin metabolite detected in human fecal samples. *Cah* does not associate with previously demonstrated urolithin-producing species or previously described tannases. These findings suggest that microbial conversion of ellagitannins is governed by a broader repertoire of specialized enzymes than previously attributed to ellagitannin breakdown and that a minimal community of 3 species is required for castalagin conversion to uroA terminal metabolite.

Due to the frequency of horizontal gene transfer in complex microbial communities, functional acquisition or loss of functional genes are prevalent^51^. Our comparative genomics and metabolism assays highlight strain specificity and non-specific taxonomic distribution of *cah*. These findings in combination with presence of MGEs in a subset of metabolizing species suggest that ellagitannin metabolism has evolved, possibly dependent on the ecological niche the species occupy, contributing to gene acquisition and uneven distribution across genomes^52,53^. The reasons for which these genes may be gained or lost across species and strains remains unknown; however, the ability of organisms within different genera to perform similar functions suggests a decoupling between taxonomic and functional composition. This pattern is consistent with prior observations that microbial communities can maintain conserved functional capabilities despite high taxonomic variability among community members, suggesting a functional redundancy of castalagin metabolism functions^54^.

Our findings demonstrate that combining functional studies with direct metabolite and targeted gene detection reveals functions not captured by metagenomic profiling of clinical datasets, highlighting an important limitation of relying solely on these datasets for functional characterization. Further, limitations in culturing, taxonomic characterization and sequencing depth likely contribute to the incomplete mapping to known species and homologous genes. Additional organisms capable of contributing to this pathway may not yet be represented in reference databases and thus linked with their *cah* homologues. Simply assigning species to a subset of type-strains or model organisms neglects the genetic variations present in pangenomes, thus masking the potential functional capabilities, metabolic activity, and host-microbe interactions^49,55^.

The identification of the *cah* operon allowed us to profile its prevalence in human gut metagenomes from a patient cohort. We highlight that CC supplementation has minimal effects on overall gut microbiome composition, including the abundance of putative metabolizing taxa. These findings suggest that the health benefits of CC may not be mediated by microbiome restructuring but rather through the production of host absorbable metabolites, although plasma or urine urolithins were not measured in this cohort^56,57^. Direct linkage of systemic exposure to ΔHFF was not possible in this dataset and will be addressed in future targeted plasma/urine LC–MS/MS analyses. Despite the low abundance of *cah*-species, we demonstrate that castalagin metabolites, urolithins and flavogallonic acid, are widely detected in the human gut and that the abundance in metabolites post-supplementation correlates with a positive response in change of HFF. While the limited number of patient and the multiple metabolites limit our statistical power, there is a trend between an increase in the intermediate metabolites EA, UM5, and uroD and a decrease in HFF, which does not apply to uroC and terminal metabolite uroA. While this could suggest a role for the intermediate metabolites, it may also be due to increased absorption of the terminal metabolites, limiting their accumulation in fecal samples. Nevertheless, the size of this pilot clinical trial limits our ability to draw additional conclusions regarding the role of these metabolites in metabolic disease. Further validation is required in future studies involving larger cohorts and complementary animal models.

In sum, this study uncovers the previously unknown enzymes and taxa involved in castalagin metabolism, and the metabolites produced from the breakdown of this ellagitannin. As interest in polyphenol supplementation grows and with the increasing number of clinical trials, elucidating how the gut microbiota metabolizes ellagitannins is a crucial step to understanding their effects and the inter-individual responses to prebiotic polyphenols^58,59^. By identifying the first step in castalagin metabolism, our work contributes to the growing body of evidence of the genes and enzymes involved in polyphenol metabolism^26,33,34,60–63^. These findings provide a framework to explain the prevalence of castalagin derived metabolites in the human gut and demonstrate that the functions for castalagin metabolism are widespread. Overall, this knowledge has implications in the development of microbiome-informed therapeutic strategies in the optimization of ellagitannin supplementation in health and disease.

## Methods

### General reagents

A list of general reagents and chemicals is provided in **Supplementary Table 2**.

### Bacterial strains and culturing conditions

The bacterial strains used in this paper are indicated in **Supplementary Table 3**. All bacterial strains were grown on BHI (Difco) agar plates at 37 °C in a Coy vinyl anaerobic chamber, which was maintained with a gas mixture of 3% H_2_, 10% CO_2_, 87% N_2_. All media was reduced in the anaerobic chamber at least 24 h prior to culture setup. To make overnight cultures, a single colony was inoculated into 5 mL of BHI liquid medium. Cultures were incubated at 37 °C for 24-48 h.

### Modified anaerobe basal broth (mABB)

Modified anaerobic basal broth was prepared as previously described^34^. The recipe is available in **Supplementary Table 4**.

### Castalagin and vescalagin extraction

Castalagin and vescalagin were extracted from lightly toasted oak wood chips as described previously^15^. Briefly, extraction from oak chips (Oak Chips Inc. (OH, USA)) was performed in 70% acetone, acetone was removed, and the solution was lyophilized. The dried product was re-extracted in 5% (v/v) acetonitrile, then castalagin and vescalagin were purified by preparative HPLC using the Waters X-select HSS C18 column (10 x 150 mm, 5 μm). Fractions were lyophilized and resuspended in water at a concentration of 5 mg/mL (5.35 mM) for metabolism assays.

### Castalin preparation

Castalin was prepared from castalagin as described previously by Quideau et al.^18^. Briefly, castalagin was dissolved in 10% (v/v) HCl and incubated at 60 °C for 48 hours. The hydrolyzed solution was centrifuged to pellet any precipitates and diluted 5-fold with MilliQ water. The solution was filtered and castalin and vescalin were purified by preparative HPLC using the Waters X-select HSS C18 column (10 x 150 mm, 5 um). Fractions were lyophilized and resuspended in water at a concentration of 5 mM for metabolism assays.

### DNA extraction from isolates

Genomic DNA was extracted from 0.5-1 mL of overnight culture using the One-4-All Genomic DNA Miniprep Kit (BioBasic #BS88503) according to the manufacturer’s instructions. Genomic DNA was then used as PCR templates for cloning and identification. The identities of all bacteria in this study were validated by full-length (V1- V9) 16S rRNA sequencing using primers 16S_V1_27_f and 16S_V9_1492_r listed in **Supplementary Table 5**. Linear/PCR Sequencing was performed by Plasmidsaurus using Oxford Nanopore Technology with custom analysis and annotation.

### Human fecal sample collection

<u>Healthy human samples:</u> Fecal samples were collected from healthy donors under the approval of protocol A04-M27-15B by the McGill Faculty of Medicine Institutional Review Board. Participants provided informed written consent for the collection and use of the fecal samples and were compensated $10 CAD for their participation. The eligibility criteria for the healthy donors were as follows: no diagnosed gastrointestinal disease, no ongoing therapeutic treatment, and no antibiotic usage 3 months prior to the start of the study. Donor information was recorded at the time of sample collection and is provided in **Supplementary Table 6**. Donors were between 21-33 years old with BMI between 22.0-26.8. The fecal samples were collected in 2024, immediately transferred to an anaerobic chamber to be aliquoted and were stored at −70°C until further use.

<u>Overweight and hypertriglyceridemia patients:</u> Stool samples were collected as previously described^19^. Briefly, samples were collected using the EasySampler Stool Collection Kit (ALPCO) prior to and after camu-camu treatment. Samples were immediately stored at −20°C until transferred to −80°C until further processing for analysis.

### Fecal sample preparation and treatment

Frozen stool aliquots (−70°C) were transferred to an anaerobic chamber and thawed. 1 mL mABB media was added per 0.1 g of stool and samples were homogenized by vortexing and using a sterile loop to break apart larger pieces. Samples were centrifuged at 700 g x 3 min to pellet larger particles. The supernatant was carefully collected, transferred to new tubes and the bacterial cells were pelleted (6,500 g x 3 min). The supernatant was discarded, and the bacterial pellet was washed with 5 mL mABB media. Resuspension was pelleted (6,500 g x 3 min), the supernatant discarded, and the cell pellet was resuspended in 300 μL mABB media per 0.05 g of starting stool. Fecal slurries were aliquoted into sterile tubes and treated with castalagin to a final concentration of 100 μM or an equal volume sterile H_2_O. Samples were incubated anaerobically at 37°C. 200 μL of slurry was collected at timepoint 0, 4, 6 and 24 h post-treatment and stored at −70°C for metabolism analysis by LC-MS using extraction method A.

### RNA extraction from fecal samples

Aliquots of 500 µL of fecal slurry (treated with castalagin or sterile H_2_O) were pelleted (10,000 g x 3 min). The supernatant was removed, and the pellets were resuspended in 800 μL of TRI Reagent (Zymo #R2050-1). The resuspended pellets were transferred to be ZR BashingBead lysis tubes (0.1 & 0.5 mm) (Zymo #S6012-50) and lysed by bead beating on a Vortexer at maximum speed for 40 minutes. RNA isolation was performed using the Direct-zol RNA Miniprep Kit (Zymo #R2051) according to the manufacturer’s instructions. An additional DNase digestion was performed on eluted DNA using the DNA- *free*™ DNA Removal Kit (Invitrogen #AM1906) according to the manufacturer instructions. The OneStep PCR Inhibitor Removal kit (Zymo #D6030) was used to further clean up the RNA. RNA quality and concentration was primarily verified by NanoDrop (Thermo Fisher ND-2000C) and 1% agarose gel electrophoresis in TAE buffer. For RNA sequencing, the samples were sent to Génome Quebec and RNA integrity was performed on total RNA and evaluated using the Bioanalyzer 2100 (Agilent). RNA integrity values (RIN) for castalagin- treated and vehicle-treated samples ranged from 5.4 - 6.1 and 5.8 - 6.9 respectively.

### Metatranscriptomics sequencing

Total extracted RNA was sent to Génome Québec for RNA sequencing. rRNA was removed from total RNA using the NEBNext rRNA Depletion Kit (Bacteria) (NEB). Paired- end libraries were prepared using the NEBNext Multiplex Oligos for Illumina kit (NEB) and sequenced on an Illumina NovaSeq instrument (PE100 – 25 M reads). Raw read FastQC (v0.12.1) output was as follows: 26.1-47.2 M reads, 43-46 % GC, mean quality 39, 0.023-0.052 % adapter content.

### Metatranscriptomic processing and statistical analysis

Paired-end metatranscriptomic reads were adapter- and quality-trimmed using Trim Galore^64^ (v0.6.10) and Cutadapt^65^ (v2.10), with a Phred quality cutoff of 5 and a minimum read length of 75 bp. Terminal Ns were trimmed, and reads containing ambiguous bases were excluded. Host reads were removed by mapping against the human reference genome hg38 using BBMap^66^ (v38.90), with a minimum identity of 0.97, a maximum indel length of 3 and a minimum of two seed hits (minhits=2), in slow mode with qtrim=rl, trimq=10 and untrim. Residual ribosomal RNA (rRNA) reads were removed using SortMeRNA^67^ (v2.1b) against its SILVA bacterial, archaeal and eukaryotic rRNA reference databases and Rfam 5S/5.8S reference databases, with an E-value threshold of 10⁻³⁵ and paired_in mode.

For assembly, non-rRNA reads were pooled by treatment and reduced using ORNA^68^ with quality-based sorting (sorting=1, base=1.7, kmer=21). The four treatment pools were co-assembled using MEGAHIT^69^ (v1.2.9), with the meta-large preset and a minimum contig length of 200 bp, yielding 260,361 contigs. Each sample’s non-rRNA reads were then pseudoaligned to this common assembly using kallisto^70^ (v0.46.2) in paired-end mode, without bootstrap replicates. Sample-level quantification used the reads before ORNA reduction.

Kallisto abundance estimates and effective lengths were imported from abundance.h5 files using tximport ^71^ (v1.38.2) and analysed at the contig level using DESeq2^40^ (v1.50.2) in R (v4.5.0). The DESeq2 dataset was constructed using DESeqDataSetFromTximport, incorporating the imported count estimates and effective-length information for normalization. All 16 endpoint samples were analysed jointly using an additive model that included block and treatment effects (∼ Block + Treatment). Each matched block (A–D) contained one aliquot for each of the four conditions: vehicle, castalagin, castalin and EA. Contigs with a count of at least 10 in at least four samples were retained, yielding 64,749 contigs for analysis.

Four preplanned comparisons were evaluated using Wald tests: castalagin versus vehicle, castalin versus vehicle, EA versus vehicle, and castalagin versus EA. Positive log2 fold changes indicated higher transcript abundance in the first-named condition. Two-sided P values were adjusted using the Benjamini–Hochberg procedure^72^ independently within each contrast, with DESeq2 independent filtering at alpha=0.05. Statistical significance was defined as an adjusted P value < 0.05, without an additional fold-change cutoff. Reported log2 fold changes were not shrunken. Contigs excluded by independent filtering retained their effect estimates but were not assigned an adjusted P value and were not counted as significant.

The overall effect of treatment was assessed for each retained contig using a likelihood- ratio test comparing the full model, which included block and treatment effects, with a reduced model containing block alone (3 degrees of freedom). P values from these tests were adjusted using the Benjamini–Hochberg procedure to control the false discovery rate at 5%. Variance-stabilized counts were used for descriptive quality checks based on principal component analysis and sample-to-sample distances.

The four blocks represented replicate incubations from a single donor rather than independent human donors; inference was therefore restricted to the MO67 incubation system. Vehicle served as a parallel endpoint control, and no metatranscriptomic time- zero sample was analysed.

### DNA extraction from fecal samples

Fecal slurries of MO67 from each experimental replicate (pre-treatment) were pooled and frozen at −70 °C before DNA extraction. Fecal samples from NCT04130321 were thawed and aliquoted (150-250 mg aliquots) before DNA extraction. Total DNA was extracted from using the ZymoBIOMICS DNA Miniprep kit (Zymo #D4300), which included a bead beating step using ZR BashingBead lysis tubes (0.1 & 0.5 mm) (Zymo #S6012-50) on a Vortexer (maximum speed x 40 minutes) or using a Bead Mill homogenizer (3 cycles x (1 min ON, 6 m/s, 5 min rest) for a total of 3 minutes of bead beating). Total DNA from each aliquot was eluted in 50 μL of nuclease-free H_2_O.

### Metagenomics sequencing

Total isolated DNA from fecal sample MO67 was sent to Génome Quebec for DNA sequencing. A paired-end DNA library was prepared using IDT dual-index adapters and sequenced with an Illumina NovaSeq instrument (PE150 – 35 M reads). Raw read FastQC (v0.12.1) output was as follows: 112.7 M reads, 45 % GC, mean quality 39, 0.027 % adapter content.

### Metagenomic processing and statistical analysis

Paired-end raw metagenomic reads obtained from Genome Quebec (for MO67) or from the European Nucleotide Archive (PRJEB76307) were trimmed using fastp (v1.0.1) with the following flags: --detect_adapter_for_pe --trim_poly_g --cut_front --cut_tail -- cut_window_size 4 --cut_mean_quality 20 --length_required 100. After trimming, reads were mapped to the T2T-CHM13v2.0 human genome (GCF_009914755.1) to remove host sequences using bowtie2 (v2.5.4) with default parameters. Gzipped unaligned concordant (--un-conc-gz) reads were used for downstream analyses. At each step of quality control, reads were verified using FastQC (v0.12.1).

For community composition analysis, quality controlled reads were analysed using MetaPhlAn (v4.1.1) and its accompanying database mpa_vJun23_CHOCOPhlAnSGB_202403. MetaPhlAn taxonomy profiles (species-level genome bins, SGB) were converted to GTDB taxonomy (release 207) using the sgb_to_gtdb_profile.py utility script. Diversity analyses and multivariable associations were performed using the vegan (v2.7-2) and MaAsLin3 (v1.1.2) R packages, respectively.

For the analysis of gene prevalence and abundance in metagenomes, quality controlled paired-end reads from the same individual (2 timepoints per individual) were co- assembled into contiguous sequences using megahit (v1.2.9) with the following flags: -- presets meta-sensitive --min-contig-len 500. Protein coding genes were predicted from the resulting contigs using prodigal (v2.6.3) with the following flags: -p meta -g 11. A simplified annotation format (SAF) file and a gene lengths file were generated from the prodigal-predicted genes. Quality controlled paired-end reads (individual timepoints) were mapped onto contigs using bowtie2 and the resulting SAM alignment file (.sam) was converted to BAM (.bam), sorted, and indexed using samtools (v1.22.1). Reads per predicted gene (based on the SAF file) were determined using the featureCounts tool in subread (v2.0.6) with the following flags: -p -B -C. Reads per kilobase per million mapped reads (RPKM) and transcripts per million (TPM) were calculated from featureCounts output.

### Identification of *cah*-harbouring species

Searches for culturable bacterial species harbouring sequences similar to upregulated transcripts of interest (nucleotide) were performed using the blastn tool against refseq_genomes and the organism limited to Bacteria (taxid:2). Searches for sequences similar to individual genes of interest from the *cah* gene cluster were performed using the blastx tool against the refseq_protein database, limiting the genetic code to Bacteria and Archaea (11) and the organism to Bacteria (taxid:2).

### Comparative genomics and bacteria genome mining

CAGECAT (https://cagecat.bioinformatics.nl/, v1.0) was used to make comparisons between genomic contexts, using the clinker tool for visualization^73^. GenBank files from accessions in **Supplementary Table 7** were downloaded from the NCBI Sequence Viewer and uploaded into clinker. Individual proteins were manually coloured to indicate different functions among the gene clusters. Individual protein percent identity and similarity matrixes were produced using the alignment.txt files generated in clinker. Cutoff percent identities used for analysis in clinker are indicated in associated figure legends. To determine the presence of *cah* and *act* genes in select *Lachnospiraceae* genomes, we downloaded all available genomes (n= 4699) from *Blautia*, *Enterocloster*, *Hungatella*, and *Marvinbryantia* in the NCBI genomes database (accessed 2026-01-05) including MAGs and atypical genomes. The genome table for the specified genera from the Genome Taxonomy Database (GTDB) (release 10-RS226 (2025-04-16)) (accessed 2025-08-06) were downloaded. Each genome was searched for individual genes belonging to the *cah* and *act* gene cluster from *Blautia* MSK.18.94 using rBLASTx (v1.2.0, doi.org/10.18129/B9.bioc.rBLAST)^74^ as done previously (doi.org/10.5281/zenodo.16740616)^26^. The criteria for matches were as follows: percent identity >60 (perc_identity 60), max_target_seqs 5 and subject_besthit. The identified matches were then filtered for sequences greater than specified lengths (> 200 for cahA, cahC, cahD, actA, actB1, actB2, actC1, actC2, > 100 for cahB and actD and >300 for actE). Only GenBank accessions that had corresponding GTDB entries were retained.

To generate a list of genes encoding proteins of interest (POI) involved in ellagitannin metabolism from human gut metagenomes, POI were aligned to assemblies using diamond (v2.1.11) with the following flags: --id 60 --subject-cover 60 -k 0. Hits from this initial protein-protein search were then compared to the UHGP-100 protein catalogue using diamond (with the same parameters as above). Results tables from both searches were merged with the UHGP metadata table (genomes-all_metadata.tsv) using pandas (from the scipy-stack/2026a package).

To identify taxa predicted to metabolize ellagic acid, genomes from o Coriobacteriales (GTDB release 232) were downloaded from the NCBI (n = 6,667 genomes). Proteins were predicted using pyrodigal (v3.5.2) with the following flags: -p meta -g 11. Functionally-validated proteins encoded by the *eam* gene cluster (Eah, Eadh1, Eadh2, Eadh3) of *A. murintestinalis*, CAAEEV01 sp011038975 (sp. ZJ176), *G. urolithinfaciens*, *G. pamelaeae*, *Gordonibacter* sp. 28C, *G. massiliensis*, *E. isourolithinifaciens*, QAMH01 sp003149935, and QAMH01 sp900544245 were used as queries to search for homologues in pyrodigal-predicted proteins from each genome with mmseqs2 easy- search with the following flags: --min-seq-id 0.5 -c 0.5. Taxa with at least 1 genome containing Eah and ≥ 1 Eadh homologue with percent identities greater or equal to 70 were predicted to metabolize ellagic acid and annotated as such in the o Coriobacteriales phylogenetic tree.

To identify taxa predicted to metabolize castalagin or to produce uroA, genomes from o Christensenellales and o Lachnospirales (GTDB release 232) were downloaded from the NCBI (n = 53,487 genomes). Proteins were predicted using pyrodigal (v3.5.2) with the following flags: -p meta -g 11. Proteins encoded by the *cah* gene cluster (CahA, CahB, CahC, CahD) of *Blautia* MSK.18.94, *H. hathewayi*, *M. formatexigens*, *E. clostridioformis* YL32, sample MO67 and proteins encoded by the *ucd* and *uxd* operons (UcdC, UcdF, UcdO, UxdF, UxdO, UxdC) of *E. asparagiformis*, *E. bolteae*, *E. citroniae*, and *E. hominis* (formerly *E. pacaense*) were used as queries to search for homologues in pyrodigal-predicted proteins from each genome with mmseqs2 easy-search with the following flags: --min-seq-id 0.5 -c 0.5. Taxa with at least 1 genome containing 3/4 Cah or 3/3 Ucd/Uxd proteins with percent identities greater or equal to 70 were predicted to metabolize castalagin or to produce uroA and annotated as such in the c Clostridia phylogenetic tree.

### Metabolism assays in isolates

Overnight cultures of *Lachnospiraceae* species were diluted 1/50 in fresh BHI media. During growth, castalagin, vescalagin, punicalagin, corilagin or an equal volume of sterile H_2_O were added to a final concentration of 100 μM. Castalin was added to a final concentration of 200 μM. Cultures were incubated at 37 °C under anaerobic conditions for 48 h. Samples were collected and stored at -70 °C until metabolite extraction. Samples were prepared as per extraction method A, and the presence of metabolites in culture were detected using HPLC-MS.

### Metabolism assays in synthetic communities

Overnight cultures of *H. hathewayi* DSM 13479 *G. urolithinfaciens* DSM 27213, *E. bolteae* DSM 15670 and *E. asparagiformis* DSM 15981 were diluted 1/100 (each) into fresh BHI media (5 mL) according to the following (5) treatment groups: media only, DSM13479, DSM13479 + DSM27213, DSM 27213 + DSM 15670, DSM27213 + DSM15981, DSM13479 + DSM 27213 + DSM 15670, DSM13479 + DSM 27213 + DSM15981. Each treatment group was split into 3 X 500 µL aliquots and treated with water or castalagin (5.35 mM stock) to a final concentration of 100 μM. Samples were incubated at 37 °C under anaerobic conditions and 200 μL samples were taken after 5 days and stored at - 70°C until metabolite extraction. Samples were prepared as per extraction method A, and the presence of metabolites in culture were detected using HPLC-MS.

### Metabolite extraction from cultures

*Extraction method A*: Frozen (−70 °C) bacterial cultures were thawed at room temperature. For quantification of EA or FGA concentration, an EA (stock 10 mM in DMSO) or FGA (stock 5 mM in DMSO) standard was prepared in an aliquot of BHI prior to extraction. Each sample was incubated at 40°C for 2 minutes prior to adding salicylic acid at a final concentration of 50 µg/ml. An equal volume of 100% methanol (1:1 sample:methanol) was added to each sample and the samples were incubated at room temperature for 10 minutes. Samples were centrifuged at 20,000 x g for 5 min to pellet insoluble material, then the supernatant was transferred to LC-MS vials. The presence of ellagic acid was then analyzed by LC-MS.

### Metabolite extraction from fecal samples

Metabolites were extracted from stool samples using *Extraction method B*: Frozen stool samples were thawed and resuspended in 1X PBS at a ratio of 300 µL / 100 mg. Homogenization was done by vortexing and using a sterile loop to break apart larger pieces. Samples were centrifuged 700 x g for 5 min at room temperature to pellet large particles. The supernatants were collected and transferred to new tubes. Salicylic acid at a final concentration of 50 μg/mL (3 mg/mL in DMSO stock) was added as an internal standard to supernatants and metabolite standards. Extraction was performed by adding 4x the volume of ethyl acetate + 1% formic acid to each sample volume (e.g. 800 µL to 200 µL of supernatant). Samples were centrifuged; the organic layers were collected, transferred to new tubes, and dried using a SpeedVac vacuum concentrator connected to a rotary evaporator (Buchi). Dried samples were resuspended in 150 µL of 50% methanol, centrifuged to pellet insoluble products, and transferred to LC-MS vials. Samples were submitted for targeted metabolomics.

### Metabolite analysis by LC-MS

Samples were injected (10 μL) into a 1260 Infinity II Single Quadrupole LC/MS system (Agilent Technologies) fitted with a Poroshell 120 EC-C18 4.6 × 50 mm, 2.7 μm column (Agilent Technologies) with a corresponding guard column maintained at a temperature of 30°C. The mobile phase was MilliQ water with 0.1% formic acid (Solvent A) and acetonitrile with 0.1% formic acid (Solvent B). The flow rate was set to 0.7 mL/min. The gradient to detect ellagic acid and urolithins was as follows: 0–8 min: 10-30 %B, 8–10 min: 30-100 %B, 10–13.5 min: 100 %B isocratic, 13.5-13.6 min: 100-10 %B, then 13.6–16.5 min: 10 %B. The gradient to detect castalin and flavogallonic acid was as follows: 0– 3 min: 0 %B, 3–7 min: 0-10 %B, 7–11.5 min: 10-20 %B, 11.5-16 min: 20-30 %B, 16–20.5 min: 30-50 %B, 20.5-23 min: 50-100 %B, 23-25 100%B isocratic, then 25-30 100-0%B. The mass spectrometer (API-ES) was run in negative mode in both scan (100– 1000 m/z) and selected ion monitoring (SIM) modes to validate peak identities. Capillary voltage was set to 3000 V, drying gas to 10.0 L/min, nebulizer pressure to 30 psig, and drying gas temperature to 350 °C. The multiple wavelength detector monitored absorbance at 305 nm. Detected peaks were validated by comparison of retention times to that of spike-in standards and mass-to-charge ratios. The integrated peak areas derived from the Agilent ChemStation software were compared to standards of known concentration for metabolite quantification. When standards were not available or overlapping peaks could not be resolved, the extracted ion chromatogram peak area was used. Ellagic acid [M-H]^−^: 301, urolithin M5 [M-H]^−^: 275, urolithin E [M-H]^−^: 259, urolithin D [M-H]^−^: 259, urolithin M7 [M-H]^−^: 243, urolithin M6 [M-H]^−^: 259, urolithin C [M-H]^−^: 243, urolithin G [M-H]^−^: 243, urolithin A [M-H]^−^: 227, isourolithin A [M-H]^−^: 227, urolithin B [M- H]^−^: 211, flavogallonic acid [M-H]^−^: 469, castalin [M-H]^−^: 631.

### Plasmid construction in *E. coli*

All plasmids used in this study were constructed through DNA assembly using the NEBuilder HiFi DNA Assembly Master Mix (NEB #E2621). Linearized pRG_BB (backbone) plasmid DNA was generated through PCR amplification of the pRG-Cas2 plasmid (University of Nottingham Synthetic Biology Research Centre) using the Q5 DNA polymerase (NEB # M0491). To reduce contamination from template pDNA, DpnI (NEB #R0176) was added to each reaction after thermal cycling and incubated for 1 h at 37 °C. DNA inserts containing individual genes or operons of interest were either generated through PCR amplification (using the Q5 DNA polymerase) or through commercial gene fragment synthesis (GenScript). In all cases, Q5 DNA polymerase PCR cycling was as follows: 30 s at 98 °C, 30 cycles (10 s at 98 °C, 20 s at Ta, reaction-specific extension time at 72 °C), 2 min at 72 °C. Primer sequences are listed in **Supplementary Table 5** and specific PCR conditions (primers, Ta, and extension time) are listed in **Supplementary Table 8**. PCR products were gel purified using the Monarch DNA Gel Extraction Kit (NEB #T1120) according to the manufacturer’s instructions for different DNA sizes. DNA fragments were mixed for HiFi DNA assembly at 50 °C for 1 h according to the manufacturer’s instructions. HiFi DNA assembly constructs (plasmid backbones and inserts) are provided in Supplementary Table **9**.

HiFi DNA assembly reactions were transformed into homemade CaCl_2_ competent *E. coli* NEB10β (NEB #C3019) via heat shock (30 s at 42 °C). The cells were recovered in SOC broth for 1 hour at 37°C and spread onto selective LB + 30 μg/mL chloramphenicol agar plates. The plates were incubated aerobically at 37 °C for 24 h. The next day, colonies were picked and grown in selective LB + 30 μg/mL chloramphenicol. Plasmids were purified using the Plasmid DNA Miniprep Kit (BioBasic #BS414) and size was confirmed with a diagnostic restriction digest. Whole plasmid sequencing was performed by Plasmidsaurus using Oxford Nanopore Technology with custom analysis and annotation. GenBank accessions for sequenced plasmids are listed in Supplementary Table **10**. Plasmid maps were generated in Benchling (https://benchling.com).

### Plasmid conjugation into *C. sporogenes*

CaCl2 competent *E. coli* CA434 conjugal donor cells were transformed with pRG series plasmids via heat shock (45 s at 42°C). The cells were recovered in SOC broth for 1 hour at 37°C and spread onto selective LB + 30 μg/mL chloramphenicol + 15 μg/mL tetracycline agar plates. The plates were incubated aerobically at 30 °C for 48 h. Single colonies of *E. coli* CA434 transformants harbouring the plasmids were inoculated into 5 mL LB + 30 µg/mL chloramphenicol + 15 µg/mL tetracycline and grown under aerobic conditions at 32°C for 48 h with shaking. The day before conjugation, a single colony of *C. sporogenes* was inoculated into 5 mL of BHI and incubated anaerobically at 37 °C overnight. 1.5 mL of the *E. coli* CA434 door was pelleted at 6,500 g x 2 min, washed with 1 mL 1X sterile PBS, re-pelleted, and brought to an anaerobic chamber on ice. An aliquot of *C. sporogenes* culture was heat shocked at 50°C for 5 min (aerobically) and subsequently returned to the anaerobic chamber. The *E. coli* pellet was resuspended in 300 µL of the heat shocked recipient culture and the mixture was spotted (3 x 20 µL) onto a BHI agar plate and grown at 37°C anaerobically for 24 h. The next day, dots were scraped with a sterile inoculating loop and resuspended in 300 µL 1X PBS (sterile and pre-reduced). 50-100 µL of the re-suspension was spread on BHI + 250 µg/mL D- cycloserine + 30 µg/mL thiamphenicol agar plates. Plates were incubated anaerobically at 37°C for 24-48 h. Individual transconjugants were picked and re-streaked on BHI + 250 µg/mL D-cycloserine + 30 µg/mL thiamphenicol, then on BHI + thiamphenicol 30 µL/mL agar plates. From re-streaks, colonies were inoculated into 5 mL BHI + 30 µL/mL thiamphenicol and grown overnight at 37°C.

### Colony PCR

To validate plasmid-positive *C. sporogenes* colonies, colony PCR was performed on overnight cultures using the OneTaq 2X Quick-Load Master Mix (NEB #M0486) with cat_cPCR primers [**Supplementary Table 5**]. An aliquot of overnight culture was heated at 95°C for 3 min and 1 µL of heated sample was used as the template for cPCR. PCR was performed as follows: 5 min at 94 °C, 35 cycles (20 s at 94 °C, 30 s at 45 °C, 45 s at 68 °C), 5 min at 68 °C. Presence of the gene was visualized by 1% agarose gel electrophoresis and compared to a 1 kb Plus DNA Ladder (NEB #N0550).

### Heterologous expression in *C. sporogenes*

All growth steps were performed in selective media (BHI + 30 μg/mL thiamphenicol) in anaerobic conditions at 37 °C. Single transformant colonies of *C. sporogenes* grown on BHI + 30 μg/mL thiamphenicol agar plates, were inoculated into 5 mL selective media and grown overnight. Overnight cultures were diluted 1/50 into 5 mL fresh selective media and grown for 6 h. Cultures were treated with 5 mM theophylline (100 mM stock in 0.1 M NaOH) to induce protein expression and incubated overnight..

For castalagin metabolism assays, 2 mL of culture was pelleted 6,000 x g x 5 min at 4°C and resuspended in 1 mL ice-cold 1X PBS, then disrupted by sonication on ice using a Misonix Sonicator 3000 with an output power of 2/10, 20 s ON, 40 s OFF, for a total of 4 min 20 s. Insoluble material was pelleted 20,000 x g x 2 min and the soluble fraction was transferred to an anaerobic chamber on ice. Activity assays were set up by aliquoting 200 μL of soluble lysate and adding castalagin to a final concentration of 300 μM. Lysates were incubated at 37 °C and collected at 48- post-treatment.

For ellagic acid or urolithin metabolism assays, cultures were treated with 100 μM EA, uroM6, uroM7, or uroC at the same time as the theophylline addition and incubated at 37 °C for 24 h. Samples were frozen at -20 °C until extraction and analysis by LC-MS.

### Targeted Metabolomics

Samples were injected (5 µL) onto an Agilent 6470 Triple Quadrupole (QQQ)–LC– MS/MS. Chromatographic separation of metabolites was done using the ultra-high pressure liquid chromatography (UHPLC) 1290 Infinity LC System (Agilent Technologies). The mobile phase was water + 0.1% formic acid (Solvent A) and acetonitrile + 0.1% formic acid (Solvent B). The flow rate was set to 0.4 mL/min. The gradient for ellagic acid and urolithin separation was as follows: 0-8 min: 10-30% B, 8-10 min: 30-100% B, 10-13.5 min: 100% B, 13.5-13.6 min: 100-10% B, 13.6-15.5 min: 10% B. For ellagic acid and urolithin analysis, metabolite separation was achieved using a ZORBAX Eclipse Plus C18 column, 1.8 µm, 2.1 x 150 mm with guard column (Agilent Technologies), the column was maintained at 30°C. The gradient for flavogallonic acid separation was as follows: 0-3 min: 0% B, 3-9 min: 0-15% B, 9-10 min: 15-100% B, 10-12 min: 100% B, 12-12.1 min: 100-0% B, 12.1-15 min: 100% B. For flavogallonic acid analysis metabolite separation was achieved using a ZORBAX Eclipse Plus C18 column, 1.8 µm, 2.1 x 50 mm with guard column (Agilent Technologies), the column was maintained at 10°C. The mass spectrometer was equipped with an electrospray ionization source and run in negative mode for urolithin detection and positive mode for flavogallonic acid detection. The source gas temperature was set to 300°C and the gas flow rate to 5 L/min. The nebulizer pressure was 45 psi. The sheath gas temperature and sheath gas flow were 250°C and 11 L/min respectively, and capillary voltage was 3500 V.

Analysis of metabolomic data was done using the MassHunter Software. Peaks were manually integrated for each individual metabolite to select for correct retention time and peak area corresponding to analytical standards. Due to reduced detection sensitivity, EA peak areas had higher normalized values, despite reduced signal, and as such were analyzed separate from urolithin metabolites.

### RNA extraction from isolates

*Blautia* MSK.18.94 was diluted 1/50 in 15 mL of BHI and grown at 37 °C anaerobically. After 24 h growth, the culture was split into 2 X 5 mL aliquots and treated with either water or castalagin (50 µM final concentration) for 24 h at 37 °C. Cultures were pelleted (6,500 g for 3 min), washed with 5 mL of pre-reduced PBS, and re-pelleted. The pellet was resuspended in 900 µL TRI Reagent (Zymo #R2050-1), transferred to ZR BashingBead lysis tubes (0.1 & 0.5 mm) (Zymo #S6012-50), and lysed by bead beating on a Vortexer set to maximum speed for 40 minutes. RNA isolation was performed using the Direct-zol RNA Miniprep Kit (Zymo #R2051) according to the manufacturer’s instructions (with on- column DNaseI digestion). An additional DNase digestion was performed on eluted RNA using the DNA-*free*™ DNA Removal Kit (Invitrogen #AM1906) according to the manufacturer instructions. DNase-treated RNA was additionally cleaned up using the OneStep PCR inhibitor removal kit (Zymo #D6030) according to the manufacturer instructions. RNA concentrations were determined using the Qubit RNA BR kit (Invitrogen #Q10210) and RNA quality was verified by 1% agarose gel electrophoresis.

### RT-PCR for operon determination

RT-PCR experiments were performed in 2 steps. Isolated RNA was reverse transcribed using the LunaScript RT Master Mix (primer-free) kit (NEB # E3025) with reverse primer MSK.18.94_RT-PCR_R2 [**Supplementary Table 5**]. Briefly, 500 ng of RNA were reverse transcribed in a reaction volume of 10 μL. The reverse primer was added to a final concentration of 1 μM. For No-RT controls, the No-RT Control Mix provided in the kit was used with the same RNA and primer amounts. Reactions were incubated at 55 °C for 10 min and 95 °C for 1 min. RT reactions were then diluted 1/20 in nuclease-free water and used as the template for PCR. PCR reactions were done using the OneTaq Quick-Load 2X Master Mix with Standard Buffer (NEB #M0486) with primer pairs MSK.18.94_RT- PCR_F4 and MSK.18.94_RT-PCR_R2 [**Supplementary Table 5**]. Cycling conditions were as follows: 94 °C for 30 s, 35 cycles of (94 °C for 20 s, 45 °C for 30 s, 68 °C for 4.5 min (∼1 min/kb)), and 68 °C for 5 min. Positive and negative controls were MSK.18.94 genomic DNA and nuclease-free water, respectively. Gel electrophoresis of PCR products was done using a 0.7% agarose gel and products were compared to the Quick- Load Purple 1 kb Plus DNA ladder (NEB #N0550).

### Phylogenetic trees

Phylogenetic trees based on Genome Taxonomy Database taxonomy (Bacteria genome trees from a set of 120 single copy marker proteins) were generated using PhyloT (v2, https://phylot.biobyte.de/) using accessions from the GTDB release 232 or GTDB release 220. Phylogenetic trees were edited (collapse clades) and exported using iTOL (v7, https://itol.embl.de/). Trees were manually annotated in Affinity Designer 2.

### Protein extraction of isolates for proteomics

All steps other than sonication were performed under anaerobic conditions. Overnight cultures of *H. hathewayi* and *E. clostridioformis* YL32 were diluted 1/50 in BHI and grown at 37 °C. After 18 h, 50 μM of castalagin, castalin or equal volume vehicle (H_2_O) were added to the cultures and incubated for 5 hours. 5 mL of culture was pelleted 6000 g x 5 min, the pellets of Hh and Ec were pooled according to the condition and resuspended in 5 mL 1X PBS. Samples were pelleted and resuspended in 0.5 mL ice-cold lysis buffer: filtered 1X PBS + 1 tablet/10 mL cOmplete mini protease inhibitor EDTA-free (Roche). The resuspended pellets were disrupted by sonication using a Misonix Sonicator 3000 set to power level 2/10 as per the following sequence: 6 x (20s ON, 40s OFF) for a total of 2 minutes of sonication. The samples were centrifuged at 20,000 x g for 2 minutes and the supernatants were used for downstream proteomic analyses.

### Proteomics analysis

Extracted proteins were submitted to the Proteomics and Molecular Analysis Platform at the Research Institute of the McGill University Health Centre (RI-MUHC). For each sample, protein lysates were loaded onto a single stacking gel band to remove lipids, detergents and salts. The gel band was reduced with DTT, alkylated with iodoacetic acid and digested with trypsin. Extracted peptides were re-solubilized in 0.1% aqueous formic acid. LC-MS/MS analysis was performed on a Thermo FAIMS Duo Pro/Orbitrap Astral mass spectrometer system coupled to a Thermo Scientific Vanquish Neo UHPLC using mobile phase A: 0.1% formic acid in water and mobile phase B: 0.1% formic acid 80% acetonitrile in water. Briefly, 200 ng of peptide/sample were loaded onto a Thermo PEPMAP Neo 300 μm X 5 mm trap column into an IonOpticks Aurora Elite 150 μm X 7 cm separation column heated to 50 °C using a 60 SPD (20-minute gradient (4-40%) mobile phase B plus wash and reconditioning steps).

On the Orbitrap Astral spectrometer a FAIMS voltage of -45 V (to emphasize 2+ m/z) was used. MS1 spectra were recorded using the Orbitrap analyzer at a resolution of 240,000 from m/z 380 to 980 using an automated gate control (AGC) target of 500% and a maximum injection time of 3 ms. For MS2 in nDIA mode using the Astral analyzer, 200 nonoverlapping isolation windows from m/z 380 to 980 of 3 m/z were used. HCD collision energy is set to 25% and the RF lens at 40%.

DIA data were searched against a protein database including *H. hathewayi* (GCF_025149285.1) and *E. clostridioformis* YL32 (GCA_016696785.1) protein fasta files using Fragpipe v32.1 using the DIA-Speclib-Quant workflow using all nDIA files to create the library with quantification using DIA-NN v1.8.2beta8 functions in Fragpipe. Fragpipe modules used in the method are: MSFragger^75,76^; MSBooster^77^; Percolator^78^; ProteinProphet^79^; Philosopher^80^; Spectral library generation^81^; DIA quantification with DIA-NN^82^. The database search results were visualized and plotted using DIA-Analyst v0.11 (https://analyst-suites.org/apps/dia-analyst/) (Monash

Proteomics and Metabolomics Platform & Monash Bioinformatics Platform, Monash University) using the following parameters: Adjusted p-value: 0.05, Log2FC: 1, FDR correction: Benjamini Hochberg, Imputation Type: Perseus-type, Normalization: VSN normalization.

### Cah gene detection in stool samples

To detect *cah* genes in patient stool samples, PCR was performed on extracted fecal DNA using Q5 High-Fidelity 2X Master Mix (NEB #M0492) with *cah*-specific primers cahCD_gDNA_PCR_f and cahCD_gDNA_PCR_r [**Supplementary Table 5**]. 2 μL of fecal DNA was used as the PCR template and PCR was performed as follows: 30 s at 98 °C, 35 cycles (5 s at 98 °C, 30 s at 65 °C, 30 s at 72 °C), 2 min at 72 °C. Presence of the amplicon was visualized by 1% agarose gel electrophoresis and compared to a 1 kb Plus DNA Ladder (NEB #N0550). Premium PCR amplicon sequencing of selected samples was performed on PCR products by Plasmidsaurus using Oxford Nanopore Technology. Amplicon consensus sequences were used as queries to search for the best match to reference sequences from *Blautia* MSK.18.94, *H. hathewayi* and *M. formatexigens* using mmseqs2.

### Statistical analyses and graphing

Statistical methods were not used to determine sample sizes, experiments were not randomized, and the investigators were not blinded. Data points for in vitro and ex vivo assays were assumed to be normally distributed, though this was not formally tested. Data points for clinical trial analyses were assumed to be non-normally distributed and non-parametric tests were performed. Alpha and beta diversity analyses of metagenomics data were performed using the vegan R package (v2.7-2). Multivariable associations between species, number of reads, and patients were performed using the MaAsLin3 (v1.1.2) R package. Statistical tests used for comparisons between patient groups of study NCT04130321 are denoted in the Figure legends. Data were plotted in GraphPad Prism (v11.0.0) or ggplot2 (v4.0.1). Figures were assembled in Affinity Designer 2 (v2.6.3).

## Supplementary Figures

Supplementary Figure 1. Castalagin is metabolized to urolithin metabolites

Schematic for the breakdown of castalagin to downstream urolithin metabolites by gut bacteria. Castalagin is hydrolyzed and releases ellagic acid and castalin. Ellagic acid is subsequently decarboxylated to urolithin M-5 which undergoes dehydroxylation reactions to produce downstream urolithin metabolites. Terminal metabolite urolithin A, representing metabotype A is coloured in a blue box. Terminal metabolites isourolithin A and urolithin B, representing metabotype B are coloured in a green box. Metabolites commonly detected in metabolism assays are indicated in bold.

Supplementary Figure 2. Complex microbial communities convert castalagin to downstream metabolites

**a** (Left) Aligned chromatograms of CG-treated (100 uM) fecal slurries of samples BZ88, CI90, MO67 in mABB media after 0, 2, 4, 6, and 24 hours. (Right) Aligned chromatograms of CG- or H2O-treated media (mABB) and vehicle-treated stool slurries BZ88, CI90, MO67 in mABB media after 24 hours. **b** Aligned chromatograms of castalagin (CG), ellagic acid (EA), castalin (CL) and vehicle (V) treated fecal slurries of MO67 (blue) or media control (grey) after 4 hours. **c** MetaPhlAn analysis of DNA-sequencing from stool sample MO67. Relative abundance of phyla is represented on the left bar plot and families in the Firmicutes_A phyla are represented on the right bar plot. **d, e** MA plots of transcripts in MO67 fecal slurries treated with (**d**) castalin and (**e**) ellagic acid. (n = 4 biological replicates). Dots in blue represent significantly upregulated transcripts (Benjamini- Hochberg (BH) FDR < 0.05), grey dots include non-significant transcripts and those without an adjusted p-value.

Supplementary Figure 3. Multiple sequence alignment of Cah proteins

Protein matrices for **a** identity and **b** similarity of individual cah proteins in validated castalagin metabolizers *Blautia* MSK.94 (MSK18.94), *M. formatexigens* (Mf), *H. hathewayi* (Hh) and *E. clostridioformis* YL32 (Ec YL32) and upregulated transcript k117_197812. Genbank accessions used for comparisons are provided in **Supplementary Table 7**.

Supplementary Figure 4. cah-harbouring species convert castalagin and vescalagin to ellagic acid

**a** Chemical structure of castalagin or its C-1 epimer vescalagin. **b** Relative ellagic acid concentration measured in liquid cultures of Blautia MSK18.94 (MSK 18.94), H. hathewayi (Hh), M. formatexigens (Mf), B. glucerasea (Bg), B. wexlerae (Bw), E. clostridioformis (Ec), E. clostridioformis YL32 Ec YL32) and media control (BHI) incubated with castalagin (green) or vescalagin (blue) (n=3 biological replicates, error bars represent mean ± SEM).

Supplementary Figure 5. Substrate scope metabolism of punicalagin and corilagin.

**a** Chemical structure of punicalagin (left) and corilagin (right). **b** Aligned chromatograms of punicalagin (100 uM, blue), corilagin (100 uM, purple) and vehicle (H2O, grey) treated cultures of *H. hathewayi* (Hh), *M. formatexigens* (Mf), *Blautia* MSK18.94 (MSK), *E. clostridioformis* YL32 (Ec YL32) and uninoculated media control (BHI) after 48 hours. Chromatograms are representative of n=2-3 biological replicates.

Supplementary Figure 6. Castalagin is not converted to ellagic acid when *cah* is heterologously expressed in whole cells

Offset chromatograms of castalagin (CG, 100 uM, blue lines) or vehicle (H2O, gray lines) -treated whole cells of wild type (Cs_WT), pRG-MSK-cahABCD (Cs_cah 6-8) and pRG_eGFP (Cs_eGFP) C. sporogenes (Cs) cultures grown in BHI ± 30 μg/mL thiamphenicol. Retention times of the ellagic acid (EA) and internal standard (IS) are indicated by dotted lines. WT wild type; Cs_cah #: Cs pRG-MSK-cahABCD clone number.

Supplementary Figure 7. Cah proteins are upregulated in castalagin and castalin conditions in bacterial isolates

**a, b** Volcano plot of (**a**) castalagin (CG) and (**b**) castalin (CL) treated cultures of *H. hathewayi* and *E. clostridioformis* YL32 (n=3 biological replicates). Significantly upregulated proteins are indicated by red dots, significantly downregulated proteins are indicated by blue dots, Cah proteins are indicated by bright red dots and annotated. Log2FC cutoff = 1, adjusted p-value < 0.05 (Benjamini Hochberg correction). **c** Heat map of differentially expressed proteins in the CG, CL and Vehicle (H2O) treatment conditions. Hierarchical clustering is represented on both proteins (rows) and samples (columns). Significantly differentially expressed proteins in different compared conditions are indicated by orange boxes.

Supplementary Figure 8. Taxonomic distribution of cah and act genes in select genera

**a** Phylogenetic tree of species from *Hungatella* (dark green), *Marvinbryantia* (blue), *Enterocloster* (gray) and *Blautia* (light green) genera from GTDB r220 accessions. The outer rings represent the presence of homologous genes of the *cah* (green) or *act* (blue) gene clusters from *Blautia* MSK18.94. Presence is based on BLASTx search with 60% ID threshold. The number of genes represents how many genes from the individual gene clusters are present. Validated metabolizers are indicated by orange circles. **b** Heat map of individual *act* genes present in select *Lachnospiraceae* genomes from the NCBI based on BLASTx matches (% ID > 60; actA, actB1, actB2, actC1 and actC2 lengths > 200, actD length > 100, actE length > 300bp). Blue bars indicate genomes with a positive match.

Supplementary Figure 9. Genomes of cultured and uncultured Lachnospirales and Christensenellales encode *cah* cluster homologs

**a** Phylogenetic tree of the *Clostridia* class based on GTDB classification (release 232). Taxa of with >1 strain encoding 3/4 Cah proteins (hydrolases and oxidoreductase) with ≥ 70 %id to validated *cah*-encoded enzymes are annotated as solid lines colored in shades of yellow according to the level of castalagin metabolism validation. Taxa of with >1 strain encoding 3/3 Uxd or Ucd proteins with ≥ 70 %id to validated *uxd*- or *ucd*-encoded enzymes are annotated as dashed or solid black lines according to the level of urolithin dehydroxylation validation. The *Christensenellales* and *Lachnospirales* orders are represented by outer rings coloured in shades of grey. Taxa validated through metabolism assays are bolded. **b** Table of Cah proteins detected in the genome searches from taxa with >1 strain encoding 3/4 Cah proteins (hydrolases and oxidoreductase) with ≥ 70 %id to validated *cah*-encoded enzymes. Underscores represent an undetected Cah homologue.

Supplementary Figure 10. Cah operon organization varies between species

**a** Forward and reverse primer design for the RT-PCR experiment targeting the *cah* gene cluster in *Blautia* MSK18.94. **b** Agarose gel image of the RT-PCR amplicons (∼4.1 kb) from *Blautia* MSK18.94 *cah* cluster in cDNA, no reverse transcriptase (No RT), genomic DNA (gDNA) and no template control (NTC) samples induced with (CG) or uninduced (H2O). A ∼4.1 kb band represents the amplicon that covers the length of the *cah* cluster. **c** Genomic context of *H. hathewayi* and *E. clostridioformis* YL32, highlighting the mobile genetic elements and regulators interspersed with and around cah genes (green) and act genes (blue). * Indicates a gene annotated as a pseudogene.

Supplementary Figure 11. Defined minimal communities dictate castalagin metabolism to urolithin A

**a** Schematic demonstrating the cross-feeding of castalagin metabolites. Select cah- harbouring *Lachnospiraceae* convert castalagin to ellagic acid (EA) and castalin (CL). eam-harbouring *Gordonibacter* spp. and *Ellagibacter* spp. metabolize ellagic acid to urolithins. Urolithin C can be further converted to terminal metabolite urolithin A by *Enterocloster* spp. via the *uxd* and *ucd* operons. **b** Aligned chromatograms (λ = 305 nm) of castalagin (CG) or vehicle (H2O) treated cultures of *H. hathewayi* (Hh), Hh + *G. urolithinfaciens* (Gu), Gu + *E. bolteae* (Eb), Gu + *E. asparagiformis* (Ea), Hh + Gu, + Eb and Hh + Gu, + Ea after 5 days. c Aligned chromatograms of castalagin (CG) or vehicle (H2O) treated cultures of Blautia MSK18.94 (MSK), MSK + Gu, Gu + Eb and MSK + Gu, + Eb after 5 days.

Supplementary Figure 12. Metabolite profiles vary among individuals from a patient cohort

**a** Heat map representing metabolite abundance (based on relative peak area) per patient pre- (left) and post- (right) camu camu (CC) supplementation. Ellagic acid (EA) is representated by the green scale. Urolithins and flavogallonic acid (FGA) are represented by the blue scale. All metbaolite abundances are normalized to their 100uM analytical standard. **b** Relative metabolite abundance for each patient before and after CC supplementation, for non-signficant changes in metabolites UM6, UM7, uroB and isoA. Analyses were performed using two-tailed paired wilcoxon signed rank test with Pratt’s method and Holm-Sidak multiple comparison correction. **c** Correlation analysis between change in percent (%) hepatic fat fraction (HFF) and metabolite abundance for all patients (n = 20). Analyses performed using two-tailed Spearman rho correlation with Benjamini- Hochberg (BH) multiple comparison test. Spearman rho (*ρ*), p-values (p) and FDR (padj) values are indicated on each graph.

Supplementary Figure 13. Microbiome composition is not altered in patient samples post CC supplementation

**a** Taxonomic composition of patient fecal samples at the phylum (top) and species (bottom) level, pre- (left) and post- (right) CC supplementation. Species in the bottom plot represent taxa of interest in the metabolism of castalagin to terminal urolithins. Bars represent the relative abundance (%) of taxa per patient. **b** Alpha diversity per patient pre- (blue dots) and post-CC (red dots) supplementation represented by Shannon (left) and Simpson (right) diversity indices. Data are represented as individual points matched by patient; Wilcoxon singed rank exact test (matching by patient). Statistical significance is denoted above the plot with P-values. **c** Beta diversity analyses per patient pre- (blue dots) and post-CC (red dots) supplementation. Principle coordinate analyses are represented by Bray-Curtis, Jaccard and Robust Aitchison distances; PERMANOVA with 999 permutations. **d** Default MaAsLin3 output of species-level differences in abundance and prevalence pre- and post-CC (treatment post), accounting for the total number of reads in each sample (as a covariate). **e** MaAsLin3 output of species-level differences for selected taxa of interest involved in the metabolism of castalagin to terminal urolithins.

Supplementary Figure 14. C*a*h genes are widely detected in patient fecal DNA samples

**a** Schematic of the cah specific primers designed targeting conserved nucleotide regions in the cahC and cahD genes of *Blautia* MSK18.94, *Mf* and *Hh*. **b** PCR products from the *cahC-D* amplification run on a 1% agarose gel. **c** Best match of sequenced amplicons to reference genes. Columns represent the patient sample, best match to the species reference amplicon, percent identity and amplicon alignment length. Hh: *H. hathewayi*, Mf: *M. formatexigens*.

Supplementary Figure 15. Cah homologs are detected in patient metagenomes

**a, b** Prevalence of species and genes in patient stool metagenomes post-CC treatment including (**a**) extended *cah* spp. including Blautia_A faecis and Enterocloster clostridioformis and (**b**) Prevalence of extended *eam*, *uxd* and *ucd* spp. including QAMH01 sp900544245 (*eam*), Ventricola sp900542445 (*uxd* and *ucd*). Extended spp. are an extension of species indicated in **Figure 4**. Presence (gray) or absence (white) of targeted metabolites ellagic acid (EA), urolithin M5 (uroM5), urolithin M6 (uroM6), urolithin M7 (uroM7), urolithin C (uroC), urolithin D (uroD), urolithin A (uroA), urolithin B (uroB), isourolithin A (isoA) and flavogallonic acid (FGA) are indicated per patient.

Supplementary Figure 16. Genomes of cultured and uncultured Coriobacteriales encode eam cluster protein homologues.

**a** Phylogenetic tree of the *Coriobacteriales* order based on GTDB classification (release 232). Taxa of with >1 strain encoding Eah (hydrolase/decarboxylase) and ≥ 1 Eadh (dehydroxylase) with ≥ 70 %id to validated eam-encoded enzymes are annotated in shades of blue according to the level of EA metabolism validation. The QAMH01 and *Eggerthellaceae* families are represented by outer rings coloured in shades of grey. **b** Table of host(s) or sampling sources for taxa annotated in (**a**) based on NCBI metadata. Source data for (**a**) are provided as a Source data file.

Supplementary Figure 17. Eah homologues from *Coriobacteriales* metabolize ellagic acid

**a** Genomic context of known EA metabolizers, *G. urolithinfaciens* and *E. isourolithinifaciens*, with annotated genomic regions of CAAEEV01 sp011038975 and *Adlercreutzia murintestinalis*. Alignments were performed using the clinker tool with genome accessions in **Supplementary Table 7**. Protein sequences with identities ≥50% are connected by lines colored according to the percent identity value. The strain identity for each contig are denoted below the contig number. **b** Schematic representation of the conjugation and heterologous expression workflow to validate Eah homologues in *C. sporogenes* (*Cs*) whole cell assays. The generic pRG plasmid map is labeled with the origin of replication (ori), chloramphenicol acetyltransferase (*cat*), and promoter of the *Cs* ferredoxin gene with riboswitch G (Pfdx-rbG). **c** Aligned selected ion monitoring (SIM) chromatograms of substrates and metabolites from mid-exponential phase *Cs* cultures treated with 100 μM of EA or DMSO. Tested conditions included sterile media or *Cs* transconjugants expressing enhanced green fluorescent protein (eGFP) or *G. urolithinfaciens* Eah (Gu_Eah). **d** Aligned SIM chromatograms following the same experimental design as (**c**) using *Cs* transconjugants expressing enhanced green fluorescent protein (eGFP),CAAEV01 sp011038975 sp. ZJ176 Eah (ZJ176_Eah), or A. murintestinalis Eah (*Am*_Eah). For **c**-**d**, retention times of EA and urolithin standards are shown with grey dotted lines. Data from n = 1 of 3 biological replicates for each transconjugant are shown.

## Data availability

Unassembled sequence data (reads) for metatranscriptomics (RNA-sequencing) and metagenomics (DNA-sequencing) of the MO67 fecal sample were deposited in the NCBI Sequence Read Archive (SRA) under BioProject ID PRJNA1499868. The mass spectrometry proteomics data have been deposited to the ProteomeXchange Consortium via the PRIDE partner repository with the dataset identifier PXD083575. DNA sequences of plasmids generated in this study were deposited in GenBank: pRG-MSK18.94- CahABCD_HF1_1 (PZ898159), pRG-Guro-Eah_HF1_1 (PZ898160), pRG-DSM112345- Eah_HF2_1 (PZ898161), pRG-ZJ176-Eah_HF1_1 (PZ898162), pRG-k141_80263_7- Eah_HF1_2 (PZ898163), pRG-MGYG1194_200-Eah_HF2_1 (PZ898164), pRG- k141_58818-ucdCFO_HF1_1 (PZ898165), pRG-k141_58818-uxdFOC_HF2_1 (PZ898166). The pRG-eGFP plasmid is available in GenBank under accession PX095337.1.

Unassembled sequence data (reads) from clinical trial NCT04130321 were retrieved from the European Nucleotide Archive (ENA) under accession PRJEB76307. The Unified Human Gastrointestinal Proteome (UHGP) Human Gut v2.0.2 catalogue was retrieved from MGnify via the EMBL-EBI <u>FTP site</u>. Genomes of bacteria were retrieved from the NCBI FTP site according to genome accessions from the GTDB.

Contigs and datasets from metatranscriptomics and metagenomics assemblies were deposited along with code used in this study in Zenodo under DOI 10.5281/zenodo.23018167 and 10.5281/zenodo.23169451. Source data are provided in this paper.

## Code availability

Scripts used for the analysis of metagenomics data were deposited in GitHub [https://github.com/reillypidgeon/castalagin_stool_metagenomics] and Zenodo under DOI 10.5281/zenodo.23018167. Scripts used to determine protein-coding sequence presence in bacterial genomes (Bacteria Genome Mining) were deposited in GitHub [https://github.com/reillypidgeon/bacteria_genome_mining] and Zenodo under DOI 10.5281/zenodo.22900171.

## Acknowledgements

This research was funded by Weston Family Microbiome Initiative’s 2021 Transformational Research grant to B.C. and A.M. and a and a Weston Family Foundation 2024 Proof-of-Principle grant to B.C. and A.M.. B.C. held a tier II Canada Research Chair (CRC) in Therapeutic Chemistry (2015-2025). A.G. is supported by the FRQS Doctoral Research Scholarship (#370614, 2025-2029). R.P. was supported by the CIHR Canada Graduate Scholarship-Doctoral (#493808 for 2023-2026).

Metatranscriptomics and metagenomics library preparation and sequencing were performed by Génome Québec. Untargeted proteomics sample preparation and analysis was performed by the Research Institute of the McGill University Health Center (RI- MUHC) Proteomics and Molecular Analysis Platform. Metabolomics was performed by the McGill Metabolomics Innovation Resource (MIR) Platform. This research was enabled in part by support provided by Calcul Québec (calculquebec.ca) and the Digital Research Alliance of Canada (alliancecan.ca).

## Contributions

A.G. designed the study, performed experiments, analyzed data, created figures, and wrote the initial manuscript with B.C. R.P. performed experiments, performed bioinformatics analyses, analyzed data, and created figures. E.G. performed bioinformatics analyses on sequencing data. E.T. performed microbiology experiments and analyzed data. L.D. consulted on experimental design and methodology. C.F.M. supervised research, obtained ethical approval for the use of human fecal samples. A.M. supervised research, obtained ethical approval for the use of human fecal samples. B.C. designed the study, supervised the research, obtained research funding, and wrote the initial manuscript with A.G. All authors reviewed and edited the manuscript.

## Supplementary Tables

Supplementary Table 1. NCBI vs GTDB release taxonomy.

**Table 1.**
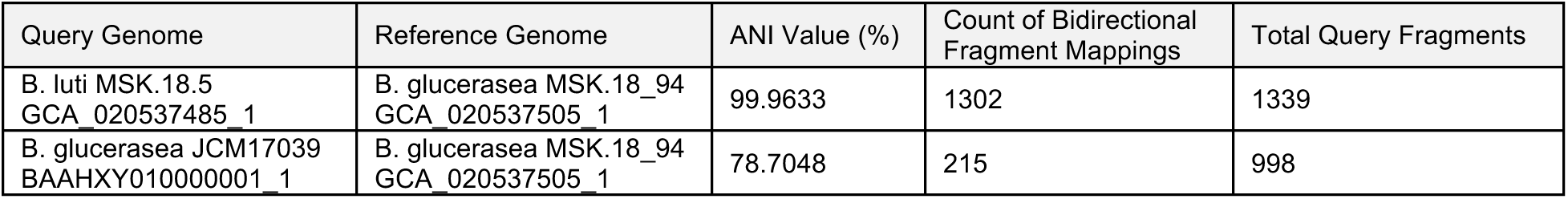
ANI analysis for Blautia_A sp905206915 against the type strain.

Supplementary Table 2. General reagents and materials.

**Table 2.** DNA-DNA hybridization analysis.

|  | **Sum of all HSP identities/overall HSP length** |  |  |  |  |
| **Reference Genome** | **Prob. DDH >= 70%** | **DDH** | **Model C.I.** | **G+C difference** | **GTDB Species Name** |
|---|---|---|---|---|---|
| Blautia glucerasea JCM 17039 | 0 | 23.8 | [21.5 - 26.2%] | 1.29 | NA |
| Blautia luti JCM 17040 | 0.1 | 29.8 | [27.4 - 32.3%] | 0.75 | s__Blautia_A luti |
| Blautia glucerasea MSK20.42 | 1.3 | 37.2 | [34.8 - 39.8%] | 0.97 | s__Blautia_A faecis |
| Blautia glucerasea MSK.21.93 | 0.26 | 32.3 | [29.9 - 34.8%] | 0.67 | s__Blautia_A ammoniilytica |
| Blautia glucerasea MSK.14.58 | 0.1 | 29.8 | [27.4 - 32.3%] | 0.93 | s__Blautia_A acetigignens |
| Blautia glucerasea MSK.18.95 | 1.51 | 37.8 | [35.3 - 40.3%] | 1.18 | s__Blautia_A faecis |
| Blautia glucerasea DFI.5.14 | 0.06 | 28.6 | [26.2 - 31.1%] | 1.39 | s__Blautia_A sp900120195 |
| Blautia glucerasea DFI.3.54 | 0.14 | 30.6 | [28.2 - 33.1%] | 0.88 | s__Blautia_A sp900548245 |
| Blautia glucerasea SUG1108 | 0 | 22.7 | [20.5 - 25.2%] | 0.08 | s__Blautia_A ammoniilytica |
| Blautia glucerasea hylon_METABAT__127 | 0 | 23.7 | [21.4 - 26.2%] | 0.32 | s__Blautia_A ammoniilytica |
| Blautia glucerasea LFYP119 | 0 | 22.4 | [20.1 - 24.8%] | 0.72 | NA |
| Blautia luti MSK.18.5 | 98.23 | 99.8 | [99.6 - 99.9%] | 0.02 | s__Blautia_A sp905206915 |
| Blautia glucerasea MSK.18.94 | 98.3 | 100 | [100 - NaN%] | 0 | s__Blautia_A sp905206915 |

Supplementary Table 3. Bacteria used in this study.

Supplementary Table 4. Modified anaerobe basal broth recipe.

Supplementary Table 5. Primers and gene fragments used in this study.

Supplementary Table 6. Human donor stool sample metadata.

Supplementary Table 7. Genome accessions for comparative genomics.

Supplementary Table 8. PCR conditions.

Supplementary Table 9. HiFi DNA assembly conditions for cloning.

Supplementary Table 10. Plasmids used in this study.

## Supplementary Notes

## Supplementary Discussion 1

In identifying the closest representative species to the MO67 *act* and *cah* genes, we identified *Blautia glucerasea* MSK.18.94 and *Blautia luti* MSK.18.5 as the species with the closest percent identities. Of note, both these species are classified as Blautia_A sp905206915 in the Genome Taxonomy Database R232. As such we proceeded with *Blautia* MSK.18.94 as the representative species in this work. Additionally, we have noted that the genome for *Blautia luti* MSK.18.95 (GCF_020537485.1) has recently been suppressed in the NCBI due to it being an unverified source organism.

Although classified as distinct species in the NCBI database, we hypothesize that these species represent different strains of the same species. Average nucleotide identity (ANI) measurement between the assembly of MSK.18.5 (GCA_020537485_1_ASM2053748v1) and MSK.18.94

(GCA_020537505_1_ASM2053750v1) was calculated to be 99.96%, suggesting that these two entries belong to the same species^83^.

The NCBI matches *Blautia glucerasea* as the best match type-strain for *B. glucerasea* MSK.18.94, with an ANI value of 83.75%, a value below the threshold of the genomes likely belonging to the same species, but instead likely belonging to a closely related species within the same genus. Additionally, the ANI calculation between *Blautia* MSK.18.94 and the *B. glucerasea* type strain (Table 1) was 78.7%, further suggesting that this bacterium belongs to the same genus but not the same species. It must be noted that the genome of the *B. glucerasea* type strain is not complete, and as such may account for the disparity between the NCBI ANI value and the value generated using FastANI. However, a genome-to-genome distance calculation analysis done on the available *Blautia glucerasea* genomes available in the NCBI database returned low probabilities that DNA-DNA hybridization (DDH) values are greater than 70% (0-1.51%) for all available *Blautia glucerasea* genomes, including the sequences available for the type strain (**Table 2**). As a positive control the probability that the DDH values for *Blautia* MSK.18.5 and MSK.18.94 were greater than 70% were 98.23% and 98.3% respectively (**Table 2**).

These values further support that this strain belongs a different species within the same genus. The DSMZ Type Strain Genome Server also identifies *Blautia glucerasea* MSK.18.94 as a new species among the *Blautia* genus, based on the available type strains in the DSMZ.

## References

1 Ferretti, P., Johnson, K., Priya, S. & Blekhman, R. Genomics of host–microbiome interactions in humans. Nature Reviews Genetics (2025). 10.1038/s41576-025-00849-8

2 Huttenhower, C. et al. Structure, function and diversity of the healthy human microbiome. Nature 486, 207–214 (2012). 10.1038/nature11234

3 Taguer, M. & Maurice, C. The complex interplay of diet, xenobiotics, and microbial metabolism in the gut: Implications for clinical outcomes. Clinical Pharmacology & Therapeutics 99, 588–599 (2016). 10.1002/cpt.366

4 Almeida, A. et al. A unified catalog of 204,938 reference genomes from the human gut microbiome. Nature Biotechnology 39, 105–114 (2021). 10.1038/s41587-020-0603-3

5 Zhang, Y. et al. Predicting functions of uncharacterized gene products from microbial communities. Nature Biotechnology (2025). 10.1038/s41587-025-02813-7

6 Hall, B. et al. BilR is a gut microbial enzyme that reduces bilirubin to urobilinogen. Nature Microbiology 9, 173–184 (2024). 10.1038/s41564-023-01549-x

7 Funabashi, M. et al. A metabolic pathway for bile acid dehydroxylation by the gut microbiome. Nature 582, 566–570 (2020). 10.1038/s41586-020-2396-4

8 Spanogiannopoulos, P., Bess, E. N., Carmody, R. N. & Turnbaugh, P. J. The microbial pharmacists within us: a metagenomic view of xenobiotic metabolism. Nature Reviews Microbiology 14, 273–287 (2016). 10.1038/nrmicro.2016.17

9 Guo, F., Bai, Y. & Lei, X. Gut microbiota alters drug metabolism and therapeutic outcomes. Cell Host & Microbe 34, 1262–1272 (2026). 10.1016/j.chom.2026.06.001

10 Koppel, N., Maini Rekdal, V. & Balskus, E. P. Chemical transformation of xenobiotics by the human gut microbiota. Science 356, eaag2770 (2017). doi:10.1126/science.aag2770

11 Ross, F. C. et al. The interplay between diet and the gut microbiome: implications for health and disease. Nature Reviews Microbiology (2024). 10.1038/s41579-024-01068-4

12. El-Saadony, M. T., et al. Polyphenols: Chemistry, bioavailability, bioactivity, nutritional aspects and human health benefits: A review. International Journal of Biological Macromolecules 277, 134223 (2024). 10.1016/j.ijbiomac.2024.134223

13 García-Villalba, R. et al. Urolithins: a Comprehensive Update on their Metabolism, Bioactivity, and Associated Gut Microbiota. Mol Nutr Food Res 66, e2101019 (2022). 10.1002/mnfr.202101019

14 Banc, R., Rusu, M. E., Filip, L. & Popa, D. S. The Impact of Ellagitannins and Their Metabolites through Gut Microbiome on the Gut Health and Brain Wellness within the Gut-Brain Axis. Foods 12 (2023). 10.3390/foods12020270

15 Messaoudene, M. et al. A Natural Polyphenol Exerts Antitumor Activity and Circumvents Anti-PD-1 Resistance through Effects on the Gut Microbiota. Cancer Discov 12, 1070–1087 (2022). 10.1158/2159-8290.Cd-21-0808

16 Mateș, L., Banc, R., Zaharie, F. A., Rusu, M. E. & Popa, D. S. Mechanistic Insights into the Biological Effects and Antioxidant Activity of Walnut (Juglans regia L.) Ellagitannins: A Systematic Review. Antioxidants (Basel) 13 (2024). 10.3390/antiox13080974

17 Medic, A., Jakopic, J., Hudina, M., Solar, A. & Veberic, R. Identification and quantification of the major phenolic constituents in Juglans regia L. peeled kernels and pellicles, using HPLC–MS/MS. Food Chemistry 352, 129404 (2021). 10.1016/j.foodchem.2021.129404

18 Quideau, S. et al. The chemistry of wine polyphenolic C-glycosidic ellagitannins targeting human topoisomerase II. Chemistry 11, 6503–6513 (2005). 10.1002/chem.200500428

19 Agrinier, A.-L. et al. Camu-camu decreases hepatic steatosis and liver injury markers in overweight, hypertriglyceridemic individuals: A randomized crossover trial. Cell Reports Medicine 5, 101682 (2024). 10.1016/j.xcrm.2024.101682

20 Anhê, F. F. et al. Treatment with camu camu (Myrciaria dubia) prevents obesity by altering the gut microbiota and increasing energy expenditure in diet- induced obese mice. Gut 68, 453–464 (2019). 10.1136/gutjnl-2017-315565

21 Pang, S. A., Elkrief, A., Capella, M. P. & Miller, W. H., Jr. Two Cases of Durable and Deep Responses to Immune Checkpoint Inhibition-Refractory Metastatic Melanoma after Addition of Camu Camu Prebiotic. Curr Oncol 30, 7852–7859 (2023). 10.3390/curroncol30090570

22 Elkrief, A. et al. The gut microbiome as a target in cancer immunotherapy: opportunities and challenges for drug development. Nature Reviews Drug Discovery (2025). 10.1038/s41573-025-01211-7

23 Piazza, S. et al. Investigation into the Anti-Acne Effects of Castanea sativa Mill Leaf and Its Pure Ellagitannin Castalagin in HaCaT Cells Infected with Cutibacterium acnes. International Journal of Molecular Sciences 25, 4764 (2024).

24 Iwatake, M., Okamoto, K., Tanaka, T. & Tsukuba, T. Castalagin Exerts Inhibitory Effects on Osteoclastogenesis Through Blocking a Broad Range of Signaling Pathways with Low Cytotoxicity. Phytotherapy Research 29, 917–924 (2015). 10.1002/ptr.5333

25 Iglesias-Aguirre, C. E. et al. Gut Bacteria Involved in Ellagic Acid Metabolism To Yield Human Urolithin Metabotypes Revealed. Journal of Agricultural and Food Chemistry 71, 4029–4035 (2023).

26 Pidgeon, R., Giurleo, A., Dridi, L. & Castagner, B. The presence and induction of regioselective dehydroxylases dictate urolithin metabolism by Enterocloster species. npj Biofilms and Microbiomes **11**, 220 (2025). 10.1038/s41522-025-00849-w

27 García-Villalba, R. o., Selma, M. a. V., Espín, J. C. & Tomás-Barberán, F. A. Identification of Novel Urolithin Metabolites in Human Feces and Urine after the Intake of a Pomegranate Extract. Journal of Agricultural and Food Chemistry 67, 11099–11107 (2019). 10.1021/acs.jafc.9b04435

28 Ryu, D. et al. Urolithin A induces mitophagy and prolongs lifespan in C. elegans and increases muscle function in rodents. Nature Medicine 22, 879–888 (2016). 10.1038/nm.4132

29 Cortés-Martín, A. et al. The gut microbiota urolithin metabotypes revisited: the human metabolism of ellagic acid is mainly determined by aging. Food & function 9, 4100–4106 (2018).

30 Li, Y. et al. Development of a (Poly)phenol Metabolic Signature for Assessing (Poly)phenol-Rich Dietary Patterns. Journal of Agricultural and Food Chemistry 72, 13439–13450 (2024). 10.1021/acs.jafc.4c00959

31 García-Villalba, R. o., et al. Gastrointestinal Simulation Model TWIN-SHIME Shows Differences between Human Urolithin-Metabotypes in Gut Microbiota Composition, Pomegranate Polyphenol Metabolism, and Transport along the Intestinal Tract. Journal of Agricultural and Food Chemistry 65, 5480–5493 (2017). 10.1021/acs.jafc.7b02049

32 Yamada, H., Wakamori, S., Hirokane, T., Ikeuchi, K. & Matsumoto, S. Structural Revisions in Natural Ellagitannins. Molecules 23, 1901 (2018).

33 Bae, M. et al. Distinct classes of gut bacterial molybdenum-dependent enzymes produce urolithins. Proceedings of the National Academy of Sciences 122, e2501312122 (2025). doi:10.1073/pnas.2501312122

34 Pidgeon, R. et al. Diet-derived urolithin A is produced by a dehydroxylase encoded by human gut Enterocloster species. Nature Communications 16, 999 (2025). 10.1038/s41467-025-56266-2

35 Pulido-Mateos, E. C., Lessard-Lord, J., Guyonnet, D., Desjardins, Y. & Roy, D. Comprehensive analysis of the metabolic and genomic features of tannin- transforming Lactiplantibacillus plantarum strains. Scientific Reports 12, 22406 (2022). 10.1038/s41598-022-26005-4

36 Pulido-Mateos, E. C., Lessard-Lord, J., Desjardins, Y. & Roy, D. Biotransformation of camu–camu galloylated ellagitannins by Lactiplantibacillus plantarum with extracellular tannase activity. Food & Function 15, 7189–7199 (2024). 10.1039/D4FO00149D

37 He, J. et al. Gut microbiome-derived hydrolases-an underrated target of natural product metabolism. Front Cell Infect Microbiol 14, 1392249 (2024). 10.3389/fcimb.2024.1392249

38 Engström, M. T. et al. The Oxidative Activity of Ellagitannins Dictates Their Tendency To Form Highly Stabilized Complexes with Bovine Serum Albumin at Increased pH. Journal of Agricultural and Food Chemistry 64, 8994–9003 (2016). 10.1021/acs.jafc.6b01571

39 Hofmann, T. et al. Protein binding and astringent taste of a polymeric procyanidin, 1,2,3,4,6-penta-O-galloyl-beta-D-glucopyranose, castalagin, and grandinin. J Agric Food Chem 54, 9503–9509 (2006). 10.1021/jf062272c

40 Love, M. I., Huber, W. & Anders, S. Moderated estimation of fold change and dispersion for RNA-seq data with DESeq2. Genome Biology 15, 550 (2014). 10.1186/s13059-014-0550-8

41 Cañadas, I. C., Groothuis, D., Zygouropoulou, M., Rodrigues, R. & Minton, N. P. RiboCas: A Universal CRISPR-Based Editing Tool for Clostridium. ACS Synthetic Biology 8, 1379–1390 (2019). 10.1021/acssynbio.9b00075

42 Dadi, T. H. et al. Lythrum salicaria L. herb and gut microbiota of healthy post- weaning piglets. Focus on prebiotic properties and formation of postbiotic metabolites in ex vivo cultures. Journal of Ethnopharmacology 261, 113073 (2020).

43 Njenga, R., Boele, J., Öztürk, Y. & Koch, H. G. Coping with stress: How bacteria fine-tune protein synthesis and protein transport. J Biol Chem 299, 105163 (2023). 10.1016/j.jbc.2023.105163

44 Osbourn, A. E. & Field, B. Operons. Cellular and Molecular Life Sciences 66, 3755–3775 (2009). 10.1007/s00018-009-0114-3

45 Bratlie, M. S., Johansen, J. & Drabløs, F. Relationship between operon preference and functional properties of persistent genes in bacterial genomes. BMC Genomics 11, 71 (2010). 10.1186/1471-2164-11-71

46 Moussa, M. R. et al. Systemic Inflammation and the Inflammatory Context of the Colonic Microenvironment Are Improved by Urolithin A. Cancer Prev Res (Phila) 18, 235–250 (2025). 10.1158/1940-6207.Capr-24-0383

47 Pfundstein, B. et al. Pilot Walnut Intervention Study of Urolithin Bioavailability in Human Volunteers. Journal of Agricultural and Food Chemistry 62, 10264–10273 (2014). 10.1021/jf5040652

48 Sivamani, R. K. et al. Prospective Randomized, Double-Blind, Placebo-Controlled Study of a Standardized Oral Pomegranate Extract on the Gut Microbiome and Short-Chain Fatty Acids. Foods 13 (2023). 10.3390/foods13010015

49 Anderson, B. D. & Bisanz, J. E. Challenges and opportunities of strain diversity in gut microbiome research. Frontiers in Microbiology **Volume** 14–2023 (2023). 10.3389/fmicb.2023.1117122

50 da Silva, A. C., Lapkin, J., Yin, Q., Muller, E. & Almeida, A. Meta-analysis of the uncultured gut microbiome across 11,115 global metagenomes reveals a candidate signature of health. Cell Host & Microbe 34, 379–392.e375 (2026). 10.1016/j.chom.2026.01.013

51 Peng, H. et al. Longitudinal gut microbiota tracking reveals the dynamics of horizontal gene transfer. Nature Communications 16, 11543 (2025). 10.1038/s41467-025-66612-z

52 Ardalani, O. et al. Annotating the pangenome reveals the diversity in the genetic basis for metabolic enzymes. Science Advances 12, eaeb3363 10.1126/sciadv.aeb3363

53 Levy, R. & Borenstein, E. Metabolic modeling of species interaction in the human microbiome elucidates community-level assembly rules. Proceedings of the National Academy of Sciences 110, 12804–12809 (2013). 10.1073/pnas.1300926110

54 Tian, L. et al. Deciphering functional redundancy in the human microbiome. Nature Communications 11, 6217 (2020). 10.1038/s41467-020-19940-1

55 Park, H., Kim, J. S., Kim, D. J. & Suk, K. T. Strain Diversity in the Human Microbiome: Personal Variation, Pathobionts, Therapeutics, and Methodological Challenges. Microorganisms 14, 720 (2026).

56 Singh, A. et al. Direct supplementation with Urolithin A overcomes limitations of dietary exposure and gut microbiome variability in healthy adults to achieve consistent levels across the population. European Journal of Clinical Nutrition 76, 297–308 (2022). 10.1038/s41430-021-00950-1

57 Andreux, P. A. et al. The mitophagy activator urolithin A is safe and induces a molecular signature of improved mitochondrial and cellular health in humans. Nature Metabolism 1, 595–603 (2019). 10.1038/s42255-019-0073-4

58 Fraga, C. G., Croft, K. D., Kennedy, D. O. & Tomás-Barberán, F. A. The effects of polyphenols and other bioactives on human health. Food & Function 10, 514–528 (2019). 10.1039/c8fo01997e

59 Rudrapal, M., de Oliveira, A. M. & Singh, R. P. Dietary polyphenols maintain human health through modulation of gut microbiota. Front Pharmacol 16, 1710088 (2025). 10.3389/fphar.2025.1710088

60 Bae, M. et al. Metatranscriptomics-guided discovery and characterization of a polyphenol-metabolizing gut microbial enzyme. Cell Host & Microbe 32, 1887–1896.e1888 (2024). 10.1016/j.chom.2024.10.002

61 Maini Rekdal, V., Bess, E. N., Bisanz, J. E., Turnbaugh, P. J. & Balskus, E. P. Discovery and inhibition of an interspecies gut bacterial pathway for Levodopa metabolism. Science 364, eaau6323 (2019). doi:10.1126/science.aau6323

62 Dong, X., Bae, M., Le, C., Aguilar Ramos, M. A. & Balskus, E. P. Enantiocomplementary Gut Bacterial Enzymes Metabolize Dietary Polyphenols. Journal of the American Chemical Society 147, 7231–7244 (2025). 10.1021/jacs.4c09892

63 Yang, G. et al. Discovery of an ene-reductase for initiating flavone and flavonol catabolism in gut bacteria. Nature Communications 12, 790 (2021). 10.1038/s41467-021-20974-2

64 FelixKrueger/TrimGalore: v0.6.10 - add default decompression path (Version 0.6.10) (Zenodo, 2023).

65 Martin, M. Cutadapt removes adapter sequences from high-throughput sequencing reads. EMBnet.journal 17, 10–12 (2011). doi10.14806/ej.17.1.200

66. Bushnell, B. in *9th Annual Genomics of Energy & Environment Meeting,*.

67 Kopylova, E., Noé, L. & Touzet, H. SortMeRNA: fast and accurate filtering of ribosomal RNAs in metatranscriptomic data. Bioinformatics 28, 3211–3217 (2012). 10.1093/bioinformatics/bts611

68 Durai, D. A. & Schulz, M. H. Improving in-silico normalization using read weights. Scientific Reports 9, 51 33 (2019). 10.1038/s41598-019-41502-9

69 Li, D., Liu, C.-M., Luo, R., Sadakane, K. & Lam, T.-W. MEGAHIT: an ultra-fast single-node solution for large and complex metagenomics assembly via succinct de Bruijn graph. Bioinformatics 31, 1674–1676 (2015). 10.1093/bioinformatics/btv033

70 Bray, N. L., Pimentel, H., Melsted, P. & Pachter, L. Near-optimal probabilistic RNA- seq quantification. Nature Biotechnology 34, 525–527 (2016). 10.1038/nbt.3519

71 Soneson, C., Love, M. I. & Robinson, M. D. Differential analyses for RNA-seq: transcript-level estimates improve gene-level inferences. F1000Res 4, 1521 (2015). 10.12688/f1000research.7563.2

72 Benjamini, Y. & Hochberg, Y. Controlling the False Discovery Rate: A Practical and Powerful Approach to Multiple Testing. Journal of the Royal Statistical Society: Series B (Methodological) 57, 289–300 (1995). 10.1111/j.2517-6161.1995.tb02031.x

73 van den Belt, M., et al. CAGECAT: The CompArative GEne Cluster Analysis Toolbox for rapid search and visualisation of homologous gene clusters. BMC Bioinformatics 24, 181 (2023). 10.1186/s12859-023-05311-2

74. Hahsler M, N. A. rBLAST: R Interface for the Basic Local Alignment Search Tool.” Bioconductor version: Release (3.19). R package version 0.99.4. (2024). 10.18129/B9.bioc.rBLAST

75. Kong, A. T., Leprevost, F. V., Avtonomov, D. M., Mellacheruvu, D. & Nesvizhskii, A. I. MSFragger: ultrafast and comprehensive peptide identification in mass spectrometry–based proteomics. Nature Methods 14, 513–520 (2017). 10.1038/nmeth.4256

76 Teo, G. C., Polasky, D. A., Yu, F. & Nesvizhskii, A. I. Fast Deisotoping Algorithm and Its Implementation in the MSFragger Search Engine. Journal of Proteome Research 20, 498–505 (2021). 10.1021/acs.jproteome.0c00544

77 Yang, K. L. et al. MSBooster: improving peptide identification rates using deep learning-based features. Nature Communications 14, 4539 (2023). 10.1038/s41467-023-40129-9

78 Käll, L., Canterbury, J. D., Weston, J., Noble, W. S. & MacCoss, M. J. Semi- supervised learning for peptide identification from shotgun proteomics datasets. Nature Methods 4, 923–925 (2007). 10.1038/nmeth1113

79 Nesvizhskii, A. I., Keller, A., Kolker, E. & Aebersold, R. A Statistical Model for Identifying Proteins by Tandem Mass Spectrometry. Analytical Chemistry 75, 4646–4658 (2003). 10.1021/ac0341261

80 da Veiga Leprevost, F., et al. Philosopher: a versatile toolkit for shotgun proteomics data analysis. Nature Methods 17, 869–870 (2020). 10.1038/s41592-020-0912-y

81 Yu, F. et al. Analysis of DIA proteomics data using MSFragger-DIA and FragPipe computational platform. Nature Communications 14, 4154 (2023). 10.1038/s41467-023-39869-5

82 Demichev, V. et al. dia-PASEF data analysis using FragPipe and DIA-NN for deep proteomics of low sample amounts. Nature Communications 13, 3944 (2022). 10.1038/s41467-022-31492-0

83 Jain, C., Rodriguez-R, L. M., Phillippy, A. M., Konstantinidis, K. T. & Aluru, S. High throughput ANI analysis of 90K prokaryotic genomes reveals clear species boundaries. Nature Communications 9, 5114 (2018). 10.1038/s41467-018-07641-9

