## Supplementary Figures for "A hydrolysis operon in *Lachnospiraceae* provides a genetic basis to understand castalagin metabolism by the human microbiome in a clinical study"

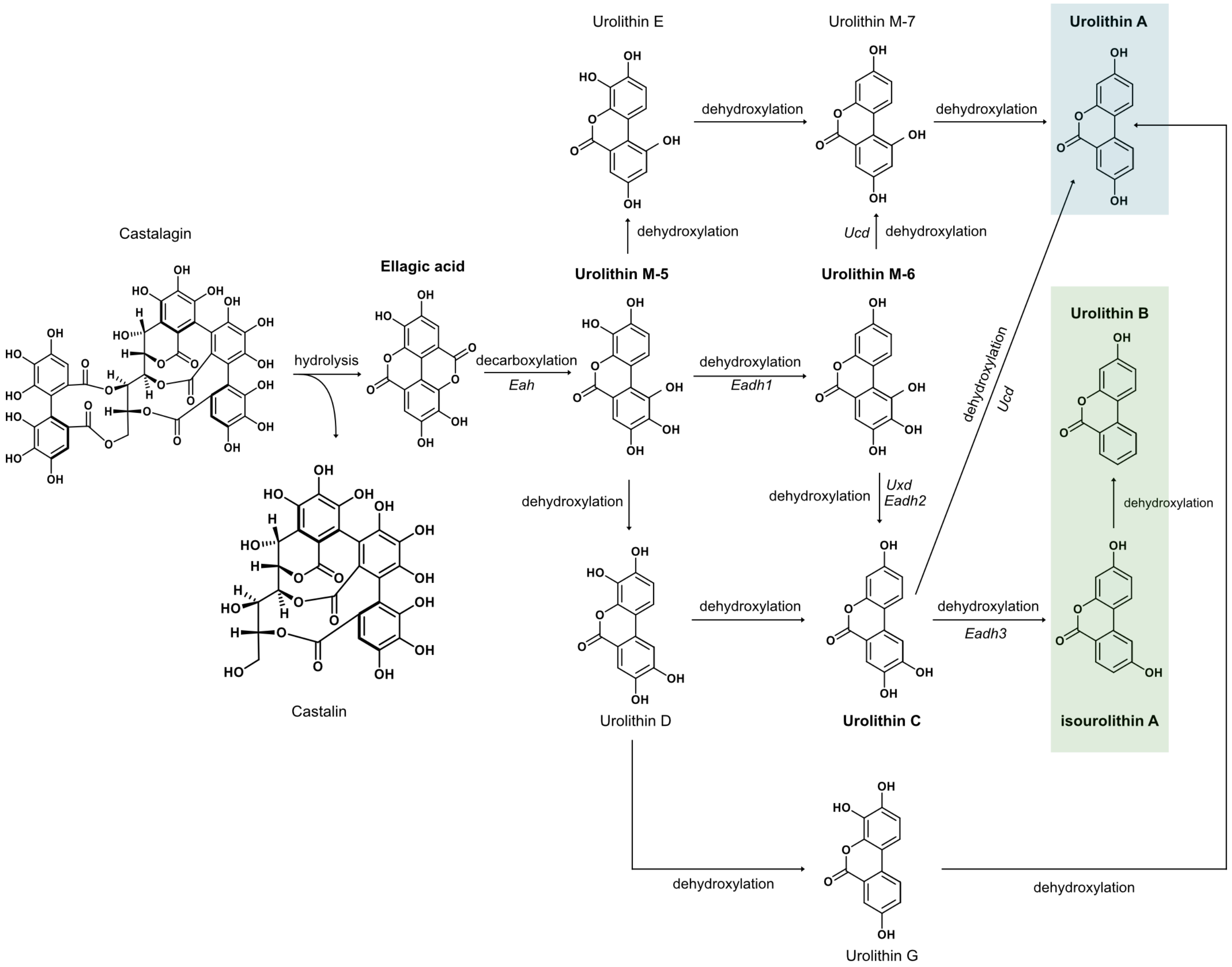


**Supplementary Figure 1.** Castalagin is metabolized to urolithin metabolites

Schematic for the breakdown of castalagin to downstream urolithin metabolites by gut bacteria. Castalagin is hydrolyzed and releases ellagic acid and castalin. Ellagic acid is subsequently decarboxylated to urolithin M-5 which undergoes dehydroxylation reactions to produce downstream urolithin metabolites. Terminal metabolite urolithin A, representing metabotype A is coloured in a blue box. Terminal metabolites isourolithin A and urolithin B, representing metabotype B are coloured in a green box. Metabolites commonly detected in metabolism assays are indicated in bold.

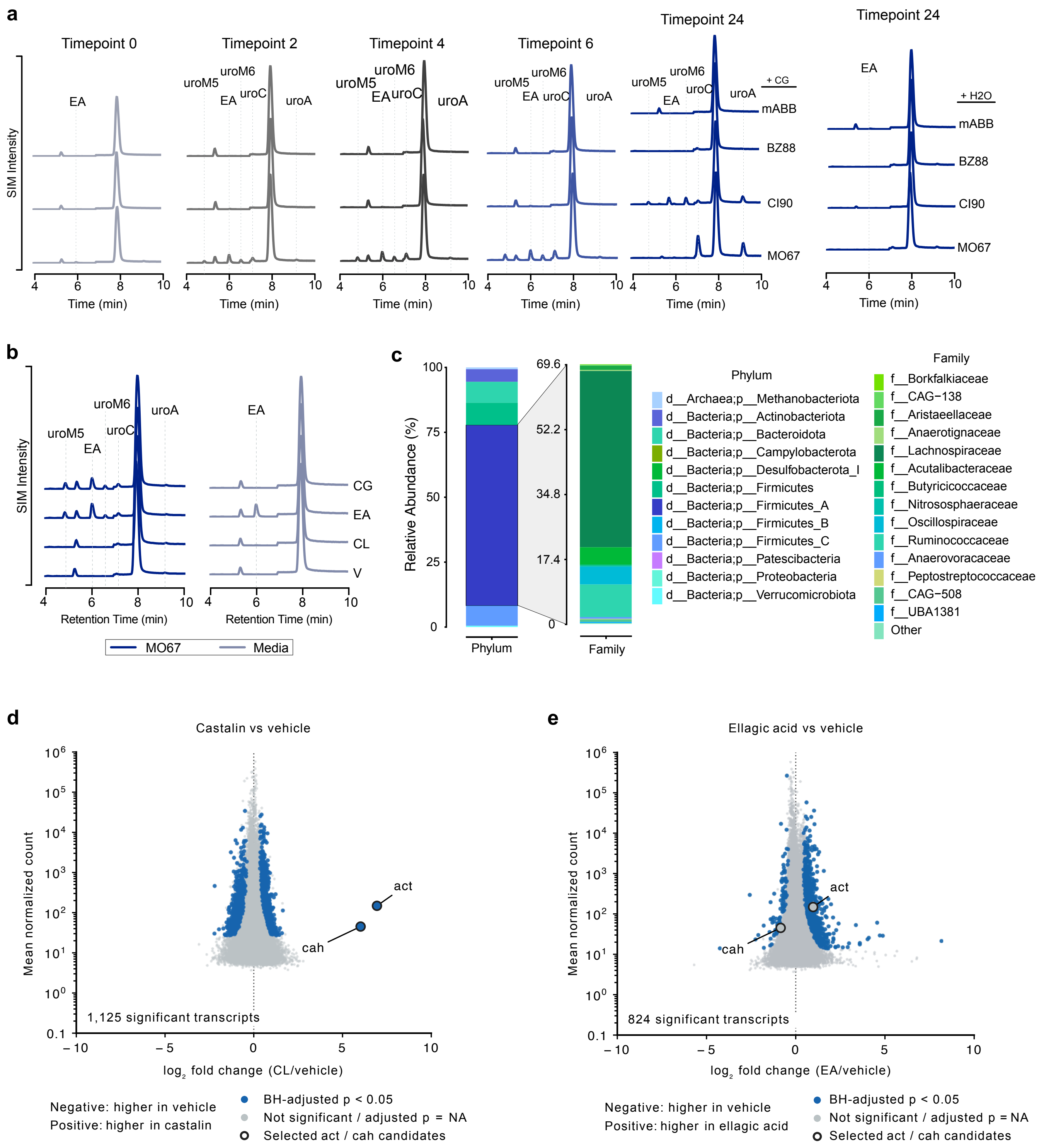


**Supplementary Figure 2.** Complex microbial communities convert castalagin to downstream metabolites

**a** (Left) Aligned chromatograms of CG-treated (100 uM) fecal slurries of samples BZ88, CI90, MO67 in mABB media after 0, 2, 4, 6, and 24 hours. (Right) Aligned chromatograms of CG- or H2O-treated media (mABB) and vehicle-treated stool slurries BZ88, CI90, MO67 in mABB media after 24 hours. **b** Aligned chromatograms of castalagin (CG), ellagic acid (EA), castalin (CL) and vehicle (V) treated fecal slurries of MO67 (blue) or media control (grey) after 4 hours. **c** MetaPhlAn analysis of DNA-sequencing from stool sample MO67. Relative abundance of phyla is represented on the left bar plot and families in the Firmicutes_A phyla are represented on the right bar plot. **d, e** MA plots of transcripts in MO67 fecal slurries treated with (**d**) castalin and (**e**) ellagic acid. (n = 4 biological replicates). Dots in blue represent significantly upregulated transcripts (Benjamini-Hochberg (BH) FDR < 0.05), grey dots include non-significant transcripts and those without an adjusted p-value.

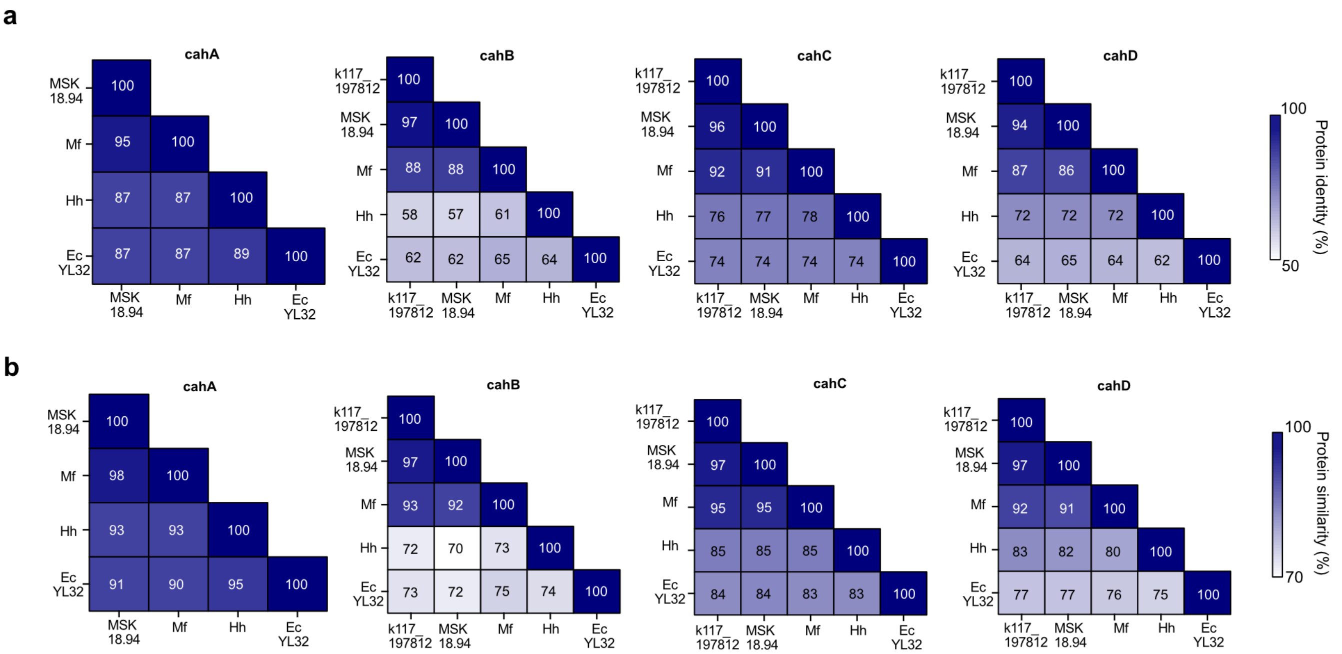


### **Supplementary Figure 3.** Multiple sequence alignment of Cah proteins

Protein matrices for **a** identity and **b** similarity of individual Cah proteins in validated castalagin metabolizers *Blautia* MSK.18.94, *M. formatexigens* (Mf), *H. hathewayi* (Hh) and *E. clostridioformis* YL32 (Ec YL32) and upregulated transcript k117_197812. Genbank accessions used for comparisons are provided in **Supplementary Table 7**.

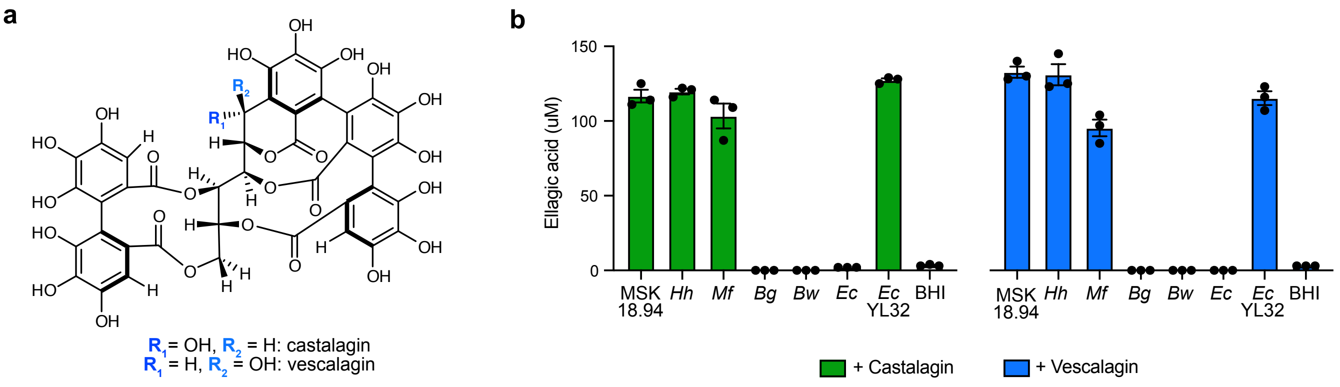


### **Supplementary Figure 4.** cah-harbounring species convert castalagin and vescalagin to ellagic acid

**a** Chemical structure of castalagin or its C-1 epimer vescalagin. **b** Relative ellagic acid concentration measured in liquid cultures of Blautia MSK18.94 (MSK 18.94), H. hathewayi (Hh), M. formatexigens (Mf), B. glucerasea (Bg), B. wexlerae (Bw), E. clostridioformis (Ec), E. clostridioformis YL32 Ec YL32) and media control (BHI) incubated with castalagin (green) or vescalagin (blue) (n=3 biological replicates, error bars represent mean ± SEM).

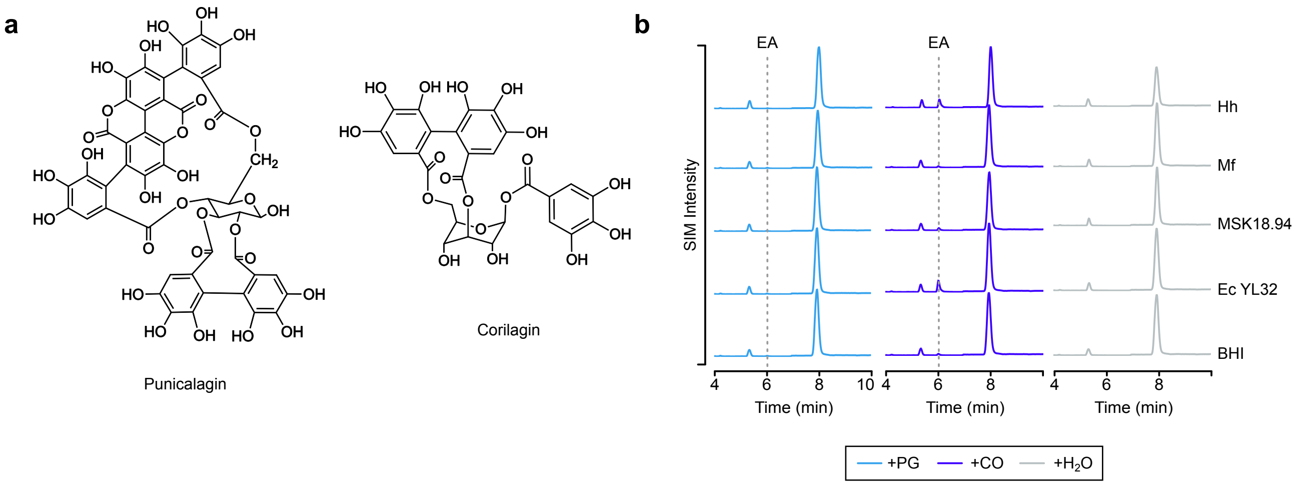


**Supplementary Figure 5.** Substrate scope metabolism of punicalagin and corilgain

**a** Chemical structures of punicalagin (left) and corilagin (right). **b** Aligned chromatograms of punicalagin (100 uM, blue), corilagin (100 uM, purple) and vehicle (H2O, grey) treated cultures of *H. hathewayi* (Hh), *M. formatexigens* (Mf), *Blautia* MSK18.94 (MSK), *E. clostridioformis* YL32 (Ec YL32) and uninoculated media control (BHI) after 48 hours. Chromatograms are representative of n=2-3 biological replicates.

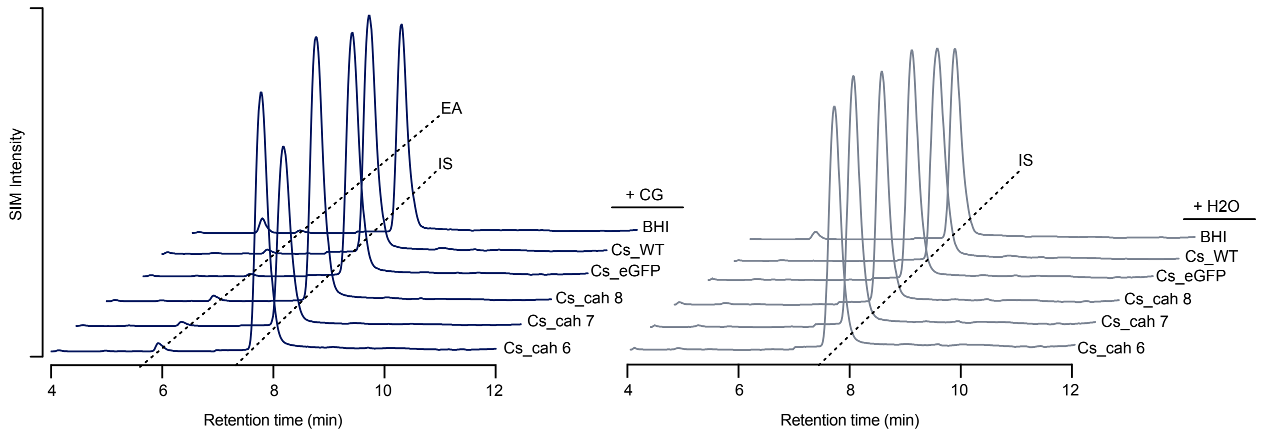


**Supplementary Figure 6.** Castalagin is not converted to ellagic acid when Cah is heterologously expressed in whole cells

Offset chromatograms of castalagin (CG, 100 uM, blue lines) or vehicle (H2O, gray lines) -treated whole cells of wild type (Cs_WT), pRG-MSK-cahABCD (Cs_cah 6-8) and pRG_eGFP (Cs_eGFP) C. sporogenes (Cs) cultures grown in BHI ± 30 μg/mL thiamphenicol. Retention times of the ellagic acid (EA) and internal standard (IS) are indicated by dotted lines. WT wild type; Cs_cah #: Cs pRG-MSK-cahABCD clone number.

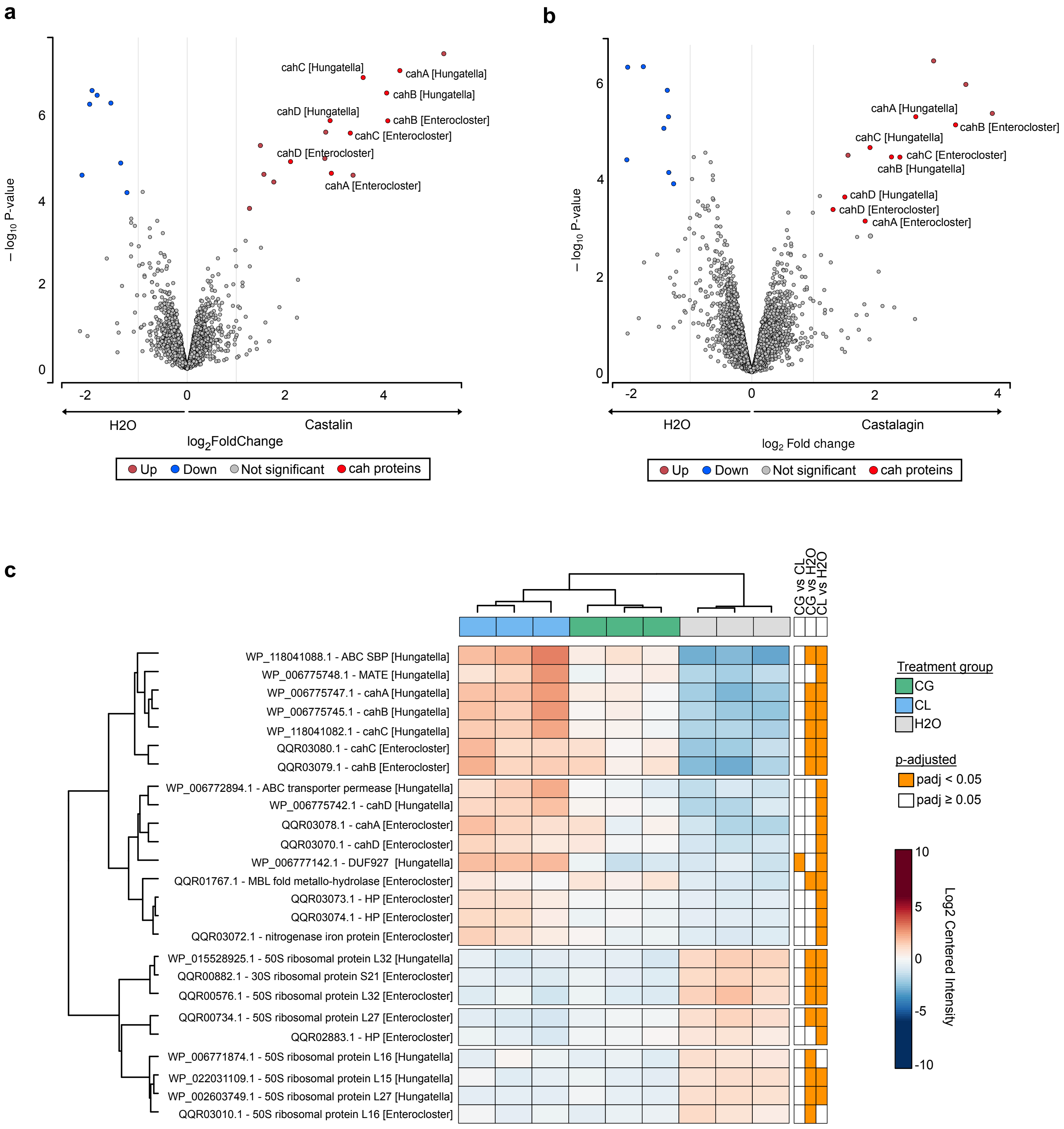


**Supplementary Figure 7.** Cah proteins are upregulated by CG and CL treatment in bacterial isolates

**a, b** Volcano plot of (**a**) castalagin (CG) and (**b**) castalin (CL) treated cultures of *H. hathewayi* and *E. clostridioformis* YL32 (n=3 biological replicates). Significantly upregulated proteins are indicated by red dots, significantly downregulated proteins are indicated by blue dots, Cah proteins are indicated by bright red dots and annotated. Log2FC cutoff = 1, adjusted p-value < 0.05 (Benjamini Hochberg correction). **c** Heat map of differentially expressed proteins in the CG, CL and Vehicle (H2O) treatment conditions. Hierarchical clustering is represented on both proteins (rows) and samples (columns). Significantly differentially expressed proteins in different compared conditions are indicated by orange boxes. SBP substrate binding protein, HP hypothetical protein, MATE multi-antimicrobial extrusion

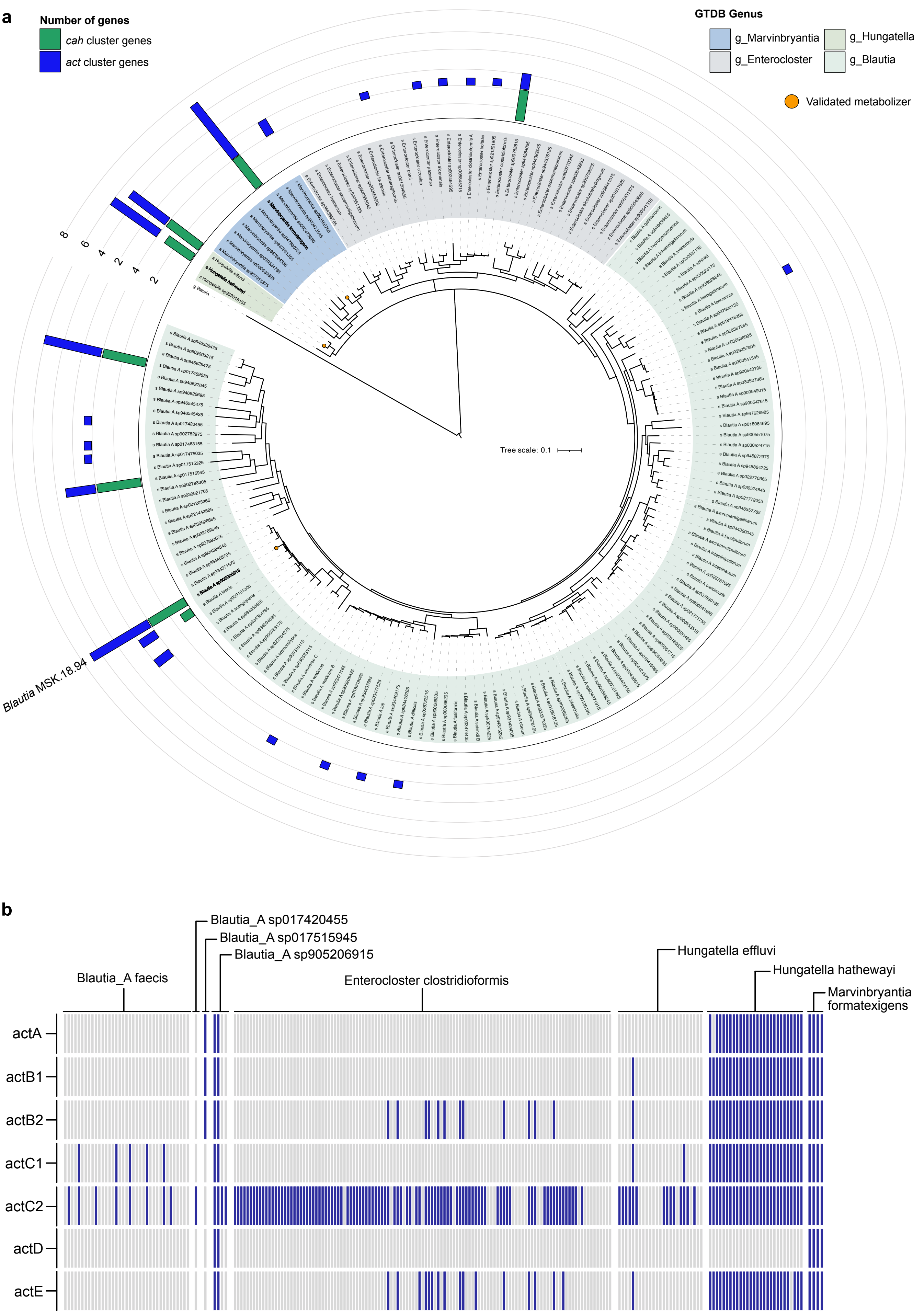
a
Number of genes
cah cluster genes
act cluster genes
GTDB Genus
g_Marvinbryantia
g_Enterocloster
g_Hungatella
g_Blautia
Validated metabolizer
8
6
4
2
4
2
Tree scale: 0.1
Blautia MSK.18.94
b
Blautia_A faecis
Blautia_A sp017420455
Blautia_A sp017515945
Blautia_A sp905206915
Enterocloster clostridioformis
Hungatella effluvi
Hungatella hathewayi
Marvinbryantia formatexigens
actA
actB1
actB2
actC1
actC2
actD
actE

**Supplementary Figure 8.** Taxonomic distribution of *cah* and *act* genes

**a** Phylogenetic tree of species from *Hungatella* (dark green), *Marvinbryantia* (blue), *Enterocloster* (gray) and *Blautia* (light green) genera from GTDB r220 accessions. The outer rings represent the presence of homologous genes of the *cah* (green) or *act* (blue) gene clusters from *Blautia* MSK18.94. Presence is based on BLASTx search with 60% ID threshold. The number of genes represents how many genes from the individual gene clusters are present. Validated metabolizers are indicated by orange circles. **b** Heat map of individual *act* genes present in select *Lachnospiraceae* genomes from the NCBI based on BLASTx matches (% ID > 60; actA, actB1, actB2, actC1 and actC2 lengths > 200, actD length > 100, actE length > 300bp). Blue bars indicate genomes with a positive match.

**a**

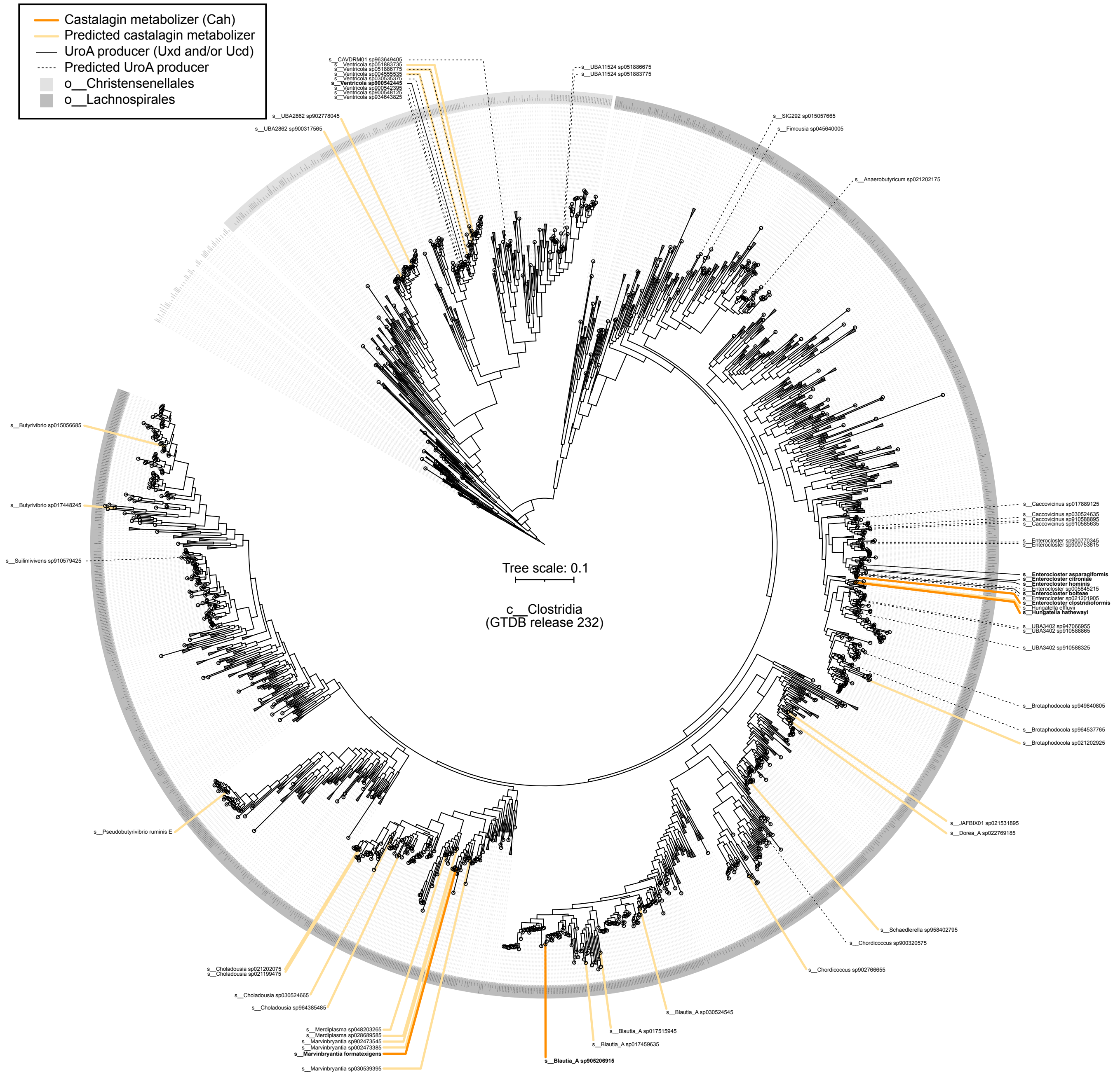


**b**

| GTDB species | Cah proteins | GTDB species | Cah proteins |
|---|---|---|---|
| s__UBA2862 sp900317565 | ABC_ | s__Blautia_A sp017459635 | ABCD |
| s__UBA2862 sp902778045 | ABC_ | **s__Blautia_A sp905206915** | ABCD |
| s__Ventricola sp004555535 | ABC_ | s__Marvinbryantia sp030539395 | ABCD |
| s__Ventricola sp051886775 | ABC_ | **s__Marvinbryantia formatexigens** | ABCD |
| s__Ventricola sp051883735 | ABC_ | s__Marvinbryantia sp002473385 | ABCD |
| **s__Enterocloster clostridioformis** | ABCD | s__Marvinbryantia sp902473545 | A_CD |
| s__Hungatella effluvii | ABCD | s__Merdiplasma sp028689585 | ABCD |
| **s__Hungatella hathewayi** | ABCD | s__Merdiplasma sp048203265 | ABCD |
| s__Brotaphodocola sp021202925 | ABCD | s__Choladousia sp964385485 | ABC_ |
| s__JAFBIX01 sp021531895 | _BCD | s__Choladousia sp030524665 | ABCD |
| s__Dorea_A sp022769185 | ABC_ | s__Choladousia sp021199475 | ABCD |
| s__Schaedlerella sp958402795 | ABC_ | s__Choladousia sp021202075 | ABCD |
| s__Chordicoccus sp902766655 | ABC_ | s__Pseudobutyrivibrio ruminis_E | ABC_ |
| s__Blautia_A sp030524545 | ABCD | s__Butyrivibrio sp017448245 | ABC_ |
| s__Blautia_A sp017515945 | ABCD | s__Butyrivibrio sp015056685 | ABC_ |

**Supplementary Figure 9.** Genomes of cultured and uncultured *Lachnospirales* and *Christensenellales* encode *cah* cluster homologs

**a** Phylogenetic tree of the *Clostridia* class based on GTDB classification (release 232). Taxa of with >1 strain encoding 3/4 Cah proteins (hydrolases and oxidoreductase) with ≥ 70 %id to validated *cah*-encoded enzymes are annotated as solid lines colored in shades of yellow according to the level of castalagin metabolism validation. Taxa of with >1 strain encoding 3/3 Uxd or Ucd proteins with ≥ 70 %id to validated *uxd*- or *ucd*-encoded enzymes are annotated as dashed or solid black lines according to the level of urolithin dehydroxylation validation. The *Christensenellales* and *Lachnospirales* orders are represented by outer rings coloured in shades of grey. Taxa validated through metabolism assays are bolded. **b** Table of Cah proteins detected in the genome searches from taxa with >1 strain encoding 3/4 Cah proteins (hydrolases and oxidoreductase) with ≥ 70 %id to validated *cah*-encoded enzymes. Underscores represent an undetected Cah homologue.

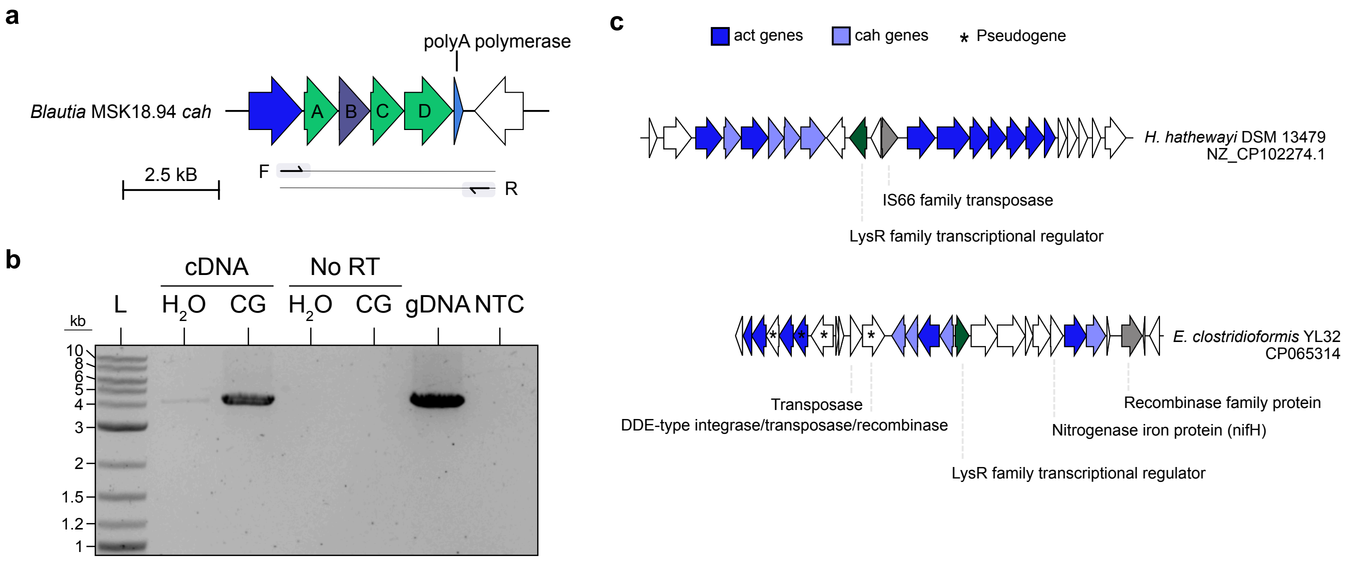


**Supplementary Figure 10.** cah operon organization varies between species

**a** Forward and reverse primer design for the RT-PCR experiment targeting the *cah* gene cluster in *Blautia* MSK18.94. **b** Agarose gel image of the RT-PCR amplicons (~4.1 kb) from *Blautia* MSK18.94 *cah* cluster in cDNA, no reverse transcriptase (No RT), genomic DNA (gDNA) and no template control (NTC) samples induced with (CG) or uninduced (H2O). A ~4.1 kb band represents the amplicon that covers the length of the *cah* cluster. **c** Genomic context of *H. hathewayi* and *E. clostridioformis* YL32, highlighting the mbile genetic elements and regulators interspereased with and around cah genes (green) and act genes (blue). (*) Indicates a gene annotated as a pseudogene.

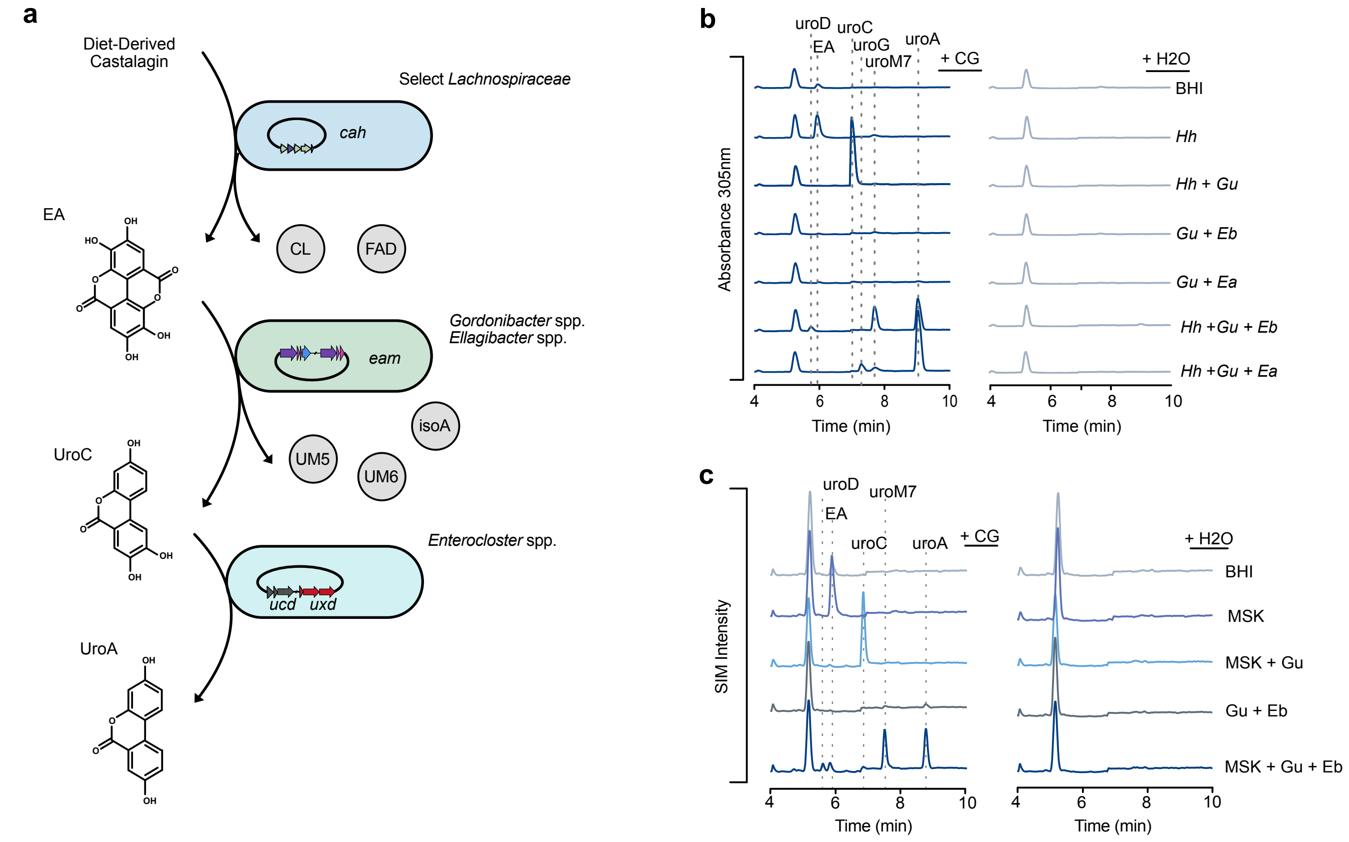


**Supplementary Figure 11.** Defined minimal communities dictate CG metabolism

**a** Schematic demonstrating the cross-feeding of castalagin metabolites. Select cah-harbouring *Lachnospiraceae* convert castalagin to ellagic acid (EA) and castalin (CL). eam-harbouring *Gordonibacter* spp. and *Ellagibacter* spp. metabolize ellagic acid to urolithins. Urolithin C can be further converted to terminal metabolite urolithin A by Enterocloster spp. via the *uxd* and *ucd* operons. **b** Aligned chromatograms ($\lambda = 305$ nm) of castalagin (CG) or vehicle (H2O) treated cultures of H. hathewayi (Hh), Hh + G. urolithinfaciens (Gu), Gu + E. bolteae (Eb), Gu + E. asparagiformis (Ea), Hh + Gu, + Eb and Hh + Gu, + Ea after 5 days. **c** Aligned chromatograms of castalagin (CG) or vehicle (H2O) treated cultures of Blautia MSK18.94 (MSK), MSK + Gu, Gu + Eb and MSK + Gu, + Eb after 5 days.

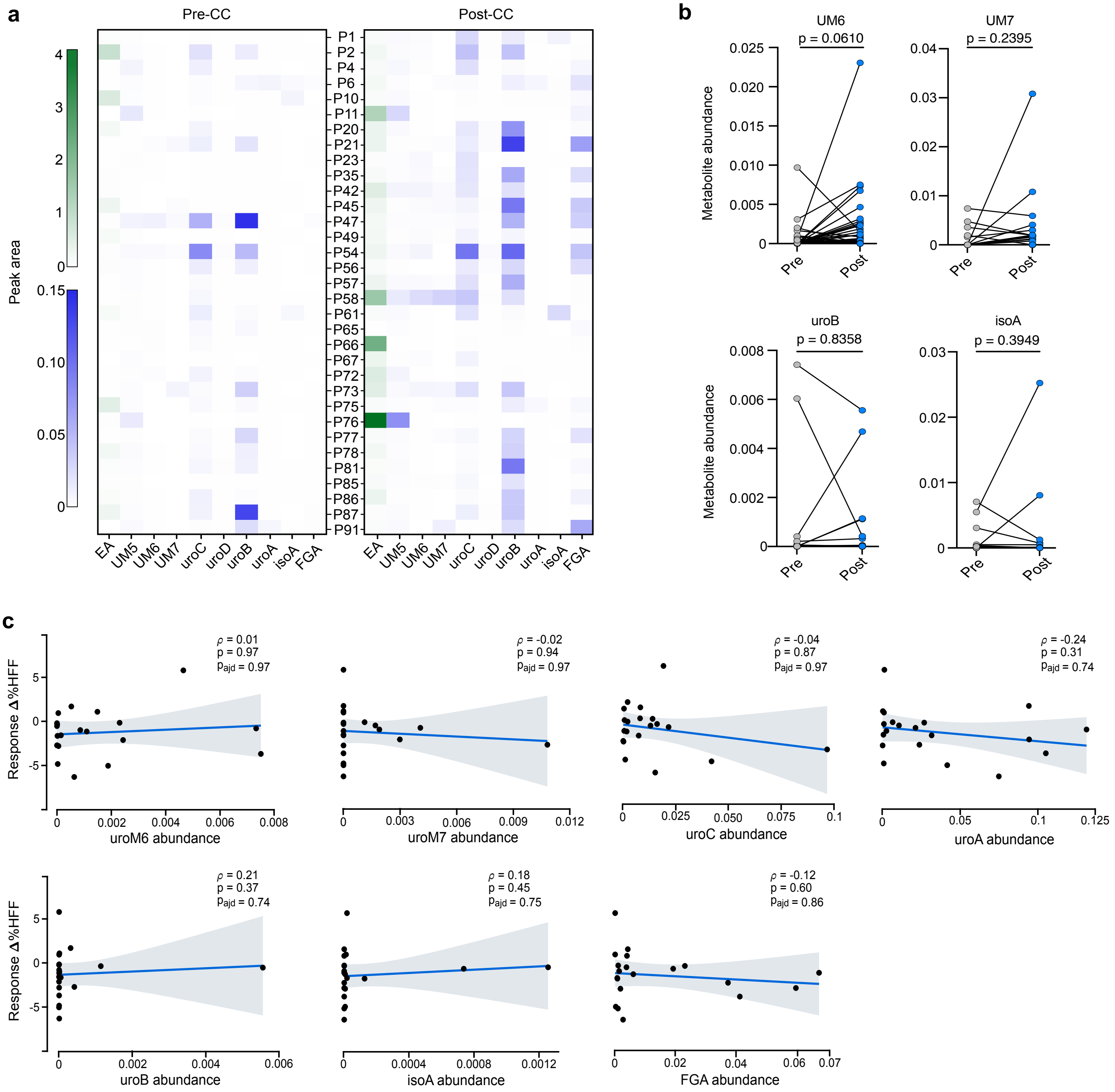


**Supplementary Figure 12.** Metabolite profiles vary among individuals in a patient cohort

**a** Heat map representing metabolite abundance (based on relative peak area) per patient pre- (left) and post- (right) camu camu (CC) supplementation. Ellagic acid (EA) is representated by the green scale. Urolithins and flavogallonic acid (FGA) are represented by the blue scale. All metbaolite abundances are normalized to their 100uM analytical standard. **b** Relative metabolite abundance for each patient before and after CC supplementation, for non-signficant changes in metabolites UM6, UM7, uroB and isoA. Analyses were performed using two-tailed paired wilcoxon signed rank test with Pratt's method and Holm-Sidak multiple comparison correction. **c** Correlation analysis between change in percent (%) hepatic fat fraction (HFF) and metabolite abundance for all patients (n = 20). Analyses performed using two-tailed Spearman rho correlation with Benjamini-Hochberg (BH) multiple comparison test. Spearman rho ($\rho$), p-values (p) and FDR (padj) values are indicated on each graph.

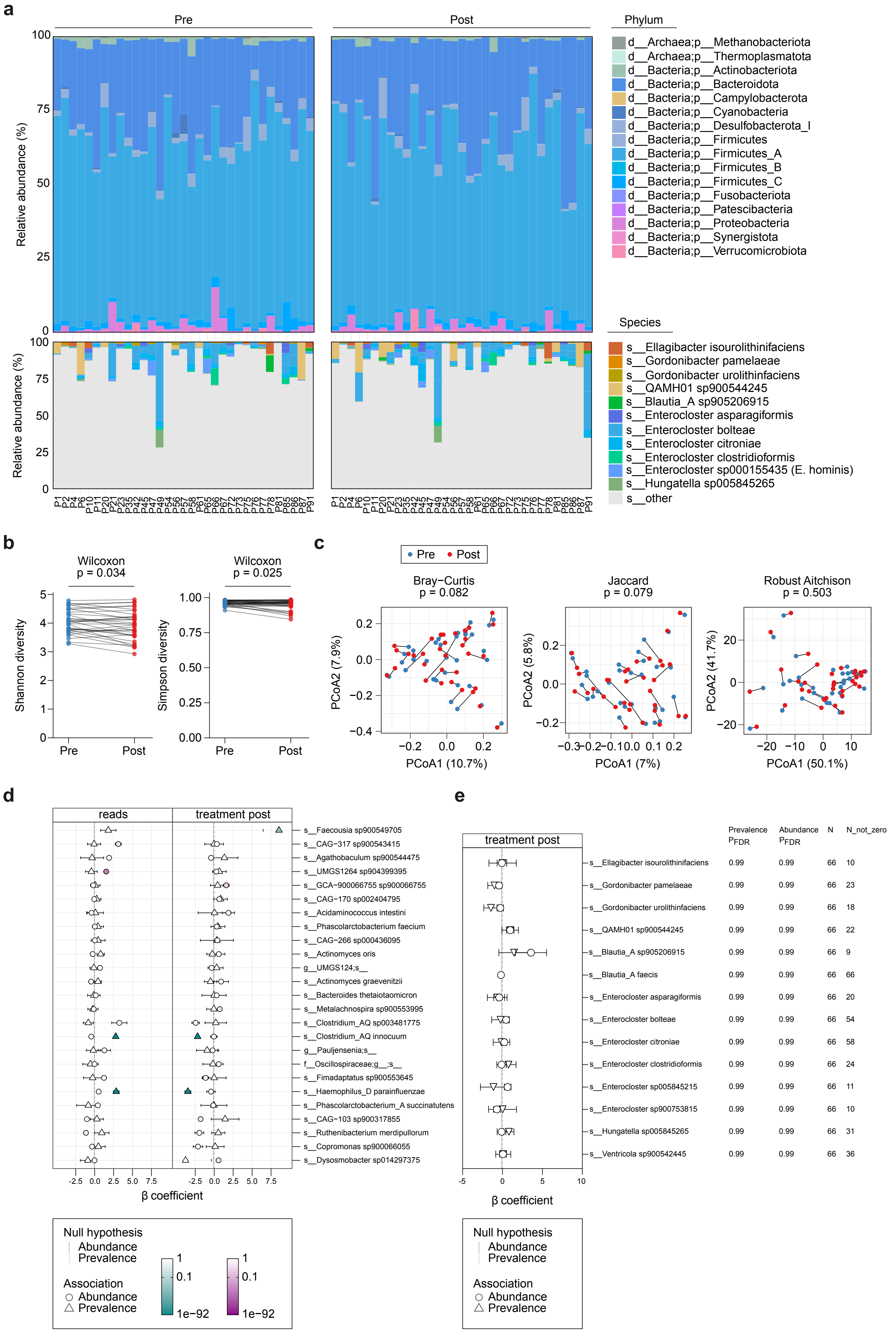
a
Pre
Post
Relative abundance (%)
Phylum
d__Archaea;p__Methanobacteriota
d__Archaea;p__Thermoplasmatota
d__Bacteria;p__Actinobacteriota
d__Bacteria;p__Bacteroidota
d__Bacteria;p__Campylobacterota
d__Bacteria;p__Cyanobacteria
d__Bacteria;p__Desulfobacterota_I
d__Bacteria;p__Firmicutes
d__Bacteria;p__Firmicutes_A
d__Bacteria;p__Firmicutes_B
d__Bacteria;p__Firmicutes_C
d__Bacteria;p__Fusobacteriota
d__Bacteria;p__Patescibacteria
d__Bacteria;p__Proteobacteria
d__Bacteria;p__Synergistota
d__Bacteria;p__Verrucomicrobiota
Species
s__Ellagibacter isourolithinifaciens
s__Gordonibacter pamelaeae
s__Gordonibacter urolithinfaciens
s__QAMH01 sp900544245
s__Blautia_A sp905206915
s__Enterocloster asparagiformis
s__Enterocloster bolteae
s__Enterocloster citroniae
s__Enterocloster clostridioformis
s__Enterocloster sp000155435 (E. hominis)
s__Hungatella sp005845265
s__other
b
Wilcoxon
p = 0.034
Wilcoxon
p = 0.025
Shannon diversity
Simpson diversity
Pre
Post
c
Pre
Post
Bray−Curtis
p = 0.082
Jaccard
p = 0.079
Robust Aitchison
p = 0.503
PCoA2 (7.9%)
PCoA1 (10.7%)
PCoA2 (5.8%)
PCoA1 (7%)
PCoA2 (41.7%)
PCoA1 (50.1%)
d
reads
treatment post
s__Faecousia sp900549705
s__CAG−317 sp900543415
s__Agathobaculum sp900544475
s__UMGS1264 sp904399395
s__GCA−900066755 sp900066755
s__CAG−170 sp002404795
s__Acidaminococcus intestini
s__Phascolarctobacterium faecium
s__CAG−266 sp000436095
s__Actinomyces oris
g__UMGS124;s__
s__Actinomyces graevenitzii
s__Bacteroides thetaiotaomicron
s__Metalachnospira sp900553995
s__Clostridium_AQ sp003481775
s__Clostridium_AQ innocuum
g__Pauljensenia;s__
f__Oscillospiraceae;g__;s__
s__Fimadaptatus sp900553645
s__Haemophilus_D parainfluenzae
s__Phascolarctobacterium_A succinatutens
s__CAG−103 sp900317855
s__Ruthenibacterium merdipullorum
s__Copromonas sp900066055
s__Dysosmobacter sp014297375
β coefficient
Null hypothesis
Abundance
Prevalence
Association
Abundance
Prevalence
1
0.1
1e−92
e
treatment post
Prevalence PFDR
Abundance PFDR
N
N_not_zero
s__Ellagibacter isourolithinifaciens 0.99 0.99 66 10
s__Gordonibacter pamelaeae 0.99 0.99 66 23
s__Gordonibacter urolithinfaciens 0.99 0.99 66 18
s__QAMH01 sp900544245 0.99 0.99 66 22
s__Blautia_A sp905206915 0.99 0.99 66 9
s__Blautia_A faecis 0.99 0.99 66 66
s__Enterocloster asparagiformis 0.99 0.99 66 20
s__Enterocloster bolteae 0.99 0.99 66 54
s__Enterocloster citroniae 0.99 0.99 66 58
s__Enterocloster clostridioformis 0.99 0.99 66 24
s__Enterocloster sp005845215 0.99 0.99 66 11
s__Enterocloster sp900753815 0.99 0.99 66 10
s__Hungatella sp005845265 0.99 0.99 66 31
s__Ventricola sp900542445 0.99 0.99 66 36
β coefficient
Null hypothesis
Abundance
Prevalence
Association
Abundance
Prevalence

**Supplementary Figure 13.** Microbiome composition is not altered in patient samples post CC supplementation

**a** Taxonomic composition of patient fecal samples at the phylum (top) and species (bottom) level, pre- (left) and post- (right) CC supplementation. Species in the bottom plot represent taxa of interest in the metabolism of castalagin to terminal urolithins. Bars represent the relative abundance (%) of taxa per patient. **b** Alpha diversity per patient pre- (blue dots) and post-CC (red dots) supplementation represented by Shannon (left) and Simpson (right) diversity indices. Data are represented as individual points matched by patient; Wilcoxon singed rank exact test (matching by patient). Statistical significance is denoted above the plot with P-values. **c** Beta diversity analyses per patient pre- (blue dots) and post-CC (red dots) supplementation. Principle coordinate analyses are represented by Bray-Curtis, Jaccard and Robust Aitchison distances; PERMANOVA with 999 permutations. **d** Default MaAsLin3 output of species-level differences in abundance and prevalence pre- and post-CC (treatment post), accounting for the total number of reads in each sample (as a covariate). **e** MaAsLin3 output of species-level differences for selected taxa of interest involved in the metabolism of castalagin to terminal urolithins.

a

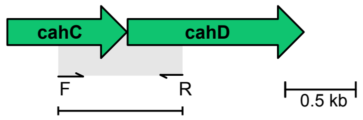


b

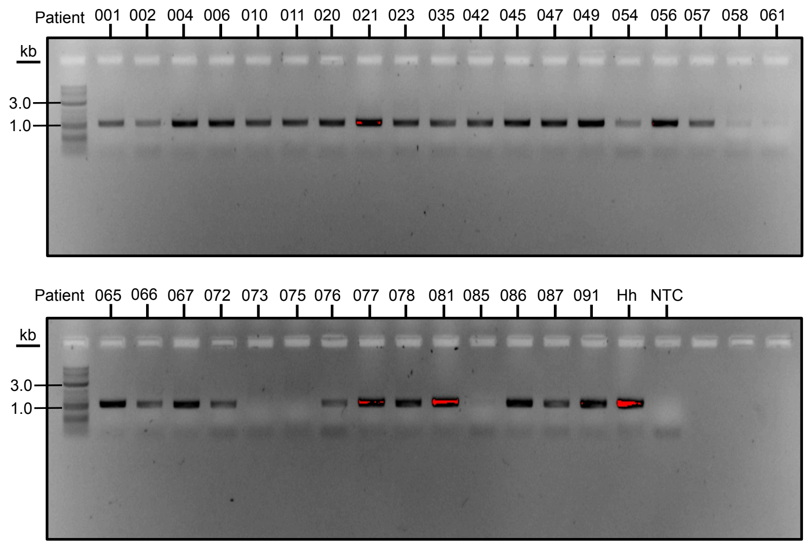


c

| Patient | Best hit | %ID | Alignment length |
|---|---|---|---|
| P10 | Hh | 99 | 1055 |
| P21 | Mf | 98.2 | 1088 |
| P23 | Hh | 100 | 1055 |
| P42 | Hh | 100 | 1046 |
| P47 | Hh | 99 | 1055 |
| P56 | Mf | 98.3 | 1088 |
| P66 | Hh | 99.2 | 1055 |
| P67 | Hh | 100 | 1055 |
| P87 | Hh | 100 | 1045 |

**Supplementary Figure 14.** cah genes are widely detected in patient fecal DNA samples

**a** Schematic of the *cah* specific primers targeting conserved nucleotide regions in the *cah*C and *cah*D genes of *Blautia* MSK18.94, *Mf* and *Hh*. **b** PCR products from the *cahC-D* amplification from stool genomic DNA, run on a 1% agarose gel. **c** Best match of sequenced amplicons to reference genes. Columns represent the patient sample, best match to the species reference amplicon, percent identity and amplicon alignment length. Hh: *H. hathewayi*, Mf: *M. formatexigens*.

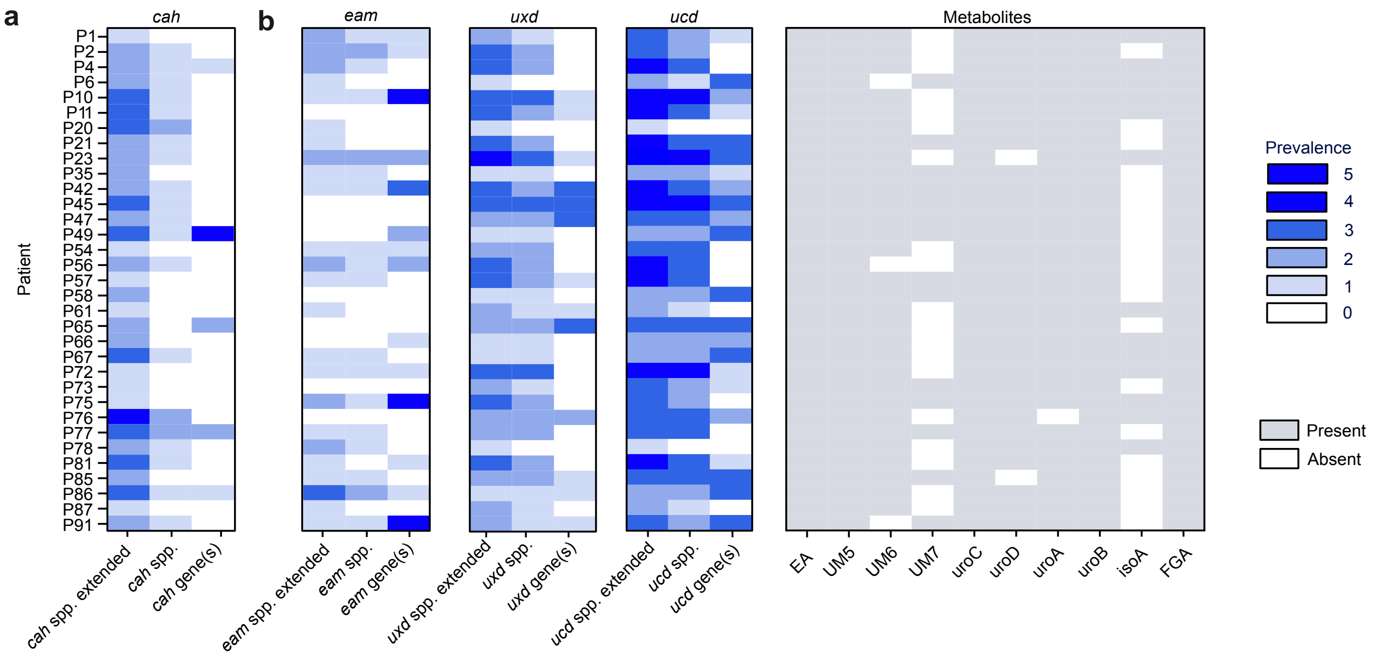


**Supplementary Figure 15.** *cah* homologs are detected in patient metagenomes

**a, b** Prevalence of species and genes in patient stool metagenomes post-CC treatment including (**a**) extended *cah* spp. including Blautia_A faecis and Enterocloster clostridioformis and (**b**) Prevalence of extended *eam*, *uxd* and *ucd* spp. including QAMH01 sp900544245 (*eam*), Ventricola sp900542445 (*uxd* and *ucd*). Extended spp. are an extension of species indicated in **Figure 4**. Presence (gray) or absence (white) of targeted metabolites ellagic acid (EA), urolithin M5 (uroM5), urolithin M6 (uroM6), urolithin M7 (uroM7), urolithin C (uroC), urolithin D (uroD), urolithin A (uroA), urolithin B (uroB), isourolithin A (isoA) and flavogallonic acid (FGA) are indicated per patient.

**a**

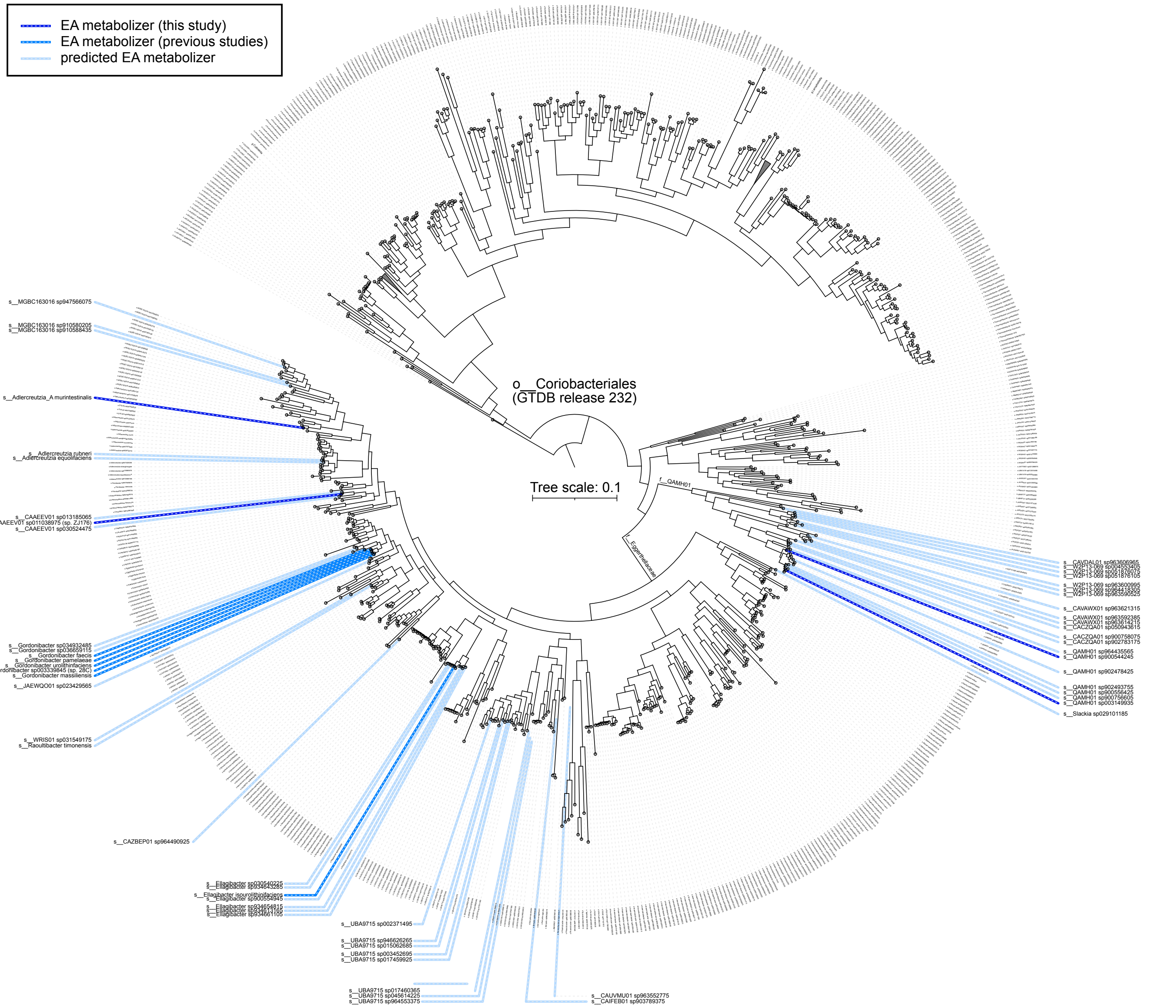


**b**

| GTDB species | Source/host(s) |
|---|---|
| s__MGBC163016 sp910580205 | mouse gut |
| s__MGBC163016 sp910588435 | mouse gut |
| s__Adlercreutzia_A murintestinalis | mouse gut |
| s__Adlercreutzia rubneri | human gut |
| s__Adlercreutzia equolifaciens | human gut |
| s__CAAEEV01 sp013185065 | marmot gut |
| s__CAAEEV01 sp011038975 (sp. ZJ176) | marmot gut |
| s__CAAEEV01 sp030524475 | owl monkey gut |
| s__Gordonibacter sp034932485 | bioreactor (groundwater) |
| s__Gordonibacter sp036659115 | salamander gut |
| s__Gordonibacter faecis | human gut, chicken gut |
| s__Gordonibacter pamelaeae | human gut, rat gut, mouse gut |
| s__Gordonibacter urolithinfaciens | human gut, chicken gut, cattle gut |
| s__Gordonibacter sp003339845 (sp. 28C) | human gut |
| s__Gordonibacter massiliensis | human gut |
| s__JAEWQO01 sp023429565 | cattle gut |
| s__WRIS01 sp031549175 | millipede gut |
| s__Raoultibacter timonensis | human gut |
| s__CAZBEP01 sp964490925 | ptarmigan gut |
| s__Ellagibacter sp030540225 | lemur gut |
| s__Ellagibacter sp934643285 | human gut |
| s__Ellagibacter isourolithinifaciens | human gut, mouse gut |
| s__Ellagibacter sp900554945 | human gut |
| s__Ellagibacter sp934654815 | human gut |
| s__Ellagibacter sp934613165 | human gut |
| s__Ellagibacter sp934661105 | human gut |
| s__UBA9715 sp002371495 | sheep gut |
| s__UBA9715 sp946626265 | sheep gut |
| s__UBA9715 sp015062685 | goat gut |
| s__UBA9715 sp003452695 | sheep gut |
| s__UBA9715 sp017459925 | deer gut |
| s__UBA9715 sp017460365 | deer gut |
| s__UBA9715 sp045614225 | cattle gut |
| s__UBA9715 sp964553375 | rat gut |
| s__CAIFEB01 sp903789375 | bioreactor (waste) |
| s__CAUVMU01 sp963552775 | human oral cavity |
| s__CAVDAL01 sp963606965 | human gut, chimpanzee gut |
| s__W2P13-069 sp004553405 | boar gut |
| s__W2P13-069 sp051876075 | boar gut |
| s__W2P13-069 sp051876105 | boar gut |
| s__W2P13-069 sp963600995 | human gut |
| s__W2P13-069 sp964418305 | unclear |
| s__W2P13-069 sp963590525 | human gut |
| s__CAVAWX01 sp963621315 | human gut |
| s__CAVAWX01 sp963592385 | chimpanzee gut |
| s__CAVAWX01 sp963614215 | human gut, bonobo gut |
| s__CACZQA01 sp050943615 | bioreactor (biogas) |
| s__CACZQA01 sp900758075 | human gut |
| s__CACZQA01 sp902783175 | human gut, pig gut |
| s__QAMH01 sp964435565 | bioreactor (biogas) |
| s__QAMH01 sp900544245 | human gut |
| s__QAMH01 sp902478425 | human gut |
| s__QAMH01 sp902493755 | human gut |
| s__QAMH01 sp900556425 | human gut |
| s__QAMH01 sp900756605 | human gut |
| s__QAMH01 sp003149935 | human gut |
| s__Slackia sp029101185 | pig gut |

**Supplementary Figure 16.** Genomes of cultured and uncultured Coriobacteriales encode eam cluster protein homologues.

**a** Phylogenetic tree of the *Coriobacteriales* order based on GTDB classification (release 232). Taxa of with >1 strain encoding Eah (hydrolase/decarboxylase) and ≥ 1 Eadh (dehydroxylase) with ≥ 70 %id to validated eam-encoded enzymes are annotated in shades of blue according to the level of EA metabolism validation. The QAMH01 and *Eggerthellaceae* families are represented by outer rings coloured in shades of grey. **b** Table of host(s) or sampling sources for taxa annotated in (**a**) based on NCBI metadata. Source data for (**a**) are provided as a Source data file.

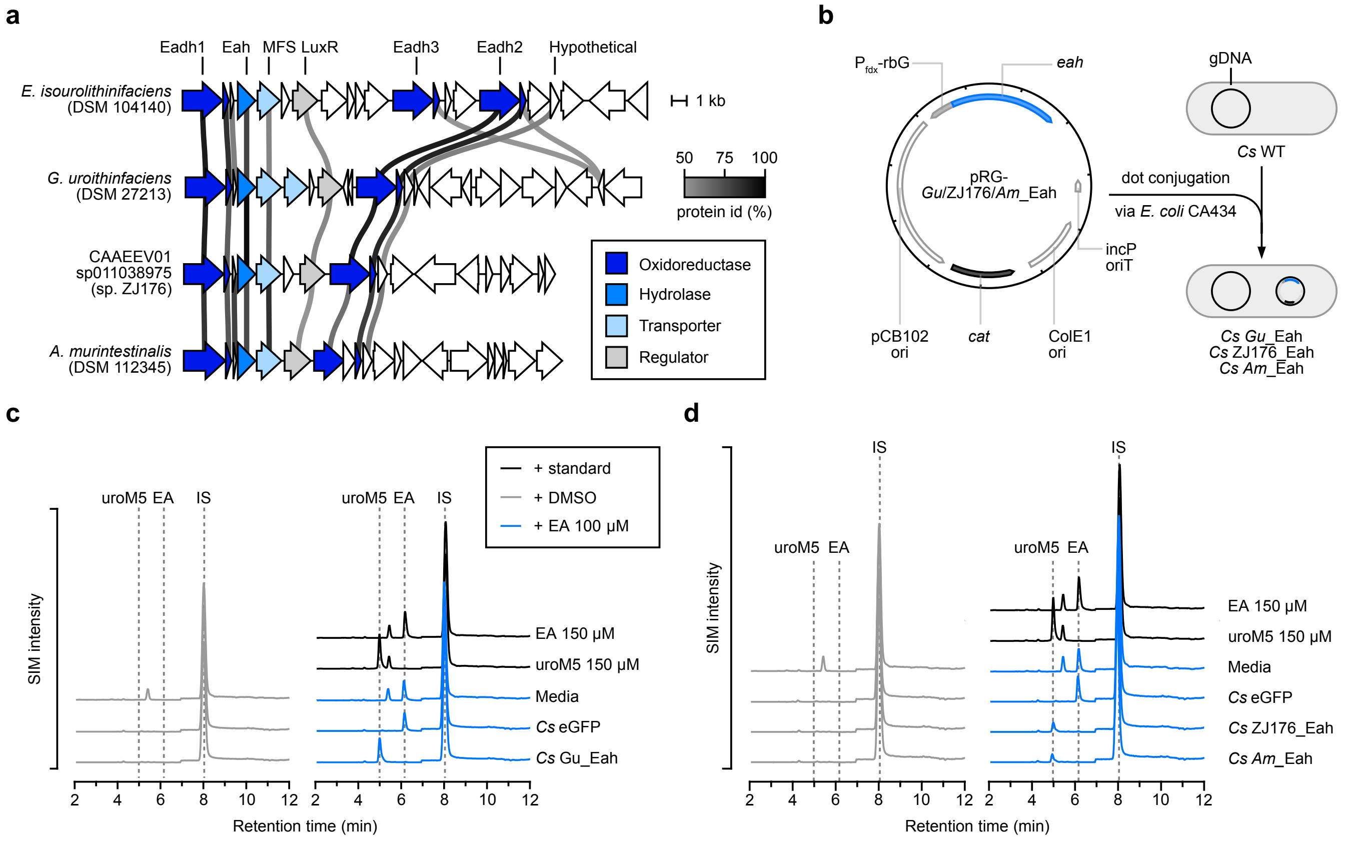


**Supplementary Figure 17.** Eah homologues from *Coriobacteriales* metabolize ellagic acid

**a** Genomic context of known EA metabolizers, *G. urolithinfaciens* and *E. isourolithinifaciens*, with annotated genomic regions of CAAEEV01 sp011038975 and *Adlercreutzia murintestinalis*. Alignments were performed using the clinker tool with genome accessions in **Supplementary Table 3**. Protein sequences with identities ≥50% are connected by lines colored according to the percent identity value. The strain identity for each contig are denoted below the contig number. **b** Schematic representation of the conjugation and heterologous expression workflow to validate Eah homologues in *C. sporogenes* (*Cs*) whole cell assays. The generic pRG plasmid map is labeled with the origin of replication (ori), chloramphenicol acetyltransferase (*cat*), and promoter of the *Cs* ferredoxin gene with riboswitch G (Pfdx-rbG). **c** Aligned selected ion monitoring (SIM) chromatograms of substrates and metabolites from mid-exponential phase *Cs* cultures treated with 100 µM of EA or DMSO. Tested conditions included sterile media or *Cs* transconjugants expressing enhanced green fluorescent protein (eGFP) or *G. urolithinfaciens* Eah (Gu_Eah). **d** Aligned SIM chromatograms following the same experimental design as (**c**) using *Cs* transconjugants expressing enhanced green fluorescent protein (eGFP),CAAEV01 sp011038975 sp. ZJ176 Eah (ZJ176_Eah), or A. murintestinalis Eah (*Am*_Eah). For **c**-**d**, retention times of EA and urolithin standards are shown with grey dotted lines. Data from n = 1 of 3 biological replicates for each transconjugant are shown.
